# PTPRF is a stress-responsive cytoskeletal checkpoint that coordinates metabolic adaptation in hepatocytes and β cells

**DOI:** 10.64898/2026.09.03.748858

**Authors:** Mayank Bansal, Wadsen St-Pierre-Wijckmans, Javier Negueruela, Carlos E. Buss, Francisco Ribeiro-Costa, Ao Li, Tiffany Lai, Vishnu M. Nair, Valerie Vandenbempt, Michela Lisjak, Israel Pérez-Chávez, Gabriel G. Hovhannisyan, Daria Ezeriņa, Mehmet Bostanciklioglu, Stéphane Demine, Lu Yu, David Oxley, Nizar Mourad, Sumeet Pal Singh, Joris Messens, Andrés Balaguer-Román, María D. Frutos, Carlos M. Martinez, Bruno Ramos-Molina, Valérie Suain, Louise Conrard, Nicolas Baeyens, Didier Vertommen, Alessandra K. Cardozo, Patrick Gilon, Hayley J. Sharpe, Eduardo H. Gilglioni, Esteban N. Gurzov

## Abstract

Cytoskeletal remodeling is essential for adaptation to nutrient availability, yet how cells coordinate actin dynamics with glucose homeostasis in metabolic organs remains unclear. Here, we identify a pathway linking metabolic stress to actin reorganization in hepatocytes and pancreatic β cells. This mechanism involves transcriptional repression of the receptor protein tyrosine phosphatase PTPRF by spliced XBP1, a key unfolded protein response factor. In hepatocytes, PTPRF loss under dietary stress enhances insulin signaling, increases mitochondrial respiration and reduces steatosis. Proteomic analyses show that PTPRF interacts with regulators of actin polymerization and cell junctions, and its deletion promotes actin filament organization, shifting metabolism toward oxidative pathways. In β cells, PTPRF deficiency similarly enhances actin polymerization and augments glucose-stimulated insulin secretion in obesity. Collectively, these findings place PTPRF as a nutrient-responsive regulator of cytoskeletal remodeling that coordinates hepatic metabolism and β-cell function, highlighting its potential as a therapeutic target for improving systemic glucose control.

**Highlights:**

- Actin cytoskeletal organization links cellular structure to metabolism through PTPRF
- PTPRF loss sustains insulin signaling and oxidative metabolism in hepatocytes
- PTPRF restrains actin-dependent insulin granule secretion in β cells
- Cytoskeletal plasticity promotes metabolic adaptation to chronic nutrient excess

## Introduction

Cell assembly into functional tissues depends on a cytoskeletal and adhesion network that maintains mechanical integrity while processing dynamic signal information from the environment. The actin cytoskeleton is a central node to sense neighboring cells, extracellular matrix stiffness, fluid shear and nutritional status [1–4]. Actin filaments also support structural scaffolding, vesicle trafficking, motility and cytokinesis, and these activities require continuous filament turnover with high, spatially restricted ATP demand [3]. As a result, cytoskeletal organization is tightly coupled to energy metabolism, where ATP production through glycolysis and oxidative phosphorylation is aligned with the energetic demand of actin polymerization under different conditions of nutrient availability [5]. When this alignment is maintained, localized ATP supply is sufficient to sustain cytoskeletal dynamics. If it fails, insufficient nutrient and ATP availability can impair cytoskeletal remodeling, disrupting cellular mechanics, energy homeostasis and tissue integrity, ultimately predisposing to pathological remodeling. Despite this fundamental interdependence between mechanics and metabolism, the molecular basis of this connection remains limited.

In obesity, nutrient excess and chronic inflammation perturb the mechano–metabolic equilibrium in the liver and pancreas. Lipid accumulation stiffens the extracellular matrix, imposing mechanical and oxidative stress on parenchymal cells [6, 7]. In hepatocytes, actin remodeling affects lipid droplet formation, mitochondrial dynamics, and insulin receptor signaling [8, 9]. In pancreatic β cells, cytoskeletal dynamics regulate vesicle trafficking and insulin secretion, directly linking mechanical properties to endocrine function [10]. Thus, the cytoskeleton operates both as a sensor and an effector of adaptation to nutrient overload in these two metabolically relevant cell types. At the plasma membrane, the actin-rich cortical cytoskeleton forms a dynamic barrier that influences receptor-proximal signaling and, in secretory cells, regulates access of insulin granules to the plasma membrane.

Protein tyrosine phosphatases (PTPs) translate extracellular and mechanical signals into intracellular responses by controlling tyrosine phosphorylation states [11, 12]. Previously thought to be simple signal terminators, PTPs are now considered modulators that fine-tune the phosphorylation status of signaling, affecting amplitude and duration. In metabolic tissues, PTPs regulate insulin and growth hormone pathways, thereby orchestrating systemic energy homeostasis [12]. In particular, receptor-type PTPs (RPTPs) link the extracellular environment and intracellular signaling through their dual-domain architecture, allowing mechanical and chemical sensing at the cell surface [13, 14].

We recently performed a systematic profiling of the hepatic phosphatome across metabolic dysfunction-associated steatotic liver disease (MASLD) progression and identified dynamic alterations in the expression of RPTPs [15]. While PTPRK, PTPRM, and PTPRE remained consistently elevated, hepatic PTPRF (also known as LAR) displayed a marked decrease throughout disease progression. PTPRF belongs to the R2A subfamily of RPTPs and regulates focal adhesions [16]. Earlier studies implicated PTPRF in insulin receptor regulation, cell adhesion, and neurite outgrowth [16–18]. PTPRF acts as a receptor for the fasting-induced hormone asprosin, that is secreted from white adipose tissue and promotes hepatic glucose production [19]. Notably, studies in invertebrates have revealed that Lar-family receptors, the closest phylogenetic orthologues of PTPRF, are not limited to adhesion, but also couple actin cytoskeleton remodeling to mitochondrial positioning and function [20]. Thus, Lar-family RPTPs can link cytoskeletal dynamics to bioenergetic control, suggesting that PTPRF may similarly influence mitochondrial localization and metabolic output in vertebrates.

Here, we identify PTPRF as a nutrient- and stress-responsive phosphatase that couples cytoskeletal remodeling to metabolic control in hepatocytes and pancreatic β cells, relevant to obesity. Collectively, our findings define PTPRF as a central integrator of cellular actin cytoskeleton organization with glycolysis, mitochondrial function and insulin signaling in obesity-associated liver and endocrine pancreas dysfunction.

## Results

### Hepatic PTPRF expression is decreased in advanced stages of MASLD

We hypothesized that PTPRF expression in hepatocytes during liver disease is stage-specific and analyzed PTPRF mRNA in transcriptomic datasets [21, 22]. RNA-seq deconvolution (**Supplementary Fig. 1a-d**) showed that the hepatocyte fraction drives bulk PTPRF levels in the liver, with a disease-associated decline attenuated after composition adjustment (**Fig. 1a**). Direct quantification of 53,225 hepatocyte nuclei from an integrated snRNA-seq atlas [22], supported a hepatocyte-intrinsic decrease of PTPRF (**Fig. 1b**, **c**). Immunohistochemistry (IHC) of human liver samples graded for steatosis (scores 0-3) showed a progressive loss of PTPRF protein with increasing steatosis severity (**Fig. 1d**; **Supplementary Fig. 1e**). To model MASLD progression, C57BL/6N mice were fed a high-fat (HFD), a high-fat/high-fructose/high- cholesterol (HFHFHCD), a high-fat/methionine–choline-deficient (HFMCD), or a control (CD) diet, for either 12 or 24 weeks (**Fig. 1e**). EchoMRI confirmed increased hepatic fat content in all 3 diets compared to controls (**Fig. 1e**). Western blot analysis showed a modest reduction in PTPRF expression after 12 weeks of HFD feeding, which became significant at 24 weeks (**Fig. 1f**). In HFHFHCD-fed mice, PTPRF was already markedly reduced at 12 weeks and remained low at 24 weeks (**Fig. 1g**). HFMCD-fed mice displayed an almost complete loss of PTPRF expression at 12 weeks (**Fig. 1h**). IHC analysis confirmed the downregulation of hepatic PTPRF in steatogenic diets compared to lean mice (**Fig. 1i**; **Supplementary Fig. 1f**). Collectively, these data demonstrate that in human MASLD, hepatic PTPRF expression declines as disease severity increases, and that this progressive decrease is recapitulated in mice exposed to diets with high-fat content.

**Figure 1.**
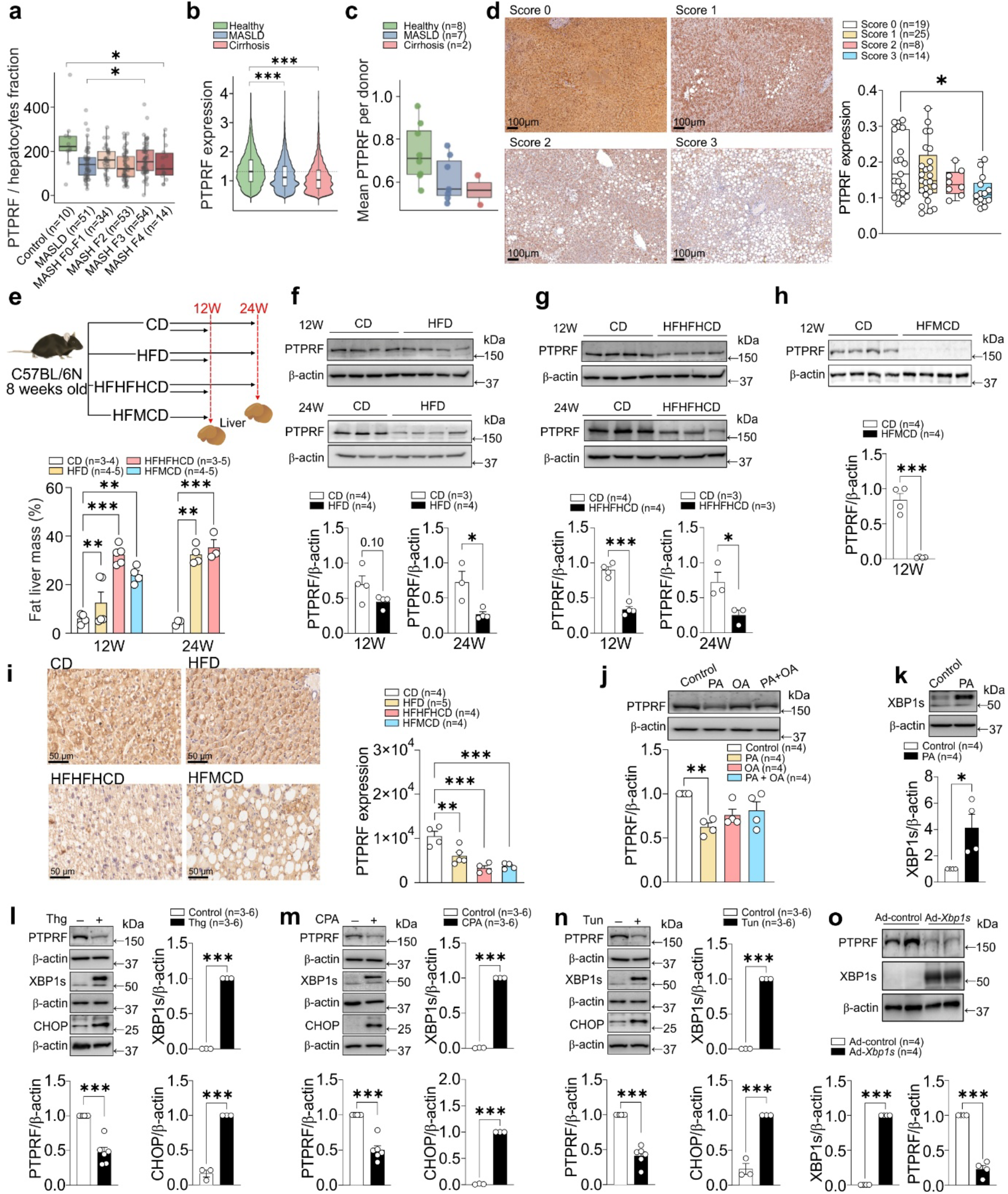
PTPRF expression is decreased in human and murine models of steatosis and in hepatocytes under ER stress. (**a**) Composition-adjusted PTPRF expression across healthy, MASH, and MASLD after correcting for differences in cellular composition in the dataset GSE135251. (**b**) Single-nucleus RNA-seq quantification of PTPRF expression in 53,225 hepatocyte nuclei from human liver samples. (**c**) Per-donor mean hepatocyte PTPRF expression across disease groups. (**d**) Immunohistochemical detection of PTPRF in human liver samples from individuals with steatosis scores ranging from 0 to 3; right, quantification of PTPRF- positive area. Scale bar: 100 μm. (**e**) Experimental design of murine dietary models of steatosis: control, HFD, HFHFHCD, and HFMCD, treated for 12 weeks (all diets) or 24 weeks (except HFMCD), and hepatic fat content (% of liver mass) determined by EchoMRI after diet interventions. (**f–h**) Western blot analysis of PTPRF protein levels in livers of mice fed HFD vs Control (f), HFHFHCD vs Control (g), and HFMCD vs Control (h). (**i**) Immunohistochemistry of PTPRF in liver sections from 12-week dietary models with quantification of staining intensity. Scale bar: 50 μm. (**j**) Western blot analysis of PTPRF expression in primary mouse hepatocytes (mHep) treated for 18 h with free fatty acids: palmitic acid (PA), oleic acid (OA), or a PA/OA mix. (**k**) XBP1s protein expression following 8-h treatment with PA in mHep; representative blot and quantification. (**l–n**) Western blot analysis of PTPRF, XBP1s, and CHOP in mHep treated with endoplasmic reticulum stressors: (**l**) 24 h Thapsigargin (Thg), (**m**) 6 h Cyclopiazonic acid (CPA), and (**n**) 6 h Tunicamycin (Tun). (**o**) Western blot showing PTPRF and XBP1s protein levels in mHep transduced with Ad-*Xbp*1s or Ad-Control. Bar graphs display quantification as mean ± SEM with individual data points. Statistical significance was determined using an unpaired t test (panels f, g, h, k, l, m, n, and o), one-way ANOVA (panels d, i, and j), or Kruskal–Wallis test followed by pairwise Wilcoxon rank-sum tests (panels a and b). \**P* < 0.05, \*\**P* < 0.01, \*\*\**P* < 0.001.

Primary mouse hepatocytes (mHep) exposed to palmitate saturated fatty acids had significantly reduced PTPRF levels (**Fig. 1j**). By contrast, inflammatory cytokines involved in MASLD (i.e., IFNγ, IL-6, TNFα, or TGF-β) did not change PTPRF levels (**Supplementary Figs. 1g-j**). Saturated free fatty acid exposure induces the unfolded protein response (UPR) [23]. We confirmed increased spliced X-box binding protein 1 (XBP1s) expression after palmitate treatment (**Fig. 1k**). Promoter motif analysis revealed high-confidence XBP1s-binding sites upstream of the PTPRF transcription start site, in both human and mouse gene, suggesting XBP1s can mediate PTPRF suppression **(Supplementary Fig. 1k)**. Analysis of XBP1s ChIP– seq datasets identified 26 XBP1s binding sites across the *Ptprf* locus **(Supplementary Fig. 1k)**. A subset of peaks contained canonical endoplasmic reticulum (ER) stress response elements (ERSE) **(Supplementary Fig. 1l)**, consistent with direct XBP1–DNA binding. Motif enrichment analysis of non-canonical peaks identified NF-Y (CCAAT box) as the predominant co-factor motif, together with signatures of additional UPR-associated transcription factors (**Supplementary Fig. 1m**). Furthermore, chemical induction of the UPR in mHep using thapsigargin, cyclopiazonic acid, or tunicamycin led to activation of XBP1s and CHOP, accompanied by consistent PTPRF downregulation (**Fig. 1l-n**). Finally, adeno-mediated expression of XBP1s in mHep reduced both *Ptprf* mRNA (**Supplementary Fig. 1n**) and PTPRF protein levels (**Fig. 1o**), suggesting direct gene regulation. Together, these findings demonstrate that PTPRF downregulation is associated with the UPR in hepatocytes under nutrient overload and are consistent with XBP1s-mediated transcriptional repression.

### Loss of hepatic PTPRF improves insulin sensitivity and attenuates diet-induced steatosis

To investigate the functional consequences of PTPRF loss in experimental MASLD, we generated a hepatocyte-specific *Ptprf* knockout mouse (*Ptprf^ΔHep^*). Targeted deletion of *Ptprf* exons 8–10 was confirmed by qRT-PCR in mHep from *Ptprf^ΔHep^* mice (**Fig. 2a**). Consistent with the implication of PTPRF in regulation of the insulin receptor [17], *Ptprf^ΔHep^* mHep displayed enhanced phosphorylation of the insulin receptor at the initial insulin stimulation and AKT after insulin stimulation compared with controls (**Fig. 2b**). Metabolic characterization showed comparable body weights, fat-lean body mass, and liver weight and fat content between *Ptprf^ΔHep^* and control mice in chow diet (**Fig. 2d**, **e**; **Supplementary Fig. 2a-c**). However, when mice were challenged with HFHFHCD for 12 weeks, *Ptprf^ΔHep^*mice gained less body weight than control littermates (**Fig. 2c**). Body composition analysis showed a pronounced increase in fat mass in control mice, whereas in *Ptprf^ΔHep^* littermates fat mass expansion was reduced (**Fig. 2d**). Following HFHFHCD feeding, control mice showed a marked increase in liver weight and fat, whereas *Ptprf^ΔHep^* mice displayed only a modest non-significant increase (**Fig. 2e**; **Supplementary Fig. 2b**, **c**). Histological analysis corroborated these findings as *Ptprf^ΔHep^* livers exhibited preserved tissue architecture with reduced signs of lipid deposition (**Fig. 2f**). Indirect calorimetry showed no differences in whole-body energy expenditure, oxygen consumption, respiratory exchange ratio, ambulatory activity, or food and water intake between control and *Ptprf^ΔHep^* mice upon HFHFHCD feeding (**Supplementary Fig. 2d-i**).

**Figure 2.**
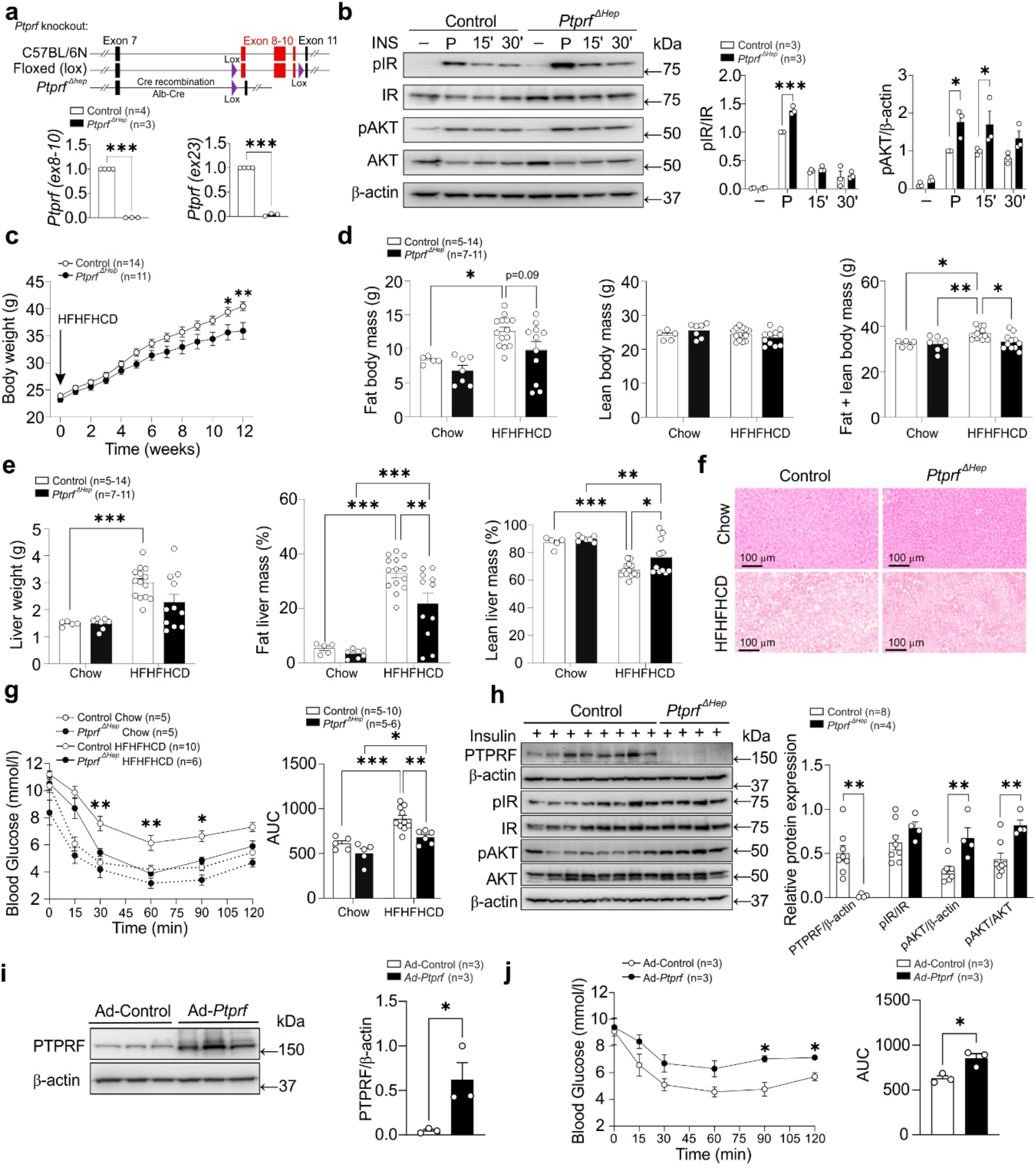
Hepatocyte-specific deletion of PTPRF improves insulin signaling and metabolic outcomes under HFHFHCD feeding. (**a**) Schematic of the strategy to generate hepatocyte- specific Ptprf knockout (*Ptprf^ΔHep^*) mice using Albumin-Cre-mediated deletion of exons 8–10. (**b**) Western blot analysis and quantification of insulin signaling in primary mouse hepatocytes (mHep) from *Ptprf^ΔHep^* and littermate control mice following insulin stimulation (100 nM) with a 10-min pulse (P), wash and chase at 15- and 30-min. Blots probed for phosphorylated and total insulin receptor (IR) and AKT. (**c**) Body weight changes in male *Ptprf^ΔHep^* and littermate control mice after 12 weeks on HFHFHCD. (**d**) Fat mass and lean mass (EchoMRI) in *Ptprf^ΔHep^* and control mice after 12 weeks on HFHFHCD and in age-matched mice on chow diet. (**e**) Liver weight and fat and lean masses in *Ptprf^ΔHep^* and control mice after 12 weeks on HFHFHCD and in age-matched mice on chow diet. (**f**) Hematoxylin and eosin (H&E) staining of liver sections from *Ptprf^ΔHep^* and control mice on chow or HFHFHCD. Scale bar: 100 μm. (**g**) Insulin tolerance test (ITT) in *Ptprf^ΔHep^* and control mice after 12 weeks of HFHFHCD, with area-under-the-curve (AUC) quantification. (**h**) Western blot analysis of liver lysates from *Ptprf^ΔHep^* and control mice collected 10 min after insulin injection (0.75 U/kg, i.p.) following 12 weeks of HFHFHCD. Blots include PTPRF, pIR, IR, pAKT, and AKT. (**i**) Western blot of PTPRF expression and quantification in mice injected with Ad-*Ptprf* or Ad-Control assessed six weeks post-injection. (**j**) ITT in mice injected with Ad-*Ptprf* or Ad-Control, assessed six weeks post-injection. Bar graphs display quantification as mean ± SEM with individual data points. Statistical significance was determined using an unpaired t test (panels a, h and i) or Two-way ANOVA (panels b, c, d, e, g and j). \**P* < 0.05, \*\**P* < 0.01, \*\*\**P* < 0.001.

Insulin tolerance tests (ITT) revealed significantly improved whole-body insulin responsiveness in *Ptprf^ΔHep^* mice after HFHFHCD feeding, with blood glucose levels comparable to those of chow-fed mice (**Fig. 2g**). Consistently, liver lysates from HFHFHCD- fed mice collected after acute intraperitoneal insulin injection had elevated phosphorylation of insulin receptor and AKT in *Ptprf^ΔHep^* group compared with controls (**Fig 2h**). Loss-of-function studies were complemented by hepatic overexpression of PTPRF using *in vivo* adenoviral delivery. Efficacy in hepatocytes was confirmed *in vitro* (**Supplementary Fig. 2j**) and *in vivo* (**Fig. 2i)**. Rescue of PTPRF in *Ptprf^ΔHep^*hepatocytes reduced phosphorylation of AKT (**Supplementary Fig. 2k**). *In vivo*, adenoviral PTPRF overexpression impaired insulin sensitivity in wild-type mice, as assessed by ITT (**Fig. 2j**). Together, these data indicate that hepatic PTPRF restrains fat accumulation and insulin signaling in the context of diet-induced obesity.

### PTPRF links actin cytoskeleton organization to lipid storage and glycolytic flux in hepatocytes

Having established that hepatic PTPRF is responsive to lipid overload, limits insulin signaling in obesity, and influences diet-induced steatosis in mice, we examined its subcellular localization in hepatocytes to identify where it engages with signaling complexes. Live-cell confocal microscopy revealed that GFP-tagged full-length PTPRF localized at the plasma membrane (**Fig. 3a**), particularly at cell–cell contact zones, indicating its enrichment at intercellular interfaces. GFP signal was also observed in intracellular vesicle-like structures, consistent with trafficking between the membrane and endosomal compartments. The result was confirmed by immunofluorescence staining of endogenous PTPRF expression showing prominent membrane localization (**Fig. 3a**).

**Figure 3.**
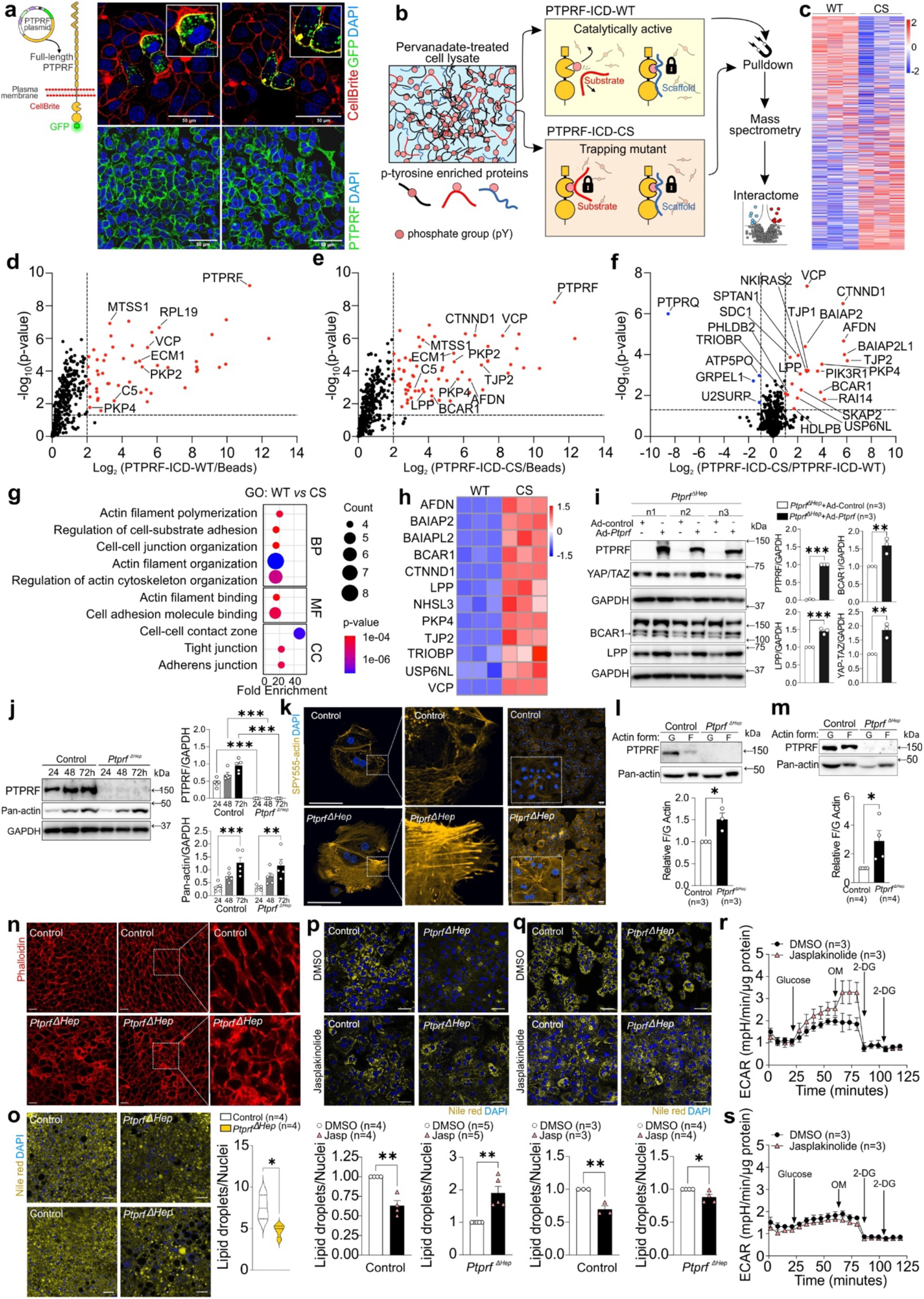
Full-length PTPRF localizes to the plasma membrane and regulates the actin cytoskeleton. **(a)** Representative fluorescence images of HepG2 cells overexpressing green fluorescent protein (GFP)-tagged full-length PTPRF, with plasma membrane staining to assess subcellular localization (top images) or HepG2 cells immunofluorescence staining of endogenous PTPRF (bottom). **(b)** Schematic representation of the interactome profiling strategy using catalytically active wildtype (WT) PTPRF intracellular domain (ICD) or a catalytically inactive cysteine-to-serine (CS) trapping mutant. **(c)** Heatmap showing differential interactors between WT and CS PTPRF ICD in primary mouse hepatocytes (mHep). **(d–f)** Volcano plots depicting differentially enriched interactors in WT, CS and beads conditions in mHep. **(g)** Pathway enrichment analysis of significantly enriched PTPRF ICD interactors in mHep. (**h**) Heatmap showing the relative enrichment of the 12 shared interacting proteins under WT and CS PTPRF ICD conditions in mHep and HepG2 cells. **(i)** Immunoblot analysis of mHep from *Ptprf^ΔHep^* mice transduced with Ad-*Ptprf* or Ad-control. **(j)** Immunoblot analysis of mHep from *Ptprf^ΔHep^* and control mice collected at the indicated time points. **(k)** Staining of F- actin in mHep from *Ptprf^ΔHep^* and control mice to assess cytoskeletal organization. Scale bar: 50 μm. **(l)** Immunoblot analysis of F actin and G actin fractions in mHep from *Ptprf^ΔHep^* and control mice. **(m)** Immunoblot analysis of F actin and G actin fractions from liver tissue of *Ptprf^ΔHep^* and control mice following 12 weeks of HFHFHCD feeding. **(n)** Phalloidin staining of liver sections from *Ptprf^ΔHep^* and control mice after 6 weeks of HFHFHCD feeding. Scale bar: 50 μm. **(o)** Nile Red staining of liver sections from *Ptprf^ΔHep^* and control mice after 6 weeks of HFHFHCD feeding. Scale bar: 50 μm. **(p, q)** Representative fluorescence images and quantification showing lipid droplet accumulation (yellow, Nile Red) and nuclei (blue, DAPI) in *Ptprf^ΔHep^* and control mHep treated with jasplakinolide. Steatosis was induced by glucose and insulin (p; 48 h) or by free fatty acids (q; 24 h; 400 μM palmitic acid and 800 μM oleic acid). Scale bar: 50 μm. **(r)** Glycolysis stress test in control mHep treated with jasplakinolide or DMSO. **(s)** Glycolysis stress test in *Ptprf^ΔHep^*mHep treated with jasplakinolide or DMSO. Bar graphs display quantification as mean ± SEM with individual data points. Statistical significance was determined using an unpaired t test (panels i, l, m, o, p and q) or One-way ANOVA (panel j). \**P* < 0.05, \*\**P* < 0.01, \*\*\**P* < 0.001.

To identify proteins associating with the intracellular domain of PTPRF in hepatocytes, affinity purification was performed as schematized in **Fig. 3b**. mHep and human HepG2 cells were treated with pervanadate to induce tyrosine phosphorylation (**Supplementary Fig. 3a**). Recombinant intracellular domains (ICD) of wild-type (WT) and substrate trapping catalytic- dead (CS) recombinant PTPRF were assessed for phosphatase activity (**Supplementary Fig. 3b**) and used to identify scaffold and catalytic substrates in pulldowns. Bound proteins were digested and identified by mass spectrometry in mHep (**Fig. 3c**) and HepG2 (**Supplementary Fig. 3c**) lysates. In mHep lysates, 52 proteins were significantly enriched in the WT PTPRF- ICD pulldown, including PKP2, VCP, and ECM1 compared to a beads-only control (**Fig. 3d**). The CS domain enriched 66 proteins, including CTNND1, PKP2, TJP2, and AFDN, that are components of junctional complexes and cytoskeletal architecture (**Fig. 3e**). Differential analysis identified 22 CS-selective and 4 WT-selective interactors (**Fig. 3f**). In HepG2 lysates, 44 proteins were significantly enriched with WT PTPRF-ICD, whereas 79 proteins were enriched with the CS domain, including PKP2/3/4, AFDN, WASF2, VCP, and IRS1 (**Supplementary Fig. 3d-f**). Gene ontology (GO) analysis indicated enrichment of pathways related to actin filament organization, actin polymerization dynamics, cell junction assembly, and adhesion binding in both mouse and human hepatocytes (**Fig. 3g**; **Supplementary Fig. 3g**). Cross-species comparison revealed conservation of shared PTPRF-interacting proteins with the CS variant (**Fig. 3h; Supplementary Fig. 3h**). Overlapping GO pathways between mHep and HepG2 cells showed common enrichment of actin cytoskeleton and cell junction proteins (**Supplementary Fig. 3i**, **j**). Consistently, overexpression of PTPRF in *Ptprf^ΔHep^*hepatocytes increased cytoskeleton/related proteins BCAR1, LPP and YAP/TAZ (**Fig 3i**). Co- immunoprecipitation assays demonstrated that both BCAR1 and LPP interact with PTPRF (**Supplementary Fig. 3m**). Collectively, these results establish PTPRF as a scaffold for actin cytoskeleton regulators and cell adhesion components in both mouse and human hepatocytes.

To study cytoskeleton assembly, we assessed extended culture of mHep and observed a gradual increase in pan-actin levels, which was not affected by *Ptprf* deletion (**Fig. 3j**). By contrast, SPY555-actin staining of mHep isolated from mice fed HFHFHCD revealed more prominent distribution of F-actin in *Ptprf^ΔHep^* mHep (**Fig. 3k**) with abundant, long and linear stress fibers, when compared to steatotic hepatocytes from control mice. Quantification of SPY555-actin staining further showed increased total and cortical F-actin intensity in *Ptprf*^ΔHep^ hepatocytes relative to controls (**Supplementary Fig. 3k**). Consistently, acute re-expression of PTPRF in *Ptprf*^ΔHep^ mHep reduced F-actin levels (**Supplementary Fig. 3l**). *Ptprf^ΔHep^*mHep also had increased F/G-actin ratio compared with control cells (**Fig. 3l**), which was confirmed in liver samples from mice fed a HFHFHCD (**Fig. 3m**). F-actin staining with phalloidin revealed a thicker cellular cortical actin (**Fig. 3n**) in liver section from *Ptprf^ΔHep^* mice when compared with controls. Taken together, our results suggest that PTPRF restrains actin polymerization in hepatocytes.

Nile Red staining revealed reduced lipid droplet accumulation in *Ptprf^ΔHep^*livers (**Fig. 3o**). To determine if F-actin dynamics influence lipid handling in hepatocytes, cells were treated with jasplakinolide, which promotes actin polymerization by increasing F/G-actin ratio (**Supplementary Fig. 3n**). Jasplakinolide upregulated PTPRF in mHep, indicating that its expression responds to actin cytoskeleton remodeling (**Supplementary Fig. 3o**). When assessing lipid droplet formation, *Ptprf^ΔHep^* hepatocytes consistently accumulated less lipid than control cells when stimulated with either glucose plus insulin or with fatty acids (**Fig. 3p**, **q**). In control hepatocytes, stabilizing F-actin with jasplakinolide reduced lipid accumulation under both conditions, indicating that increased F-actin restrains lipid deposition (**Fig. 3p, q**). In *Ptprf^ΔHep^*mHep, F-actin stabilization still reduced fatty acid-induced lipid accumulation; however, glucose-driven lipid accumulation increased, suggesting that in the absence of PTPRF, glucose-driven lipogenesis is less sensitive to F-actin dynamics.

In the liver, glycolysis is tightly linked to stress fiber integrity [24]. Consistently, jasplakinolide- induced cytoskeletal stabilization increases the extracellular acidification rate (ECAR) in control hepatocytes compared with untreated cells (**Fig. 3r**; **Supplementary Fig 3p**). However, no changes in ECAR were observed in *Ptprf^ΔHep^* mHep after F-actin stabilization (**Fig. 3s**, **Supplementary Fig. 3q**). These results indicate that F-actin abundance enhances glycolysis, but this effect is uncoupled upon PTPRF deficiency.

### PTPRF drives coordinated remodeling of glycolysis, mitochondrial respiration, and F-actin networks in hepatocytes

To directly test whether PTPRF controls glycolysis in hepatocytes, we examined the effects of PTPRF rescue in *Ptprf^ΔHep^*cells. ECAR measurements revealed that gain-of-function PTPRF enhanced the glycolytic response to glucose in *Ptprf^ΔHep^* cells (**Fig. 4a**). Inhibition of mitochondrial ATP production further amplified this effect, increasing maximal glycolytic capacity and glycolytic reserve (**Fig. 4a**). The results were confirmed by using the HYlight biosensor for fructose 1,6-bisphosphate [25]. Under glucose stimulation, *Ptprf^ΔHep^*mHep re- expressing PTPRF accumulated more fructose 1,6-bisphosphate than control (**Fig 4b**). We examined mitochondrial function in mHep upon PTPRF deficiency and rescue (**Fig. 4c**). *Ptprf^ΔHep^* mHep displayed the greatest basal and maximal oxygen consumption, while rescuing PTPRF expression reduced spare respiratory capacity and maximal respiration (**Fig. 4d**). Our results indicate that PTPRF shifts hepatocyte metabolism toward glycolysis at the expense of mitochondrial oxidative capacity, whereas PTPRF loss maintains an oxidative, mitochondria-driven metabolism.

**Figure 4.**
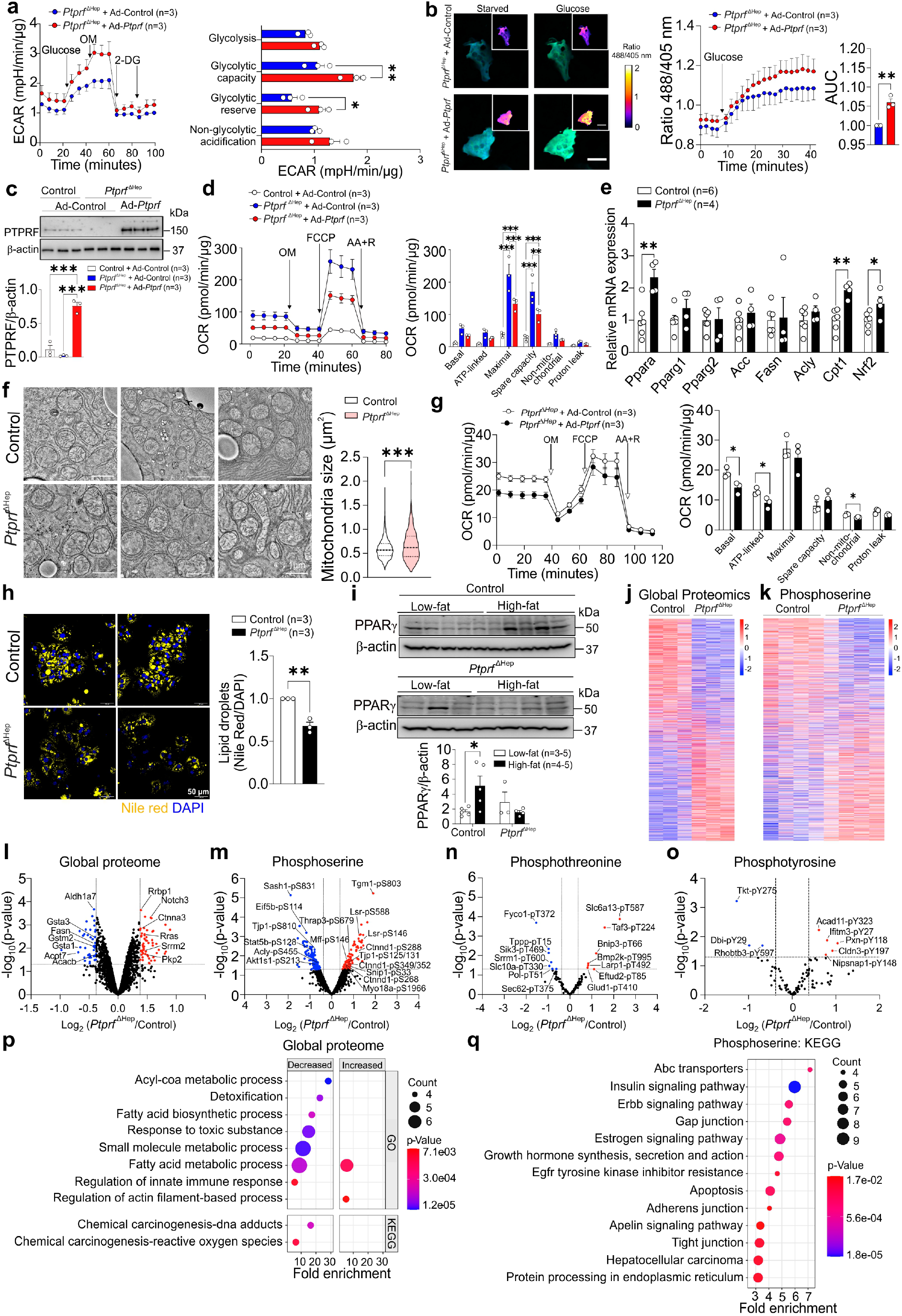
PTPRF regulates lipid oxidation and mitochondrial function in steatotic liver. **(a)** Glycolysis stress test performed in *Ptprf^ΔHep^* primary mouse hepatocytes (mHep) transduced with Ad-Control or Ad-*Ptprf***. (b)** HYlight mediated monitoring of fructose 1,6-bisphosphate dynamics in *Ptprf^ΔHep^*mHep transduced with Ad-*Ptprf* or Ad-Control. Scale bar: 1 μm. (**c**) Western blot of *Ptprf^ΔHep^* mHep infected with Ad-*Ptprf* or Ad-Control, and control wild-type mHep. **(d)** Oxygen consumption rate (OCR) trace from Seahorse mitochondrial stress test of *Ptprf^ΔHep^* mHep transduced with Ad-*Ptprf* or Ad-Control, and control mHep. Quantification of mitochondrial parameters is shown. **(e)** Relative mRNA levels of lipid metabolism genes in *Ptprf^ΔHep^* and control livers after 12 weeks of HFHFHCD feeding, assessed by qPCR. (**f**) Electron microscopy images of liver mitochondria from *Ptprf^ΔHep^* and control mice after 12 weeks of HFHFHCD, with corresponding quantification of mitochondrial size. Scale bar: 1 μm. **(g)** OCR analysis of *Ptprf*^ΔHep^ mHep transduced with Ad-*Ptprf* or Ad-Control under palmitate– supported respiration. **(h)** Nile Red and DAPI staining of high-fat mHep isolated from *Ptprf^ΔHep^*or control mice and quantification of lipid droplets. Scale bars, 50 μm. **(i)** Western blot analysis of PPARγ expression in low-fat and high-fat mHep from *Ptprf^ΔHep^* or control livers. **(j)** Heatmap showing differential protein expression from global proteomics analysis of mHep from *Ptprf^ΔHep^* or control livers. **(k)** Heatmap of phosphoserine site differences from phosphoproteomic analysis of mHep from *Ptprf^ΔHep^* or control livers. **(l-o)** Volcano plots of global proteomic (l), phosphoserine (m), phosphothreonine (n), or phosphotyrosine (o) changes in mHep from *Ptprf^ΔHep^* or control livers. **(p)** GO enrichment of proteins altered in global proteomics in mHep from *Ptprf^ΔHep^* or control livers. **(q)** KEGG pathway enrichment of proteins with differentially regulated phosphoserine sites in mHep from *Ptprf^ΔHep^*or control livers. Bar graphs display quantification as mean ± SEM with individual data points. Statistical significance was determined using an unpaired t test (panels a, b, e, f, h and i) or One-way ANOVA (panels c and d). \**P* < 0.05, \*\**P* < 0.01, \*\*\**P* < 0.001.

We next assessed whether changes in lipid accumulation are due to changes in hepatic expression of lipid metabolism genes. Hepatic *Pparα* and *Cpt1* were significantly upregulated in *Ptprf^ΔHep^* mice on HFHFHCD (**Fig. 4e**), consistent with increased mitochondrial long-chain acyl-CoA uptake and fatty acid oxidation. *Nrf2* was also elevated, suggesting an adaptive antioxidant response. By contrast, expression of lipogenic genes remained unchanged (**Fig. 4e**). These data indicate that the lower hepatic lipid content is associated with enhanced mitochondrial respiratory capacity and oxidative lipid catabolism in liver-specific PTPRF- deficient mice, rather than due to changes in lipogenic gene expression. Actin cytoskeleton dynamics exert strong control over mitochondrial morphology, positioning and bioenergetic output. In *C. elegans* and *Drosophila*, secreted VAPB-derived MSP peptides act through Lar-like receptors to promote Arp2/3-dependent actin organization and the recruitment of mitochondria to actin-rich subcellular domains [20]. *Ptprf^ΔHep^* livers from HFHFHCD displayed a more heterogeneous distribution of mitochondrial sizes, consistent with altered mitochondrial dynamics (**Fig. 4f**). Consistent with the results in the *Ptprf^ΔHep^* liver, PTPRF decreased basal and ATP-linked respiration in mHep using saturated free fatty acids as nutrient, indicating reduced mitochondrial β-oxidation (**Fig. 4g**).

We next isolated mHep from HFHFHCD-fed mice separating low-fat and high-fat populations using differential centrifugation and Percoll gradients. High-fat control hepatocytes retained higher lipid content than *Ptprf^ΔHep^* cells after plating (**Fig. 4h**). Lipogenic signaling, reflected by PPARγ expression discriminated low and high fat cells in control hepatocytes but not in *Ptprf^ΔHep^* mHep (**Fig. 4i**). We performed integrated total and phosphoproteomic profiling of high-fat hepatocytes from control and *Ptprf^ΔHep^* mice (**Supplementary Fig. 3r-u**). Total proteome (**Fig. 4j**) and phosphoserine (**Fig. 4k**) profiles revealed distinct clusters of differentially expressed proteins. In *Ptprf^ΔHep^* mHep, global proteomics detected 70 upregulated and 42 downregulated proteins, and phosphoproteomics identified 97 upregulated and 113 downregulated phosphoserine sites, 7 up and 7 down phosphothreonine sites, and 5 up and 3 down phosphotyrosine sites (**Fig. 4l–n**). Among phosphotyrosine-modified proteins, Acad11, a mitochondrial acyl-CoA dehydrogenase supporting β-oxidation, and paxillin, a focal adhesion adaptor linking integrins to actin filaments, were increased (**Fig. 4o**). Interestingly, paxillin was also detected in the PTPRF interactome (**Supplementary Fig. 3f**), and rescue of PTPRF in *Ptprf^ΔHep^*mHep reduced phospho-paxillin (**Supplementary Fig. 3v**), supporting paxillin as a direct substrate of PTPRF. Global proteomics showed downregulation of acyl-CoA metabolic pathways, fatty acid biosynthesis, and innate immune response, while actin filament-based processes were increased in *Ptprf^ΔHep^*mHep (**Fig. 4p**). The phosphoserine-enriched dataset showed changes in insulin signaling, ErbB signaling, and growth hormone pathways, as well as pathways involved in junctional integrity: gap, adherents, and tight junctions (**Fig. 4q**). Taken together, integrated proteomic and phosphoproteomic analyses are consistent with remodeling of metabolic, cytoskeletal, and junctional signaling networks in PTPRF-deficient hepatocytes, supporting a model where they contribute to metabolic protection.

To determine whether the molecular programs discovered using the PTPRF-deficient mouse model are present in human disease, we performed proteomic and phosphoproteomic profiles of human liver samples (healthy, steatosis, MASH, **Supplementary Fig. 4a-s**). Steatotic liver samples showed increased lipid metabolism proteins (PLIN2, FASN, ACLY), while MASH liver samples have upregulated inflammatory and remodeling proteins (FGG, VTN, CTNNB1) and altered immune and cytoskeletal proteins (IGHG2, ITIH2, KRT17, **Supplementary Fig. 4t**). We compared our human and mouse proteomic datasets and found that *Ptprf^ΔHep^*mHep reversed a subset of pathological protein modifications observed in the human disease, particularly those linked to junctional integrity and actin-associated signaling (**Supplementary Fig. 4u**). These analyses support the observations that PTPRF loss modulates pathways that are dysregulated in MASLD, selectively mitigating phosphorylation and protein-abundance signatures associated with disease progression.

### PTPRF constrains insulin secretion under nutrient overload in β cells, consistent with an actin-linked mechanism

Given the importance of the endocrine pancreas in response to nutrient overload, we next assessed if a similar regulatory axis operates in cytoskeleton-dependent pancreatic β-cell function. RNA-sequencing datasets from pancreatic islets identified PTPRF expression increased in β cells from type 2 diabetic donors compared to obese non-diabetic β cells (**Fig. 5a-c**). The UPR is dysfunctional in type 2 diabetic β-cells under ER stress [26, 27]. XBP-1s maintains β-cell identity and is decreased in islets from type 2 diabetic patients [26, 27]. To determine whether PTPRF is dynamically regulated during human β-cell stress, we examined its relationship with UPR signaling. Thapsigargin treatment activated the UPR in human islets, as shown by increased XBP1s and BiP expression, and reduced PTPRF protein levels (**Fig. 5d**). A similar stress-dependent downregulation of PTPRF was observed with thapsigargin in primary mouse islets and tunicamycin in human EndoC-βH1 β-cells (**Supplementary Fig. 5a**, **b**). Consistent with this, adenoviral overexpression of XBP1s in human EndoC-βH1 β-cells decreased PTPRF expression (**Supplementary Fig. 5c**), placing PTPRF under control of UPR/XBP1s signaling in β cells, in line with our findings in hepatocytes.

**Figure 5.**
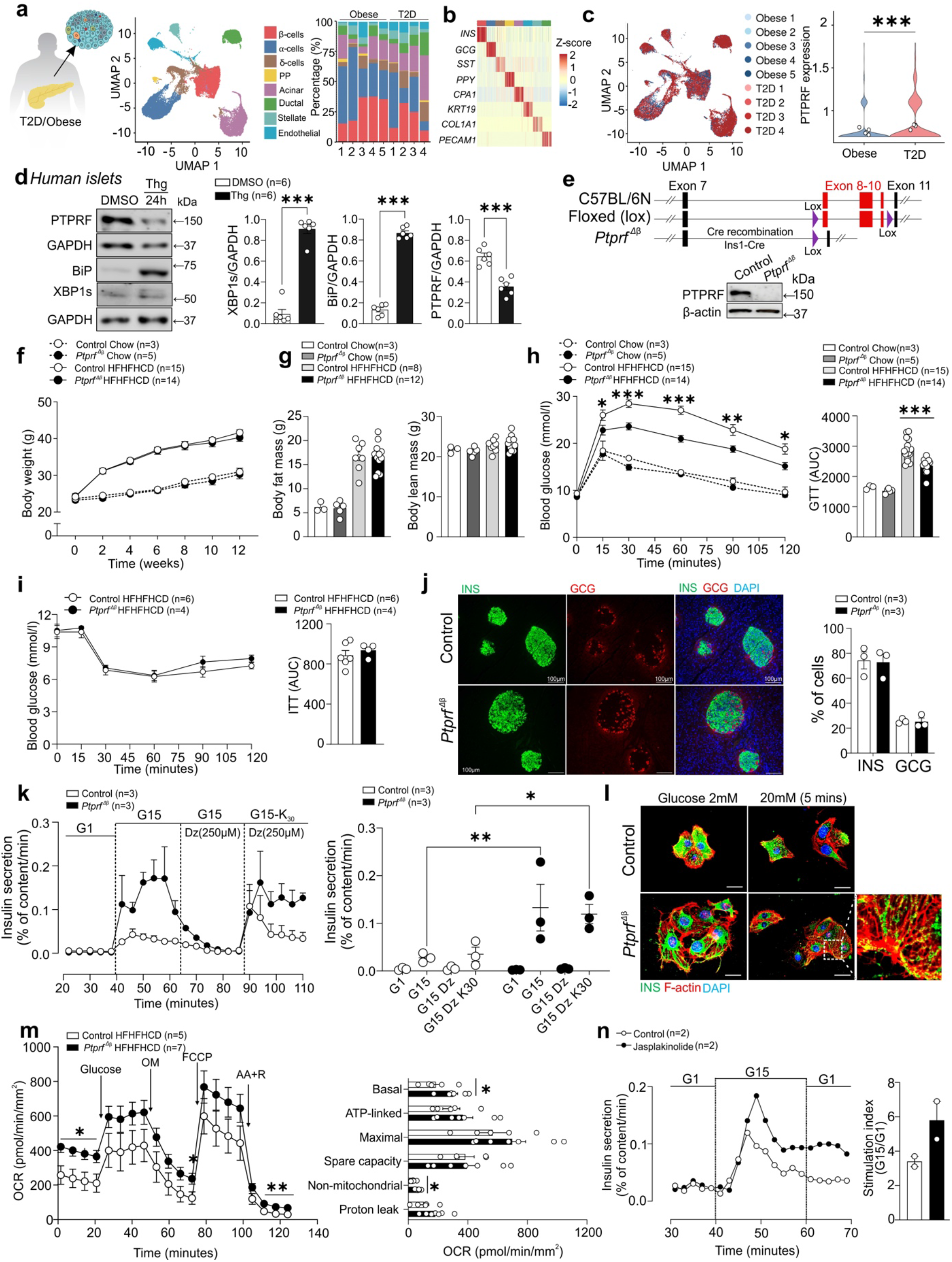
PTPRF is regulated by the unfolded protein response and constrains actin- dependent insulin secretion in β cells under nutrient overload. **(a)** UMAP representation of integrated single-cell RNA sequencing datasets from human pancreatic islets across obese and type 2 diabetic donors, identifying major endocrine and non-endocrine pancreatic cell populations. Stacked bar plots indicate the relative proportion of each cell type across conditions. **(b)** Heatmap of canonical marker gene expression across identified cell clusters, confirming annotation of β cells (INS), α cells (GCG), δ cells (SST), PP cells (PPY), acinar cells (CPA1), ductal cells (KRT19), stellate cells (COL1A1), and endothelial cells (PECAM1) **(c)** UMAP projection colored by donor group (obese and type 2 diabetes) and corresponding violin plot showing PTPRF expression in β cells across conditions. **(d)** Immunoblot analysis of PTPRF protein expression in human islets following induction of endoplasmic reticulum stress with thapsigargin, alongside markers of unfolded protein response activation (BiP and spliced XBP1). **(e)** Schematic of β-cell–specific deletion of *Ptprf* using the Cre–loxP system. LoxP sites flank exons 8–10 of the *Ptprf* gene, and Cre-mediated recombination results in excision of this region and generation of a loss-of-function allele. *Ptprf^flox/flox^* mice were crossed with *Ins1- Cre* mice to achieve β-cell–specific deletion (*Ptprf^Δβ^*). Representative immunoblot showing the absence of PTPRF protein in isolated islets from *Ptprf^Δβ^* mice compared with controls. **(f–h)** Body weight (f), body composition (g), and intraperitoneal glucose tolerance tests (h) with corresponding AUC quantification in *Ptprf^Δβ^* and littermate control mice maintained on chow or HFHFHCD over 12 weeks. **(i)** Insulin tolerance tests in *Ptprf^Δβ^* and littermate control mice following HFHFHCD feeding with corresponding AUC quantification. **(j)** Representative immunofluorescence images of pancreatic sections and quantitative proportion of insulin- positive β cells (INS⁺) and glucagon-positive α cells (GCG⁺) in *Ptprf^Δβ^* and control mice following 12 weeks HFHFHCD feeding. Scale bars: 100 µm. **(k)** Dynamic perifusion profiles of insulin secretion (4 min intervals) following sequential stimulation with low glucose (G1), high glucose (G15), diazoxide (Dz), and KCl (K_30_) on isolated *Ptprf^Δβ^* and control islets after HFHFHC feeding for 12 weeks. Scatter plot of insulin secretion across conditions. **(l)** Representative confocal images of isolated β cells from control and *Ptprf^Δβ^* mice stained with SPY555 to visualize F-actin (red), insulin (INS, green), and nuclei (DAPI, blue). Scale bar: 50 μm. **(m)** Oxygen consumption rate trace from Seahorse mitochondrial stress test in primary mouse islets from *Ptprf^Δβ^* and control mice and quantification of mitochondrial parameters. **(n)** Dynamic perifusion analysis of insulin secretion from human islets treated acutely with jasplakinolide during glucose stimulation (G1, G15). In c, d, f-n results are shown as means ± SEM. Statistical analyses were done using two-tailed unpaired Student’s t test (c, d m), one-way ANOVA (g-j) or two-way repeated measures ANOVA plus Tukey’s test (k) and denoted as \**P* < 0.05, \*\**P* < 0.01, \*\*\**P* < 0.001.

We next generated mice with β-cell-specific deletion of *Ptprf* (*Ptprf^Δβ^*) and confirmed efficient loss of PTPRF protein in isolated islets by immunoblotting **(Fig. 5e)**. *Ptprf^Δβ^* mice and littermate controls were then maintained on either a chow or HFHFHCD. Under chow-diet conditions, *Ptprf^Δβ^*mice were comparable to controls in body weight, body composition, and glucose tolerance, consistent with preserved basal glucose homeostasis (**Fig. 5f-h**). By contrast, after HFHFHCD feeding, *Ptprf^Δβ^* mice showed significantly improved glucose tolerance, as shown by lower blood glucose levels in glucose tolerance tests and reduced area under the curve, despite comparable weight gain and adiposity with control mice (**Fig. 5f-h**). Metabolic cage analyses revealed no genotype-dependent differences in energy balance under HFHFHCD conditions, excluding this as a driver of improved glucose tolerance (**Supplementary Fig. 5d- i**). Insulin tolerance was unchanged, ruling out altered peripheral insulin sensitivity (**Fig. 5i**). Islet architecture, endocrine composition, whole pancreatic insulin content, and circulating insulin levels were also comparable (**Fig. 5j**; **Supplementary Fig. 5j**, **k**). We next determined whether basal cytoskeletal organization was altered in β cells. *Ptprf^Δβ^* islets displayed an increased F/G-actin ratio compared with controls (**Supplementary Fig. 5l**), indicating enhanced actin polymerization at baseline, similar to hepatocytes. Dynamic perifusion revealed enhanced glucose-stimulated insulin secretion (GSIS) from *Ptprf^Δβ^* islets in response to glucose and KCl-induced depolarization, with only a modest effect under chow conditions (**Supplementary Fig. 5m**) and a markedly amplified phenotype following HFHFHCD feeding (**Fig. 5k**). By contrast, cytosolic Ca²⁺ dynamics were largely similar between genotypes under HFHFHCD conditions, suggesting that enhanced secretion is not driven by altered Ca²⁺ handling (**Supplementary Fig. 5n**). Consistent with the increased F/G-actin ratio, SPY555 staining showed enhanced cortical F-actin organization in *Ptprf^Δβ^* β cells (**Fig. 5l**).

Mitochondrial respiration was unchanged under chow conditions (**Supplementary Fig. 5o**), but a significantly higher basal oxygen consumption was noted, together with increased non- mitochondrial respiration in *Ptprf^Δβ^*islets following HFHFHCD feeding (**Fig. 5m**), consistent with a modest remodeling of β-cell metabolic function under nutrient overload.

Cortical actin organization restricts insulin granule access to the plasma membrane. We therefore tested whether a perturbation of actin dynamics would phenocopy PTPRF loss. In human islets, acute treatment with jasplakinolide markedly potentiated GSIS during perifusion (**Fig. 5n**). Jasplakinolide treatment also reduced PTPRF protein levels in both human and mouse islets, indicating that cytoskeletal perturbation feeds back to suppress PTPRF expression (**Supplementary Fig. 5p**, **q**). Together, these data identify PTPRF as a stress-responsive constraint on actin-dependent insulin secretion that is selectively relieved under nutrient overload.

### Distinct β-cell adaptive states in PTPRF-deficient islets under nutrient stress

To define the transcriptional programs underlying β-cell adaptation to chronic nutrient stress, single-cell RNA sequencing of pancreatic islets isolated from *Ptprf^Δβ^* and control mice following HFHFHCD feeding was performed. Unsupervised clustering identified the major endocrine populations, including α, β, and δ cells (**Fig. 6a-b**), and overall endocrine cell composition was comparable between genotypes (**Fig. 6c**). Subclustering of β cells revealed multiple transcriptionally distinct β-cell states, consistent with functional heterogeneity under metabolic stress. Differential abundance analysis demonstrated a genotype-dependent redistribution of β cells, with PTPRF-deficient islets enriched in specific β-cell states, relative to controls (**Fig. 6d**). Further analysis resolved five transcriptionally distinct β-cell clusters across *Ptprf^Δβ^* and control islets (**Fig. 6d**, **g**). Cluster-2 was strongly enriched in *Ptprf^Δβ^* islets (∼63% versus ∼3% in controls in the integrated data) and showed upregulation of pathways related to actin cytoskeleton remodeling, focal adhesion, insulin signaling, and vesicle trafficking, consistent with enhanced secretory competence. By contrast, cluster-3 was preferentially enriched in control islets (∼35% versus ∼12% in *Ptprf^Δβ^* islets in the integrated data) showing strong activation of ER stress and UPR pathways, indicative of β-cell stress under metabolic load. Additional clusters included a transcriptionally intermediate β-cell population, a cluster (cluster-4) with an exocrine-like transcriptional signature that was predominantly represented in control islets, and a small proliferative β-cell cluster present in both genotypes (**Fig. 6d**, **g**). Such stress-associated loss of β-cell identity has been reported under chronic metabolic stress and is characterized by partial adoption of non–β-cell transcriptional programs rather than true lineage conversion [28].

**Figure 6.**
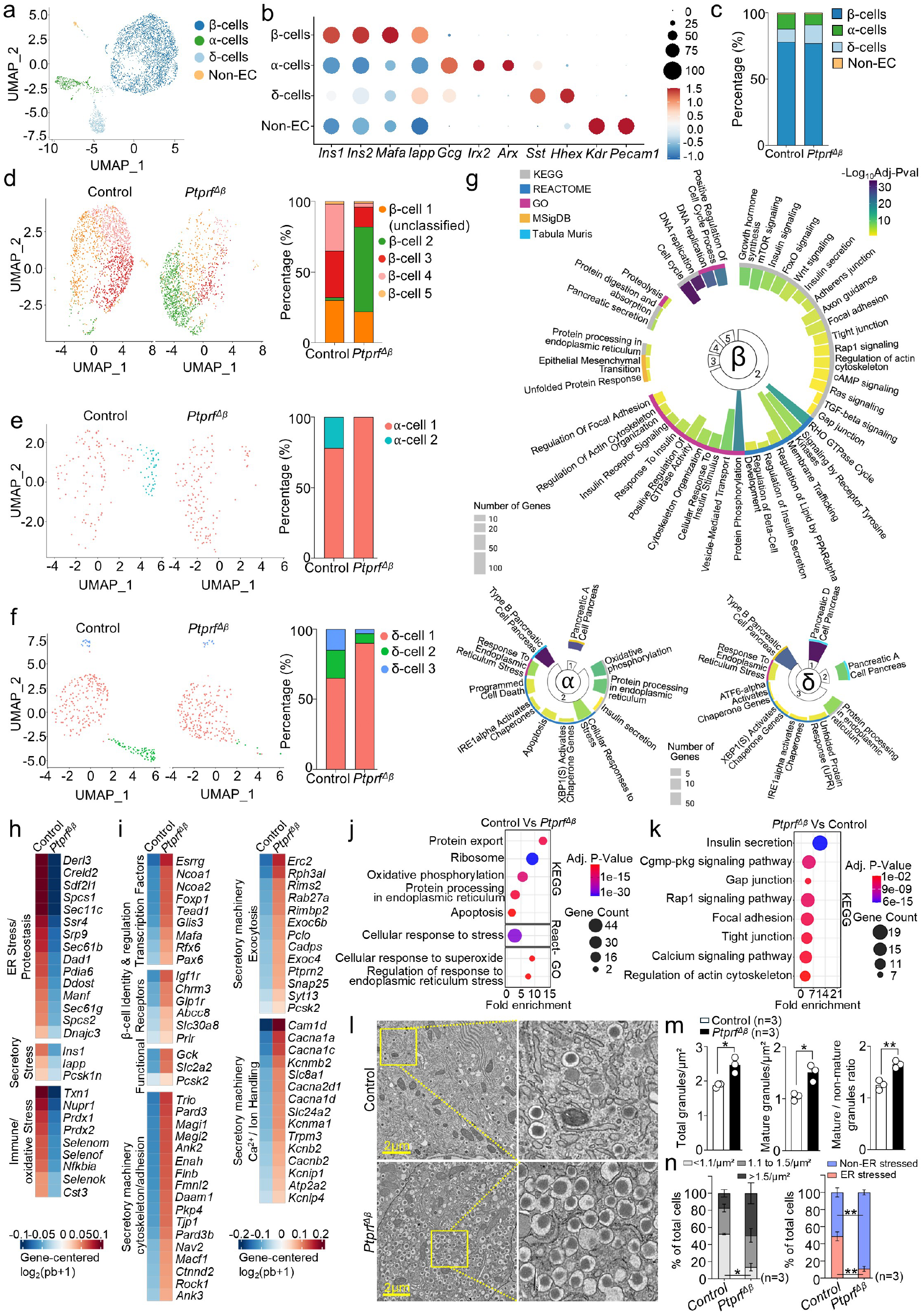
PTPRF deficiency promotes adaptive β-cell transcriptional states and preserves secretory ultrastructure under nutrient stress. **(a)** UMAP projection of integrated single- cell transcriptomes from *Ptprf^Δβ^* and control islets, annotated by major endocrine cell types, including β, α, and δ cells. **(b, c)** Relative proportions of major endocrine cell populations in *Ptprf^Δβ^* and control islets (b), showing comparable overall endocrine composition between genotypes (c). **(d-f)** UMAP projection and differential abundance analysis of β-cell clusters (d), α-cell clusters (e), and δ-cell clusters (f) demonstrating genotype-dependent redistribution of cell states in *Ptprf^Δβ^* islets relative to controls. **(g)** GO and pathway enrichment analyses of the genes significantly upregulated with a log₂ fold change > 0.25 and adjusted P < 0.05 across endocrine populations in different clusters highlighting stress-associated transcriptional programs in the integrated dataset. **(h, i)** Heatmap showing expression of representative ER/immune/oxidative stress, β-cell identity, cytoskeletal, and insulin secretory pathway genes significantly upregulated with a log₂ fold change > 0.25 and adjusted P < 0.05 between control (h) and *Ptprf^Δβ^* (i) β cells. **(j)** GO and pathway enrichment analyses of the genes significantly upregulated with a log₂ fold change > 0.25 and adjusted P < 0.05 in control β cells showing upregulation of stress-associated gene programs. **(k)** Pathway enrichment analyses of the genes significantly upregulated with a log₂ fold change > 0.25 and adjusted P < 0.05 in *Ptprf^Δβ^* β cells showing upregulation of gene programs related to actin cytoskeleton remodeling, focal adhesion, vesicle trafficking, insulin secretion, and metabolic adaptation. **(l)** Transmission electron microscopy analysis of β-cell ultrastructure from *Ptprf^Δβ^* and control mice following HFHFHC feeding. Representative TEM images of control at 10,000x magnification. Inset shows enlarged views of representative insulin granules. Scale bar: 2 µm. **(m)** Quantification of β-cell ultrastructural parameters: global insulin granule abundance, mature granule abundance, mature vs non-mature granules ratio. **(n)** β-cell segregation based on their ultrastructural features: measured density of granules and visible ER stress. Data represent mean values from three independent mice per genotype, with 60–80 β cells analyzed per mouse. In m, n results are shown as means ± SEM. Statistical analyses were done using two-tailed unpaired t-test with Welch’s correction (m, n) and denoted as \**P* < 0.05, \*\**P* < 0.01.

Consistent with cluster-level findings, control β cells showed enrichment of gene programs linked to ER stress, proteostasis imbalance, ribosomal and protein synthesis overload, and mitochondrial stress/dysfunction (**Fig. 6h**, **j**; **Supplementary Fig. 6a**). By contrast, β cells from *Ptprf*-deficient islets were enriched for gene programs related to actin cytoskeleton remodeling, vesicle trafficking, insulin granule mobilization, and regulated exocytosis, accompanied by increased expression of core β-cell identity and insulin secretion genes (**Fig. 6i**, **k**; **Supplementary Fig. 6b**). These transcriptional changes occurred without concomitant induction of ER or mitochondrial stress programs, indicating a shift toward an adaptive β-cell state under nutrient overload.

To determine whether β-cell–specific *Ptprf* gene inactivation is associated with broader effects on islet endocrine homeostasis, we examined transcriptional states of α- and δ-cell populations. In control islets, α- and δ-cell clusters displayed pronounced enrichment of pathways associated with ER stress, UPR activation, and apoptotic signaling under HFHFHCD (**Fig. 6e–g**). By contrast, these stress-associated signatures were less prominent in α- and δ-cell clusters from *Ptprf*-deficient islets (**Fig. 6e–g**), consistent with reduced islet-level stress in the context of enhanced β-cell adaptation.

To validate these findings at the ultrastructural level, we performed transmission electron microscopy of pancreatic islets from HFHFHCD-fed mice. Mitochondrial ultrastructure and the proportion of cytoplasmic area occupied by mitochondria were comparable between genotypes (**Supplementary Fig. 6c**, **d**). Quantitative analysis demonstrated that *Ptprf^Δβ^* β cells contained a significantly higher number of insulin granules per cytoplasmic area together with an increased proportion of morphologically mature granules compared with controls (**Fig. 6l**, **m**; **Supplementary Fig. 6e**), consistent with the enrichment of secretory and vesicle trafficking programs identified in adaptive β-cell clusters by single-cell transcriptomics. Quantification of granule distribution relative to the plasma membrane also showed a significant increase in membrane-proximal granules (0–100 nm) in *Ptprf^Δβ^* β cells (**Supplementary Fig. 6f**). Classification of β cells based on granule density revealed a significant reduction in cells with a low granule number (<1.1 granules/µm²) and a corresponding increase in granule-dense cells (>1.5 granules/µm²) in *Ptprf*-deficient islets (**Fig. 6n**). This indicates a shift in β-cell granule organization and cell-state heterogeneity without altering the total pancreatic insulin pool. In parallel, ultrastructural features consistent with ER stress were significantly less frequent in *Ptprf^Δβ^*β cells, with a reciprocal increase in cells lacking ER stress morphology (**Fig. 6n**; **Supplementary Fig. 6g**), indicating reduced engagement of stress-associated structural programs. Together, these data indicate that PTPRF loss selectively enhances insulin granule maturation and storage while limiting ER-stress–associated ultrastructural remodeling under nutrient overload.

### PTPRF restricts transcriptional remodeling and in vivo maturation of human stem cell– derived islets

Stem cell–derived islets (SC-islets) are a promising renewable source of human β cells for disease modelling and cell replacement therapies in diabetes. Although cytoskeletal remodeling has been implicated in endocrine induction during pancreas development and human stem-cell differentiation into β-cells [29–31], whether cytoskeleton-associated checkpoints regulate the timing and efficiency of β-cell functional maturation remains unclear. Thus, we examined the impact of PTPRF inactivation in human stem cell–derived β-like cells and SC-islets. We generated PTPRF-deficient H1-human embryonic stem cell (hESC) lines using CRISPR– Cas12a-mediated gene editing (**Supplementary Fig. 7a**), with no detectable off-target effects at predicted Cas12a sites (**Supplementary Fig. 7b**). Efficient PTPRF deletion was confirmed at both mRNA and protein levels in undifferentiated hESCs (**Fig. 7a**), which increased F/G- actin ratio compared to controls (**Fig. 7b**). *PTPRF*-deficient and control hESCs were differentiated through a multistage protocol toward pancreatic endocrine lineages. PTPRF protein expression was increased after definitive endoderm specification in control cells (**Supplementary Fig. 7c**). However, PTPRF deletion did not affect cell viability or differentiation efficiency across pluripotent (Stage 0, S0), definitive endoderm (S1), pancreatic progenitor (S4), or SC-islet (S7) (**Fig. 7c**, **d**; **Supplementary Fig. d–f**), indicating that PTPRF is dispensable for *in vitro* endocrine lineage specification.

**Figure 7.**
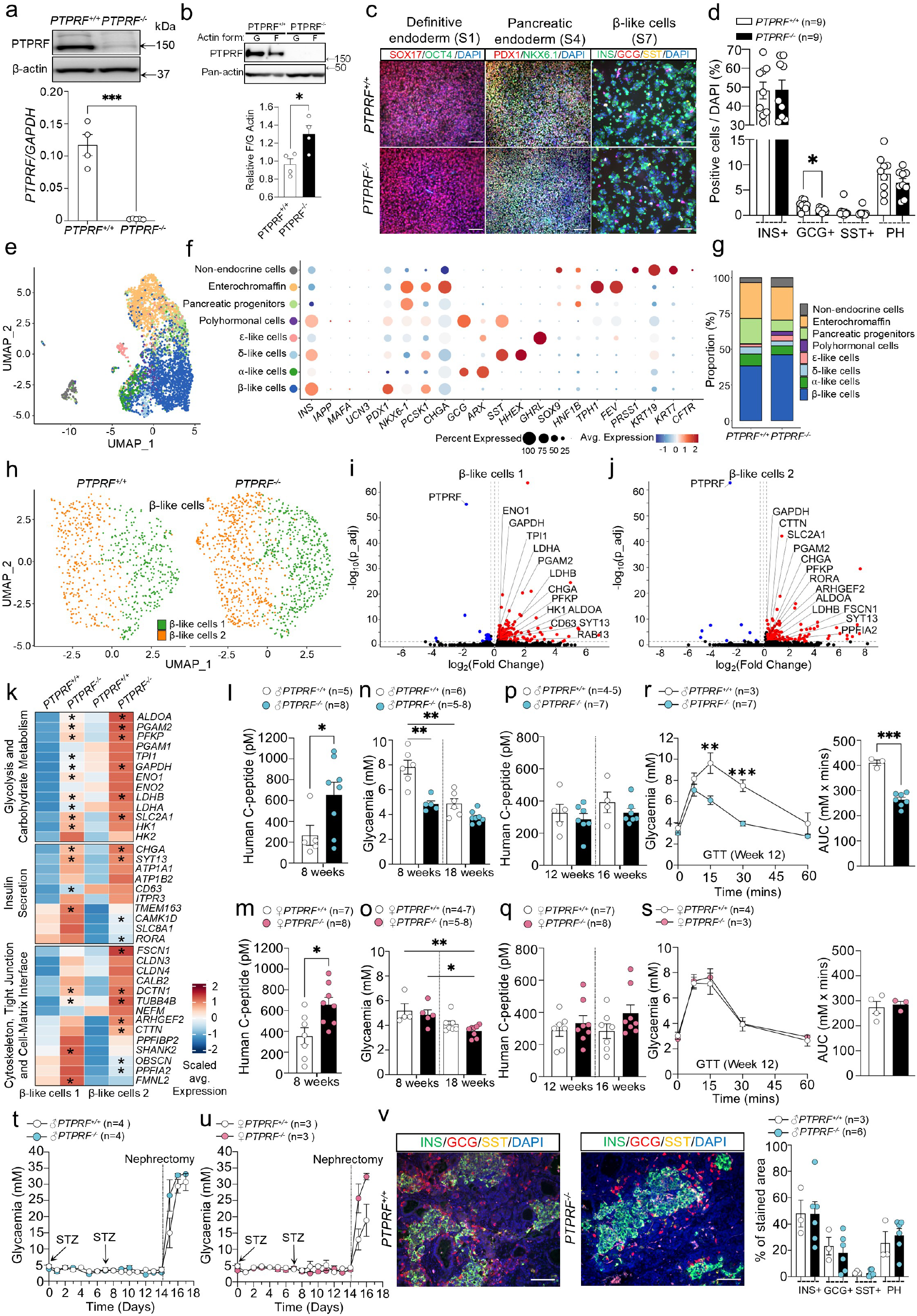
PTPRF deletion remodels transcriptional programs in stem cell-derived islets and enhances *in vivo* metabolic remodeling. (a) Western blot and qPCR for PTPRF in *PTPRF^-/-^* and *PTPRF^+/+^* H1 hESC clones. **(b)** Immunoblot analysis of F actin and G actin fractions from *PTPRF^-/-^*and *PTPRF^+/+^* H1 hESC. **(c)** Immunofluorescence analysis of differentiation stages showing SOX17⁺/OCT4⁻ definitive endoderm (stage 1), PDX1⁺/NKX6.1⁺ pancreatic progenitors (stage 4) and INS⁺, GCG⁺ or SST⁺ endocrine cells (stage 7). Scale bar: 100 µm. **(d)** Quantification of endocrine populations at stage 7 obtained by CellProfiler analysis. The proportion of β-like (INS⁺), α-like (GCG⁺), δ-like (SST⁺) and polyhormonal (PH) cells is shown for *PTPRF^-/-^* and *PTPRF^+/+^* SC-islets. **(e)** UMAP projection of integrated single- cell transcriptomes from *PTPRF^-/-^* and *PTPRF^+/+^* stage-7 SC-islets. **(f)** Expression of canonical markers used to assign cluster identities, showing transcriptionally distinct endocrine and non- endocrine clusters, including β-like, α-like, δ-like, ductal-like and mesenchymal-like populations. **(g)** Proportions of major cell populations in *PTPRF^-/-^* and *PTPRF^+/+^* SC-islets. **(h)** UMAP split by genotype showing two distinct transcriptional signatures between *PTPRF^-/-^* and *PTPRF^+/+^* β-like endocrine clusters. **(i, j)** Differential gene expression analysis displayed as volcano plots for *PTPRF^-/-^* and *PTPRF^+/+^* β-like cells 1 cluster (i) and β-like cells 2 cluster (j). **(k)** Expression profiles of representative cytoskeletal, glycolytic and insulin secretory pathway genes altered in *PTPRF^-/-^* β-like cells 1 cluster and β-like cells 2 cluster. Genes marked with an asterisk are significantly upregulated in *PTPRF^-/-^* SC-islets (log₂ fold change > 0.25 and adjusted P < 0.05). **(l, m)** Non-fasted plasma human C-peptide concentrations of male (l) or female (m) mice implanted with *PTPRF^-/-^* or *PTPRF^+/+^* SC-islets after 8 weeks. **(n, o)** Baseline glycaemia measured in non-fasted male (n) and female (o) mice at 8 and 16 weeks post- implantation. **(p, q)** Fasted plasma human C-peptide levels measured at 12- and 16-weeks post- implantation in male (p) and female (q) mice. **(r, s)** Intraperitoneal glucose tolerance tests performed 12 weeks after implantation, with corresponding area-under-the-curve quantification in male (r) and female (s) mice. **(t, u)** Blood glucose measurements during streptozotocin treatment to ablate endogenous murine β cells, followed by unilateral nephrectomy to remove the graft-bearing kidney in male (t) and female (u) mice. **(v)** Representative images of retrieved SC-islet grafts from kidneys of male mice implanted with *PTPRF^-/-^*and *PTPRF^+/+^* SC-islets and the quantification of endocrine cell populations. Scale bars: 100 µm. In a, b, d, l-v results are shown as means ± SEM. Statistical analyses were done using two-tailed unpaired t-test (a, b, d, l, m, r, s, v) or two-way ANOVA (n-q) and denoted as \**P* < 0.05, \*\**P* < 0.01, \*\*\**P* < 0.001.

Single-cell RNA sequencing of stage-7 SC-islets and unsupervised clustering identified major endocrine and non-endocrine populations with comparable distributions between genotypes (**Fig. 7e-g**; **Supplementary Fig. 7g**). Within the β-like cell population, both *PTPRF*-deficient and control cells segregated into two distinct transcriptional states (**Fig. 7h**; **Supplementary Fig. 7h**). Differential gene expression analysis of the two β-like clusters revealed marked differences in gene programs associated with β-cell maturation (**Supplementary Fig. 7i**). β- like cell cluster 1 was enriched for genes involved in β-cell development and identity, consistent with a developmentally associated β-like state. By contrast, β-like cell cluster 2 exhibited higher expression of genes linked to insulin production and secretion, and components of the secretory stress response, consistent with a more functionally mature β-like transcriptional program. Differential gene expression and pathway analyses revealed enrichment of gene programs associated with cytoskeletal organization, glucose metabolism, and insulin secretory function in *PTPRF*-deficient β-like cells across both identified β-like clusters and the merged β-like population (**Fig. 7i–k**; **Supplementary Fig. 7j–m**). These results suggest that PTPRF loss remodels β-like cell transcriptional states toward enhanced functional metabolic remodeling rather than altering lineage allocation.

We next assessed whether these transcriptional changes translate into functional outcome *in vivo*. *PTPRF*-deficient and control SC-islets were implanted under the kidney capsule of immunodeficient NOD-SCID mice of both sexes. Because of differences in the timing of functional maturation between sexes [32], we analyzed male and female cohorts separately. *PTPRF*-deficient grafts showed significantly higher plasma human C-peptide levels at 8 weeks post-implantation in both male and female recipients compared with control grafts (**Fig. 7l**, **m**), indicating accelerated onset of functional insulin secretion. Longitudinal assessment revealed convergence of basal human C-peptide levels between genotypes at later time points (12–16 weeks) in both sexes (**Fig. 7p**, **q**), indicating that PTPRF inactivation accelerates metabolic kinetics without increasing long-term basal secretory output.

In female recipients, this early increase in insulin secretion was accompanied by rapid normalization of basal glycaemia, with both *PTPRF*-deficient and control grafts reaching comparable human-range glycemic set points (∼5 mM) by 8 weeks post-implantation and no sustained genotype-dependent differences in glucose tolerance thereafter (**Fig. 7o**, **s**). By contrast, male recipients implanted with *PTPRF*-deficient SC-islets reached the basal human- range glycemic set point by 8 weeks and displayed significantly enhanced glucose tolerance at 12 weeks post-implantation relative to controls (**Fig. 7n**, **r**), whereas control grafts required a longer engraftment period to achieve comparable metabolic regulation.

*In vivo* glucose-stimulated insulin secretion assessed 12- and 16-weeks post-implantation revealed comparable responses between genotypes in female recipients and convergence of responses in male recipients by 16 weeks (**Supplementary Fig. 7n**, **o**). As GSIS assays probe maximal secretory capacity, whereas glucose tolerance tests reflect integrated physiological glucose control, these data indicate that PTPRF deletion accelerates the timing of *in vivo* β-cell functional metabolic remodeling without increasing maximal insulin secretory capacity regardless of sex.

To confirm graft dependence of glucose homeostasis, endogenous murine β-cells were ablated using streptozotocin, followed by nephrectomy of the graft-bearing kidney. *PTPRF*-deficient and control grafts maintained normoglycemia following streptozotocin treatment, whereas graft removal resulted in rapid hyperglycemia in both sexes (**Fig. 7t**, **u**). Immunostaining of retrieved grafts confirmed strong insulin-positive endocrine tissue with no changes in the relative distribution of endocrine cells between the genotypes (**Fig. 7v**).

Together, these data show that PTPRF contributes to actin remodeling in stem cells and, while dispensable for human β-cell lineage specification *in vitro*, regulates the transcriptional and functional metabolic remodeling of SC-islets.

## Discussion

Our work advances the field by identifying PTPRF as a regulator of the actin-supported cortical cytoskeleton at the plasma membrane, linking cytoskeletal organization to insulin signaling, mitochondrial engagement, and lipid routing in hepatocytes. Simultaneously, PTPRF controls actin-dependent insulin secretion in primary β cells and contributes to metabolic reprograming in SC-islets. Together, our data support a model in which PTPRF functions as an energetically restrictive cortical checkpoint that stabilizes actin–adhesion assemblies under basal conditions but becomes maladaptive during chronic nutrient excess. Metabolic homeostasis requires cells to continuously align energetic demand with structural dynamics. This is particularly critical in hepatocytes and pancreatic β cells, which serve as primary sensors and executors of nutrient availability. Hepatocytes rapidly reorganize membrane domains, lipid handling machinery, and mitochondrial positioning in response to nutrient excess, while β cells repeatedly mobilize insulin granules through actin-supported cortical barriers during sustained secretory demand. In both cell types, these adaptive responses rely on dynamic actin cytoskeleton remodeling, impose localized ATP consumption, and depend on tightly coordinated regulatory events.

PTPRF deficiency enables a more dynamic, energetically adaptive actin architecture. Although the downstream consequences diverge in hepatocytes and β cells in accordance with their specialized functions, they converge on a common physiological outcome. Specifically, this enhances resilience to chronic nutrient excess and mitigates diet-induced metabolic dysfunction. Thus, PTPRF is a shared regulator of metabolic adaptability in these two cell types that functions through cell-intrinsic cytoskeletal mechanisms rather than classical transcriptional control of metabolic enzymes. This potential dual catalytic and scaffolding function restricts adaptor mobility, stabilizes adhesion complexes, and dampens actin remodeling dynamics.

In obesity, hepatocytes accommodate large fluxes of glucose and lipids while maintaining insulin responsiveness and preventing excessive triglyceride storage. Early steatosis disrupts both microtubules and F-actin, displacing mitochondria and triggering a compensatory increase in basal and maximal respiration that persists even after lipid removal and apparent cytoskeletal restoration [9, 33]. The cytoskeleton not only adapts to the nutrient status but actively feeds back into metabolic regulation [34, 35]. PTPRF limits insulin receptor phosphorylation and attenuates AKT activity during nutrient stress, thereby diverting substrate handling toward triglyceride storage rather than oxidative metabolism. PTPRF deficiency redistributes metabolic flux characterized by elevated mitochondrial respiration, and reduced glycolysis. In this context, hepatocytes lacking PTPRF avoid maladaptive insulin resistance that typically follows nutrient overload. These effects are particularly striking given that PTPRF suppression itself is a stress-responsive event. Under chronic nutrient pressure, maintaining rigid adhesion and actin structures imposes a sustained energetic burden. By relieving PTPRF-mediated constraints, hepatocytes increase focal adhesion turnover, enhance mitochondrial engagement, and redistribute lipid processing, directing metabolism towards oxidation. This provides a mechanistic explanation for why mice lacking hepatic PTPRF remain leaner and accumulate less hepatic fat despite equivalent caloric intake.

Pancreatic β cells must dynamically balance insulin secretory efficiency when nutrient availability fluctuates. Failure to appropriately adapt this balance during chronic nutrient excess contributes to β-cell dysfunction and progression toward type 2 diabetes [36]. Consistent with this, analysis of human single-cell RNA-sequencing datasets revealed increased PTPRF expression in β cells from donors with type 2 diabetes. PTPRF acts as a stress-responsive, cytoskeleton-associated checkpoint that constrains actin-dependent insulin secretion under basal conditions that is selectively relieved during nutrient overload, thereby enabling adaptive insulin release without pathological hypersecretion. PTPRF regulates distal steps of stimulus– secretion coupling. Cortical actin is a well-established gatekeeper of insulin granule access to release sites, acting as both a physical and regulatory barrier that is dynamically remodeled during glucose stimulation [37]. PTPRF deficiency enhances insulin secretion downstream of glucose metabolism and membrane depolarization without altering Ca²⁺ influx, placing its action at the level of actin-dependent granule mobilization and exocytosis rather than upstream glucose sensing or excitation–secretion coupling. The ability of actin stabilization to suppress PTPRF expression further suggests a feedback loop linking cytoskeletal state to secretory restraint [38]. Importantly, relieving this constraint does not result in constitutive hypersecretion. Instead, PTPRF loss improves the efficiency and timing of insulin release under metabolic stress while leaving the maximal secretory capacity largely unchanged. This distinction is critical, as chronic insulin hypersecretion has been implicated in β-cell exhaustion and failure [39]. By restricting distal exocytotic capacity when demand is high, PTPRF can limit adaptive compensation and contribute to β-cell vulnerability.

We found that the UPR, acting through XBP1s, suppresses PTPRF expression in hepatocytes and β-cells. This suggests that PTPRF loss is not merely a marker of metabolic dysfunction but is part of an adaptive response engaged when static cortical assemblies become energetically costly. ER stress is a central driver of β-cell dysfunction in obesity and type 2 diabetes [36]. Activation of the UPR, particularly XBP1s signaling [26], preserves β-cell identity and protects against stress-induced failure during metabolic challenge. We identified PTPRF as a downstream target of UPR and XBP1s signaling in β cells. The stress-dependent suppression of PTPRF places it within an adaptive UPR-regulated program rather than as a result of β-cell failure. This interpretation is reinforced by the context dependence of the phenotype, PTPRF loss is functionally silent under basal conditions, yet it confers a clear advantage when the secretory demand is elevated, consistent with a role in gating, rather than driving, insulin secretion.

At transcriptional and ultrastructural levels, PTPRF deletion biases β cells toward adaptive functional states characterized by enhanced cytoskeletal organization, vesicle trafficking, and insulin granule maturation, while limiting engagement of ER and mitochondrial stress programs [40, 41]. Rather than inducing dedifferentiation, loss of PTPRF appears to preserve β-cell identity under nutrient stress, consistent with prior reports that chronic metabolic stress can drive erosion of β-cell identity through activation of non-β-cell transcriptional programs rather than bona fide lineage conversion [28]. The reduced prevalence of stress-associated β-cell states in PTPRF-deficient islets, together with improved insulin granule density and maturation, supports the concept that cytoskeletal checkpoints regulate not only secretory output but also β-cell resilience. Extension of these findings to human SC-islets further supports a role for PTPRF in regulating functional metabolic remodeling, rather than lineage specification [29, 42]. PTPRF deletion remodels the transcriptional programs in stem cell-derived β-like cells toward enhanced metabolic and secretory competence and accelerates the acquisition of *in vivo* function. This remodeling includes upregulation of late glycolytic enzymes and glucose handling genes, relieving glycolytic bottlenecks known to constrain metabolic coupling in immature β-like cells [43, 44]. Importantly, this configuration preserves mitochondrial coupling and redox balance, as indicated by preferential expression of *LDHB* without dominant induction of *LDHA*, thereby preventing lactate shunting and basal hypersecretion [44, 45]. Loss of PTPRF selectively accelerates the timing and kinetics of β-cell functional metabolic remodeling without increasing maximal secretory capacity, revealing genotype-dependent effects that are most apparent in contexts of delayed maturation in male mouse recipients.

PTPRF may also affect cytoskeleton–metabolism coupling in other metabolically active cells, such as myocytes. In muscle cells, where actin organization is tightly linked to mitochondrial function, PTPRF could constrain cytoskeletal dynamics and substrate utilization under changing energetic demand, as demonstrated in *C. elegans* [20]. Moreover, numerous chronic conditions - such as inflammatory and neurological disorders, myocardial infarction, and cancer - are characterized by actin remodeling. Therefore, the findings of this study may have broad implications for understanding pathogenic cell adaptation.

In conclusion, PTPRF is a crucial adaptive protein engaged when metabolic work must be repeatedly deployed. Our work suggests that targeting cytoskeleton-associated checkpoints may enhance hepatocyte and β-cell resilience, without compromising long-term metabolic function. Modulation of PTPRF in defined metabolic states could provide coordinated protection of hepatic insulin sensitivity and β-cell secretory capacity; whether these effects are additive or synergistic remains to be determined. Future pharmacological or peptide-based PTPRF inhibition [46, 47] will be essential to assess its potential to prevent or delay obesity- mediated disease progression.

## Methods

### Sex as a biological variable

Male and female human liver samples and pancreatic islets were used for gene/protein expression, proteomic and histology analysis. We performed *in vivo* metabolic studies using male mice because female mice are resistant to the development of features of human obesity and liver dysfunction following obesogenic diet feeding. We performed human SC-islet implantations in male and female NOD/SCID mice. Primary hepatocytes and mouse islets were isolated from both male and female mice in this study.

### Human liver samples

Human liver samples were obtained from patients with obesity who underwent bariatric surgery at the Virgen de la Arrixaca Clinical University Hospital (Murcia, Spain) between January 2020 and December 2022. Clinical characteristics are provided in **Supplementary Tables 1 and 2**.

### Human pancreatic islets

Primary human pancreatic islets were provided by the Network for Islet Transplantation (Université Catholique de Louvain). Upon arrival, islets were cultured on ultra-low attachment plates in Ham’s F-10 nutrient mixture supplemented with bovine serum albumin, GlutaMAX, penicillin–streptomycin, fetal bovine serum and IBMX. Donor characteristics are provided in **Supplementary Table 3**, and full media composition and culture conditions are described in the Supplementary Methods. Single-cell RNA sequencing data from human pancreatic islets of nine female donors were obtained from the Human Pancreas Analysis Program (HPAP, https://hpap.pmacs.upenn.edu). Donor characteristics are provided in **Supplementary Table 4**.

### Mouse models and dietary interventions

Mice were maintained at 22 °C under a 12-h light and 12-h dark cycle with *ad libitum* access to food and water. Mice carrying a conditional *Ptprf* allele were generated inhouse for this study on a C57BL/6N background with loxP sites flanking exons 8–10 of the *Ptprf* gene (Cyagen), such that Cre-mediated recombination results in a frameshift and loss-of-function allele. Targeting was performed in C57BL/6N embryonic stem cells using homologous recombination, followed by germline transmission. A hepatocyte specific *Ptprf* conditional knockout line (*Ptprf^ΔHep^*) was generated by crossing *Ptprf* floxed mice with Alb-Cre transgenic mice (The Jackson Laboratory, B6.Cg-*Speer6-ps1^Tg(Alb-cre)21Mgn^*/J; JAX #003574). A β-cell specific *Ptprf* conditional knockout line (*Ptprf^Δβ^*) was also generated by crossing *Ptprf* floxed mice with Ins1-Cre transgenic mouse line (The Jackson Laboratory, B6(Cg)-*Ins1^tm1.1(cre)Thor^*/J; JAX #026801).

Eight-week-old control, *Ptprf^ΔHep^ or Ptprf^Δβ^* and littermate control mice were randomized to receive a reference chow (control diet), a high-fat diet (HFD), a high-fat high-fructose high- cholesterol diet (HFHFHCD) or a methionine and choline-deficient high-fat (HFMCD) diet for the indicated durations, ranging from 6 to 24 weeks.

### Metabolic phenotyping

Whole body and liver fat and lean mass were measured body composition analyzer by EchoMRI, USA. Glucose and insulin tolerance were assessed after 6h fasting using intraperitoneal glucose or insulin injections with serial tail blood glucose measurements.

Mice fed HFHFHCD were placed in metabolic cages TSE Phenomaster setup (TSE, Germany) for a duration of 72 h. Following a 24 h period of acclimatization, metabolic parameters, including physical activity, energy expenditure, and substrate utilization were assessed by indirect calorimetry. Experimental details for glucose and insulin tolerance tests and indirect calorimetry, including fasting durations, doses, and acquisition settings, are described in the Supplementary Methods.

### Cell culture

mHep were isolated by two step collagenase perfusions followed by Percoll based enrichment, and isolation details are provided in the Supplementary Methods. For experiments using lipid rich hepatocytes from steatotic livers, we followed the published protocol indicated in the Supplementary Methods. Cells were plated on collagen coated plates and maintained in William’s based media.

HepG2 cells were cultured using DMEM with 10% heat-inactivated FBS and Penicillin- Streptomycin.

### *In vitro* stimulation, genetic perturbations and viral transduction

For signaling and metabolic assays, mHep were serum starved and stimulated with combinations of insulin, glucose, BSA-conjugated saturated and unsaturated fatty acids, inflammatory cytokines or unfolded protein response inducers.

PTPRF overexpression was achieved using adenoviral vectors or plasmid-based expression of full length PTPRF fused to mGreenLantern in HepG2 cells. The active spliced form of Xbp1 was overexpressed by adenoviral delivery. Stimulation conditions, reagent concentrations, viral transduction parameters are detailed in the Supplementary Methods.

### Immunoblotting, imaging and ultrastructural analysis

For immunoblotting, cells and tissues were lysed in detergent containing buffers with protease and phosphatase inhibitors, separated by SDS PAGE and probed with antibodies listed in **Supplementary Table 5**.

Neutral lipids were visualized using Nile Red staining. Plasma membrane was labelled with the CellBrite™ Fix 555 membrane dye. The actin cytoskeleton was delineated by phalloidin based staining of filamentous F actin, and nuclei were identified by nuclear counterstaining with DAPI. For ultrastructural analysis, mice were perfusion fixed, and liver samples were processed for resin embedding, ultrathin sectioning, and transmission electron microscopy to examine mitochondrial morphology and subcellular organization. Image acquisition settings, fluorophore excitation and detection parameters, and quantification workflows are described in the Supplementary Methods.

### Seahorse analysis of mitochondrial respiration and glycolysis

Mitochondrial respiration and glycolytic activity in primary hepatocytes and islets were assessed using Seahorse XF Flex or HS Mini Analyzers (Agilent Technologies, USA). For mitochondrial stress tests, oxygen consumption rate was measured at baseline and after sequential additions of ATP synthase inhibitor, uncoupler and respiratory chain inhibitors. For glycolytic stress tests, extracellular acidification rate was recorded in response to glucose, ATP synthase inhibition and glycolytic blockade. Rates were normalized to protein content and basal, maximal and spare respiratory capacity as well as glycolytic capacity and reserve were calculated according to manufacturer’s recommendations. Detailed conditions and concentrations are included in Supplementary Methods.

### HYlight live cell imaging of fructose 1,6-bisphosphate

Dynamic changes in glycolytic flux were monitored in mouse primary hepatocytes using the genetically encoded HYlight biosensor for fructose 1,6-bisphosphate [25]. mHep expressing HYlight were imaged by ratiometric confocal microscopy under glucose stimulation, and excitation ratio changes were quantified in single cell regions of interest (ROIs) over time. Transfection conditions, imaging media, microscope settings, excitation and emission parameters, and ratio quantification workflows are described in the Supplementary Methods.

### Interactome profiling of PTPRF

To map PTPRF associated protein complexes, we used recombinant biotinylated intracellular domains of PTPRF, wild type and substrate trapping mutant (CS), immobilized on streptavidin beads. Pervanadate treated lysates from mHep or HepG2 cells were incubated with these baits, and bound proteins were eluted and identified by LC-MS.

High confidence interactors were defined based on enrichment relative to beads only and differential enrichment between wild type and trap mutant. Functional annotation and clustering of enriched interactors were performed using Gene Ontology based enrichment and term reduction. Pervanadate preparation, lysis and pull-down conditions, on bead digestion, LC-MS acquisition settings, and interactor filtering criteria are described in the Supplementary Methods.

### Global proteomics and phosphoproteomics

Global proteome and phosphoproteome profiling were performed on liver tissue from human liver samples and on high-fat mHep from control or *Ptprf^ΔHep^* mice. For mHep, phosphotyrosine sites were enriched by immunoaffinity purification and phosphoserine and phosphothreonine sites by titanium dioxide-based methods (Cell Signaling, USA). For total proteomics, tandem mass tag based labelling and high pH fractionation were used for multiplexed quantification. Peptides were analyzed on an Orbitrap mass spectrometer in data dependent acquisition mode. Raw data were processed with Proteome Discoverer and Sequest HT against UniProt reference proteomes, with false discovery rate controlled at 1%. Phosphopeptide enrichment strategies, TMT labelling and fractionation, Orbitrap acquisition parameters, database search settings, and downstream preprocessing and pathway enrichment workflows are described in the Supplementary Methods.

### Histology and immunohistochemistry

Formalin fixed paraffin embedded liver tissue from mice and humans was stained with hematoxylin and eosin and Sirius Red for assessment of steatosis, inflammation and fibrosis. PTPRF protein expression was evaluated by immunohistochemistry using in house validated antibodies (**Supplementary Table 5**), and images were captured using NanoZoomer Digital Pathology (Hamamatsu Photonics K.K., version SQ 1.0.9). Protocols and quantification criteria are described in Supplementary Methods.

### Primary mouse islet isolation

Mouse pancreatic islets were isolated by collagenase digestion. Briefly, pancreata were perfused and digested with collagenase prepared in serum-free M199 medium (Gibco Cat# 11150059), followed by repeated hand-picking under a stereomicroscope to obtain highly enriched islet preparations. Collagenase digestion conditions and islet purification procedures are described in the Supplementary Methods.

### Islet dissociation and fluorescence-activated cell sorting for single-cell RNA sequencing

For mouse islet single-cell RNA sequencing, islets were isolated from *Ptprf^Δβ^* mice and control littermates after 12 weeks of HFHFHC diet feeding and dispersed into single cells using enzymatic dissociation in the presence of DNase I (20μg/mL, Hofman-La Roche, Basel, Switzerland). Live cells were purified by fluorescence-activated cell sorting (FACS) (Becton Dickinson FACS Aria) using propidium iodide (1mg/mL, Sigma-Aldrich) exclusion with forward- and side-scatter gating to remove debris and doublets. Sorted cells were filtered to remove residual aggregates, counted, and only samples meeting viability thresholds were processed for single-cell library preparation. Dissociation conditions, viability thresholds, and FACS gating principles for exclusion of debris, dead cells, and doublets are described in the Supplementary Methods.

### Dynamic secretion experiments with isolated islets

Glucose-stimulated insulin secretion dynamics were assessed by perifusion of batches of isolated mouse or human islets at 37 °C. After equilibration in low-glucose buffer, islets were sequentially stimulated with the indicated glucose concentrations and secretagogues (including diazoxide and KCl where noted) with timed fraction collection. Secreted insulin was normalized to total islet insulin content measured at the end of the experiment. Perifusion setup, equilibration conditions, stimulation sequences, fraction collection timing, and normalization to total insulin content are described in the Supplementary Methods.

### Intracellular Ca²⁺ imaging in isolated mouse islets

Cytosolic Ca²⁺ dynamics were measured in isolated mouse islets loaded with the ratiometric dye fura-2 LR (Sigma-Aldrich). Islets were perifused in Krebs–Ringer buffer and sequentially stimulated with low and high glucose followed by diazoxide and KCl, as indicated. Fluorescence ratios were recorded in small- to medium-sized islets and quantified using dedicated imaging MetaFluor software (Molecular Devices). Imaging parameters are detailed in Supplementary Methods.

### CRISPR–Cas12a–mediated PTPRF knockout in hESCs

PTPRF-deficient H1 hESC lines (WiCell, Madison, WI) were generated using CRISPR– Cas12a ribonucleoprotein electroporation with guide RNAs targeting exon 23. Edited clones were isolated by single-cell cloning and validated by PCR and Sanger sequencing. Potential off-target sites predicted in silico were amplified and sequenced to verify genome integrity. Single guide RNA and oligonucleotide sequences are detailed in the **Supplementary Table 6.**

### Differentiation of hESCs into stem cell–derived islets and preparation for single-cell RNA sequencing

*PTPRF^-/-^* and *PTPRF^+/+^* (isogenic control) H1 hESC clones were differentiated into stem cell– derived islets (SC-islets) using a seven-stage protocol [48]. Differentiation through definitive endoderm and pancreatic progenitor stages was performed in adherent culture, followed by reaggregation and three-dimensional differentiation through endocrine induction and maturation. For single-cell RNA sequencing, stage-7 SC-islets were dissociated into single cells, filtered to remove aggregates, assessed for viability and processed for library preparation. Detailed media, timing and quality control criteria are provided in Supplementary Methods.

### SC-islet implantation and *in vivo* functional assessment

To assess *in vivo* maturation, SC-islets derived from *PTPRF^-/-^*and *PTPRF*H1 hESCs were implanted under the kidney capsule of immunodeficient NOD–SCID mice of both sexes and monitored longitudinally [48]. Plasma human C-peptide and glycaemia were measured at defined time points, and intraperitoneal glucose tolerance and *in vivo* GSIS assays were performed after standardized fasting with intraperitoneal glucose administration. To test graft dependence, endogenous murine β cells were ablated with streptozotocin followed by nephrectomy of the graft-bearing kidney. Full implantation, sampling and assay procedures are described in Supplementary Methods.

### Pancreatic and graft immunofluorescence and quantification

Mouse pancreata and SC-islet graft-bearing kidneys were fixed, paraffin embedded and sectioned. Sections were subjected to antigen retrieval, permeabilization and blocking, followed by immunofluorescence staining using antibodies listed in **Supplementary Table 5**. Images were acquired on Zeiss fluorescence platforms and endocrine composition was quantified using automated image analysis with CellProfiler. Full staining conditions are provided in Supplementary Methods.

### Pancreatic insulin content

Pancreatic insulin content was quantified from tissue extracts generated in acidified ethanol and measured using a commercial insulin ELISA (Mercodia, Cat#10-1247-01). Values were normalized to pancreas weight. Detailed extraction and assay conditions are provided in Supplementary Methods.

### Single-cell RNA sequencing and computational analysis

Mouse islet and SC-islet single-cell libraries were generated using the 10x Genomics (Pleasanton, CA) Chromium Single-Cell 3′ platform and processed using standard pipelines. Ambient RNA correction and doublet removal were performed prior to normalization, dimensional reduction and clustering. Cell types were assigned based on canonical marker expression and module scoring, and differential expression and pathway enrichment analyses were performed using established R packages with multiple-testing correction. Full computational workflows, thresholds and software versions are described in Supplementary Methods.

### Statistics

Sample size n refers to individual mice, independent hepatocyte and islet preparations, individual human liver samples or independent experiments as indicated in figure legends. Data are presented as mean ± standard error of the mean, unless indicated otherwise. Statistical comparisons between two groups were performed using two-sided Student’s t tests, and comparisons among multiple groups used one way, two way or repeated measures analysis of variance, followed by appropriate post hoc tests as indicated.

For global proteomics and phosphoproteomics, differentially abundant proteins and phosphosites were identified using moderated *t*-statistics with empirical Bayes shrinkage (limma) at a significance threshold of *P* < 0.05. For single-cell and single-nucleus RNA sequencing, differential expression was assessed using the Wilcoxon rank-sum test with Bonferroni correction for the number of genes tested, and pathway and gene set enrichment analyses were corrected by the Benjamini–Hochberg false discovery rate method. Exact *P* values are reported in figure panels and source data. Statistical tests, multiple-comparison procedures and software versions are specified in figure legends and Supplementary Methods.

### Study Approval

The study was conducted in accordance with the Declaration of Helsinki and approved by the Ethics and Clinical Research Committees of the Virgen de la Arrixaca Clinical University Hospital (reference number 2025-10-9-HCUVA).

All animal experiments complied with Belgian regulations for animal care and were approved by the Commission d’Éthique du Bien Être Animal at the Faculty of Medicine, Université libre de Bruxelles, dossiers 732N, 917N and 918N.

## Data availability

The following publicly available datasets were used: GSE135251 for bulk RNA-seq deconvolution of human MASLD liver, performed with the MuSiC algorithm using the liver single-cell atlas as reference [49]; integrated single-nucleus RNA-seq datasets from [22] for independent validation of hepatocyte-level expression; GSE150889 for mouse XBP1 ChIP-seq analysis and ENCODE dataset ENCSR988EVQ for human enhancer annotation at the *PTPRF* locus; and the Human Pancreas Analysis Program (HPAP) repository for single-cell RNA sequencing of human pancreatic islets across metabolic conditions. The RNA-Seq datasets generated during the sequencing procedure is deposited in the Gene Expression Omnibus database (access number GSE306485 and GSE306292), the mass spectrometry proteomics and peptidomics datasets have been deposited to the ProteomeXchange Consortium via the PRIDE partner repository (access numbers PXD076084) and available from the corresponding author upon request.

## Author Contributions

E.N.G. was responsible for the conceptualization and design of the study. M.B., W.S.-W. and E.H.G were responsible for the design and execution of most of the experiments. J.N., F.R.-C., A.L., T.L., V.M.N., M.L., I.P.-C., G.G.H., and S.D. performed experiments. C.E.B., V.V. and S.P.S. performed bioinformatic analyses. L.Y., D.O., and D.V. designed and performed mass proteomic analyses. A.B.R., M.D.F., C.M.M., B.R.-M., and N.M. provided human liver or islet samples, patient characterization and data analysis. D.E. and J.M. provided materials and designed the biosensor experiments. M.B. and P.G. provided the protocol, assistance and data interpretation for the perifusion islet analysis. V.S. and L.C. provided TEM analysis and data interpretation. N.B., AK.C., and H.J.S. provided assistance in experimental plan and data interpretation. E.N.G., M.B. and E.H.G. wrote the manuscript and interpreted the data with intellectual input and approval from all authors.

## Acknowledgements

We thank André Dias, Madalina Popa, Erick Arroba, Mariana Nunes, Anne Van Praet, Rabéa Dahili and Cláudia Pinto (Université libre de Bruxelles); Chloé Despontin, Gaëtan Herinckx, Firas Khattab (Université catholique de Louvain) for their experimental and technical support. We thank Latifa Bakiri (Medical University of Vienna), Sarah-Maria Fendt (KULeuven), and Alfredo Giménez-Cassina (Karolinska Institutet) for critical reading of the manuscript.

