## Supplementary Information for "PTPRF is a stress-responsive cytoskeletal checkpoint that coordinates metabolic adaptation in hepatocytes and β cells"

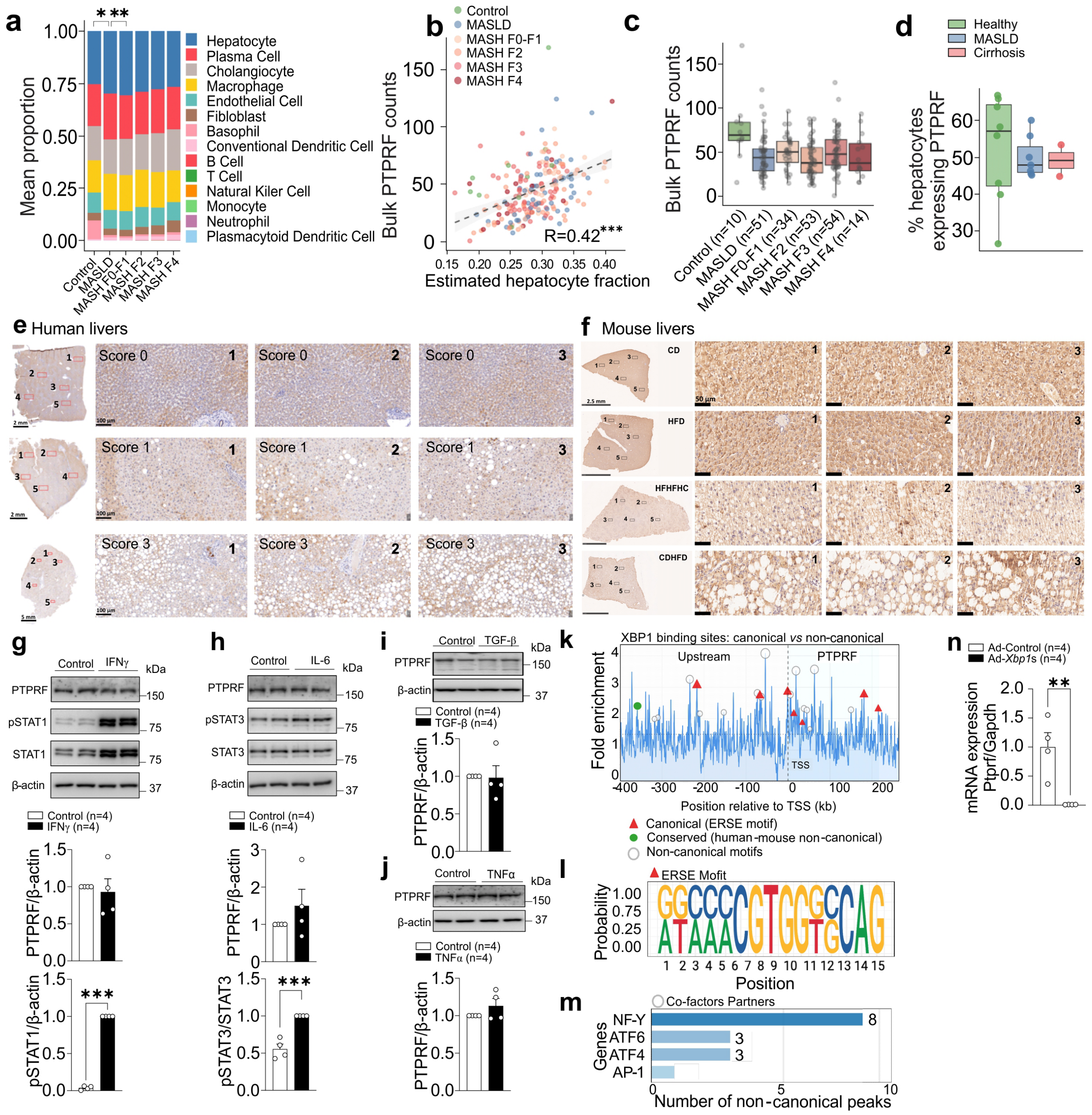


**Supplementary Figure 1. PTPRF downregulation in MASLD is hepatocyte-intrinsic and insensitive to cytokine exposure** **(a)** Estimated liver cell-type composition across MASLD stages, obtained by MuSiC deconvolution of bulk liver transcriptomes from GSE135251 (n =

216). Hepatocyte proportion significantly declines with disease progression (Kruskal–Wallis $p = 0.0065$), with pairwise comparisons showing reduced hepatocyte fraction in MAFLD ($p = 0.012$) and MASH F0–F1 ($P = 0.003$) relative to Control (Wilcoxon rank-sum test). (**b**) Association between bulk liver PTPRF expression and hepatocyte fraction across samples. Bulk PTPRF expression positively correlates with estimated hepatocyte abundance ($\beta = 219$, $P = 5.8 \times 10^{-10}$). (**c**) Bulk liver PTPRF expression across disease groups. (**d**) Detection rate of PTPRF across hepatocyte nuclei in each disease group. **(e)** Immunohistochemical detection of PTPRF in human liver biopsies from individuals with steatosis scores ranging from 0 to 3. Scale bars: 2.5 mm or 50 μm. **(f)** Immunohistochemistry of PTPRF in liver sections from 12-week dietary mouse models, high-fat diet (HFD), high-fat high-fructose high-cholesterol diet (HFHFHCD), choline-deficient high-fat diet (CDHFD) or control diet (CD) . Scale bars: 2mm, 5 mm or 100 μm. (**g–j**) Western blot analysis of PTPRF protein levels in primary mouse hepatocytes treated for 24 h with pro-inflammatory or pro-fibrotic cytokines: (**g**) IFNγ, (**h**) IL-6, (**i**) TGFβ and (**j**) TNFα treatments. (**k**) XBP1 ChIP-seq enrichment profile across the mouse *Ptprf* locus (GSE150889) showing 26 XBP1 binding sites spanning upstream regulatory regions to the gene body. Red triangles indicate canonical peaks containing ERSE motifs (CGTGG; n = 7, 26.9%), representing direct XBP1–DNA binding, including the primary enhancer at −54 kb (4.08× enrichment) and a promoter peak at −1 kb (2.85× enrichment). Grey circles denote non-canonical peaks lacking consensus ERSE motifs (n = 19, 73.1%), suggesting indirect XBP1 recruitment via co-factor partnerships. A green circle marks an evolutionarily conserved distal regulatory region at −359 kb (2.38× enrichment), identified through syntenic mapping to human ENCODE data (ENCSR988EVQ, −352.7 kb, 5.5% positional drift). Triangle size reflects ChIP-seq enrichment strength. Dashed line indicates the transcription start site (TSS); dotted line marks the significance threshold (1.5× enrichment). Grey shading indicates the upstream region; blue shading indicates the *Ptprf* gene body (216.7 kb). **(l)** Sequence logo of ERSE motifs (ER stress response elements) extracted from the 7 canonical XBP1 binding peaks. The conserved CGTGG core sequence (positions 6–10) represents the canonical XBP1 recognition motif, consistent with established UPR target genes. Flanking sequences show additional conservation, suggesting importance for binding specificity. The logo was generated from aligned 15-bp sequences centered on ERSE motifs using information content calculated from observed nucleotide frequencies. **(m)** Motif enrichment analysis of non-canonical XBP1 peaks revealing transcription factor binding motifs associated with UPR regulatory complexes, including NF-Y, ATF6, ATF4, and AP-1, suggesting co-factor–mediated recruitment of XBP1 to the *Ptprf* locus. **(n)** *Ptprf* mRNA levels in primary mouse hepatocytes following adenoviral

XBP1s overexpression. Bar graphs display quantification as mean ± SEM with individual data points. Statistical significance was determined using an unpaired t test for panels g, h, i, j and n, and using the Kruskal–Wallis test followed by pairwise Wilcoxon rank-sum tests for panel a. Asterisks denote significant pairwise comparisons; non-significant comparisons are not shown. *$P < 0.05$, **$P < 0.01$, ***$P < 0.001$.

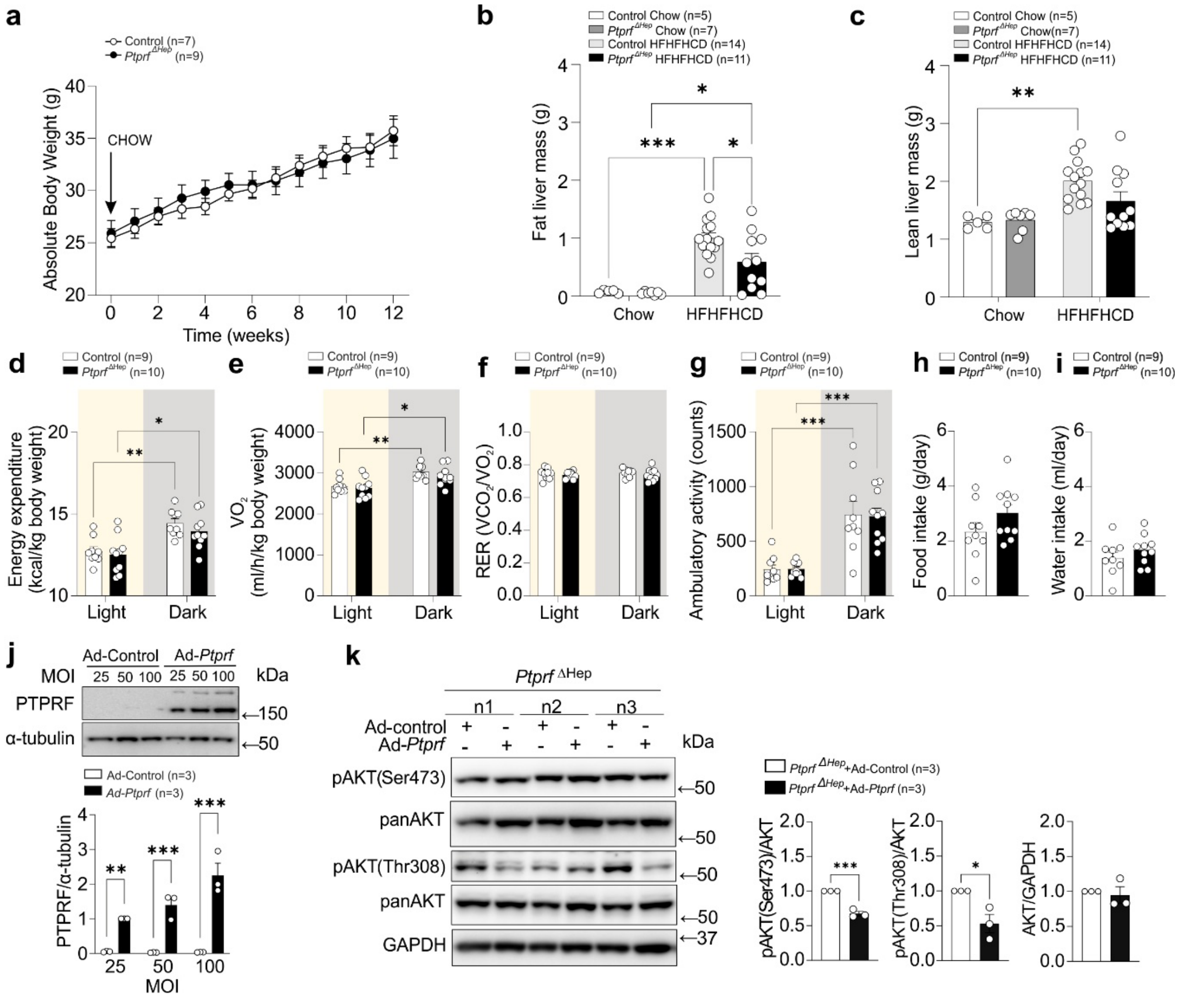


**Supplementary Figure 2. Metabolic characterization of *Ptprf*$^{\Delta Hep}$ and control mice under chow or HFHFHCD feeding. (a)** Body weight of *Ptprf*$^{\Delta Hep}$ and control mice after 12 weeks on chow diet. (**b**) Fat liver mass and (**c**) lean liver mass (in grams) in *Ptprf*$^{\Delta Hep}$ and control mice after the chow diet or HFHFHCD feeding. **(d)** Energy expenditure measured by indirect calorimetry in HFHFHCD fed *Ptprf*$^{\Delta Hep}$ and control mice. **(e)** Oxygen consumption rate ($VO_2$) under basal conditions in *Ptprf*$^{\Delta Hep}$ and control mice. **(f)** Respiratory exchange ratio (RER) in *Ptprf*$^{\Delta Hep}$ and control mice. **(g-i)** Ambulatory activity (g), food intake (h) and water intake (i) over a 24-h period in *Ptprf*$^{\Delta Hep}$ and control mice. (**j**) Western blot analysis of liver lysates derived from control mice 2 weeks after viral transduction with different MOIs as indicated. (**k**) Immunoblot analysis of *Ptprf*$^{\Delta Hep}$ primary mouse hepatocytes transduced with Ad-control or Ad-*Ptprf*. Bar graphs display quantification as mean ± SEM with individual data points. Statistical significance was determined using an unpaired t test (panels h-k) or Two-way ANOVA (panels b-g). *$P < 0.05$, **$P < 0.01$, ***$P < 0.001$.

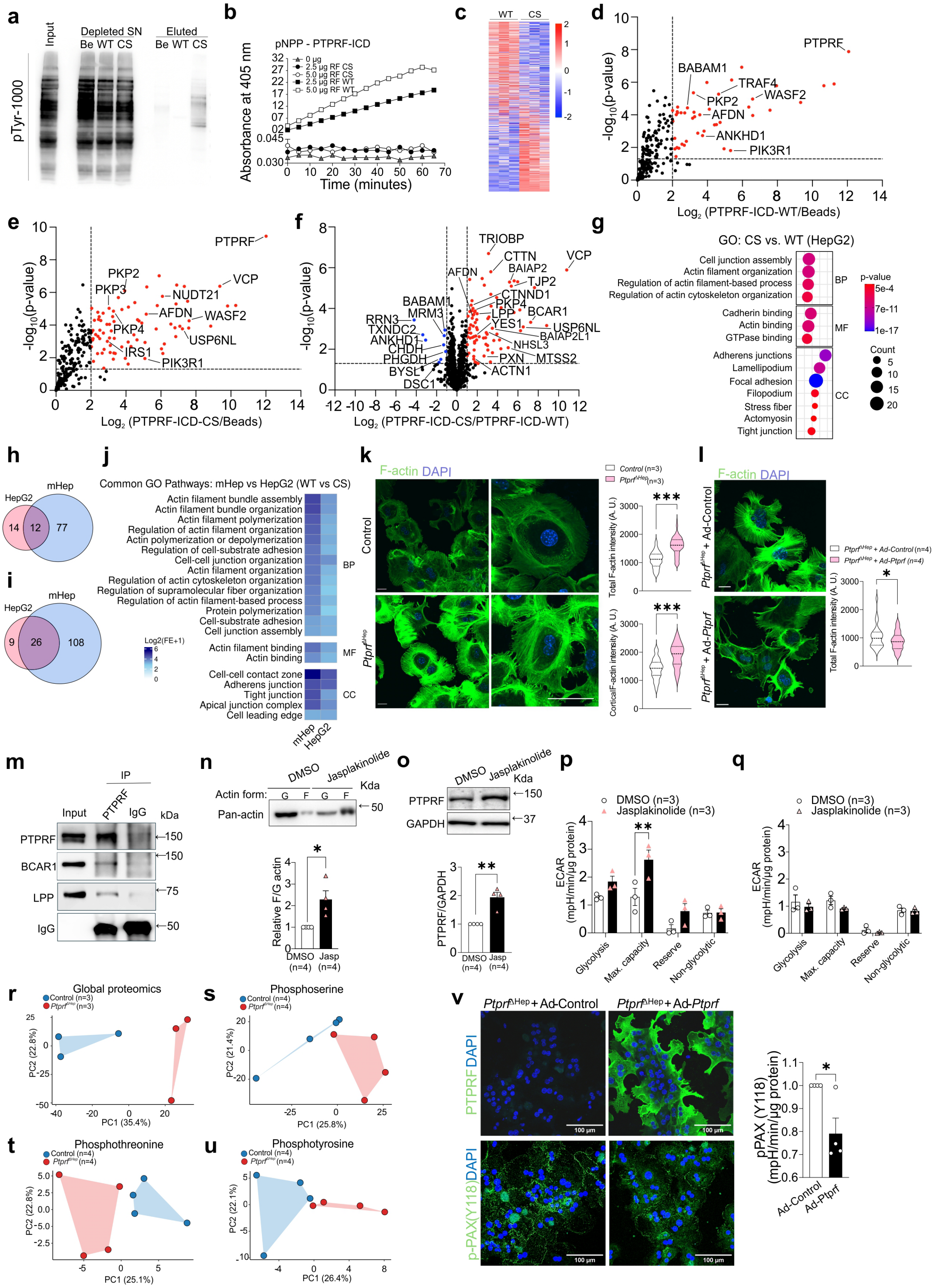


**Supplementary Figure 3. Extended interactome, functional cytoskeleton studies, and liver phosphoproteomics.** **(a)** Western blot for phosphotyrosine (pY) in eluates and depleted supernatants from pervanadate-treated primary mouse hepatocytes (mHep) lysates after pull-down with WT or CS mutant ICDs. **(b)** *In vitro* dephosphorylation assay confirming catalytic activity of PTPRF ICD WT and absence of catalysis with PTPRF ICD CS mutant **(c)** Heatmap of differentially enriched interactors in HepG2 cells lysates exposed to WT vs. CS PTPRF ICD. PCA of phosphothreonine profiles. **(d–f)** Volcano plots showing differentially enriched proteins in the PTPRF interactome of HepG2 cells. **(g)** Gene Ontology enrichment analysis of interactors identified in HepG2 cells. **(h)** Venn diagram showing overlapping enriched proteins is CS vs. WT PTPRF ICD between mHep and HepG2 interactome analysis. (**i**) Venn diagram showing overlapping enriched GO pathways identified in mHep and HepG2 interactome analyses. (**j**) Overlap of GO pathways (BP: Biological Pathways; MF: Molecular Function; CC: Cellular Components) between mHep and HepG2 cells. (**k**) Staining of F-actin (SPY-actin) in mHep from $Ptprf^{\Delta Hep}$ and control mice and quantification of the total intensity and the fluorescence intensity of the stress fibers the boarder of the cells (n=3, 108 cells/group). (**l**) Staining of F-actin (SPY-actin) in primary hepatocytes from $Ptprf^{\Delta Hep}$ mice transduced with Ad-*Ptprf* or Ad-control and F-actin quantification (n=4, 45 cells/group). (**m**) Co-immunoprecipitation of LPP and BCAR1 with PTPRF in HepG2 cells following AdV-mediated PTPRF overexpression. (**n**) Immunoblot analysis of F actin and G actin fractions following jasplakinolide treatment in mHep. (**o**) Immunoblot analysis of PTPRF protein levels in primary mouse hepatocytes following jasplakinolide treatment. (**p, q**) Glycolytic parameters derived from the glycolysis stress test in control mHep or (p) mHep from $Ptprf^{\Delta Hep}$ mice (q). **(r-u)** PCA of global proteome, phosphoserine, phosphothreonine and phosphotyrosine profiles shown in **Fig. 4j-q**. (**v**) Representative fluorescence images and quantification showing phospho paxillin (pPAX, Y118, green) and nuclei (blue, DAPI) in $Ptprf^{\Delta Hep}$ mHep transduced with Ad-*Ptprf* or Ad-control. Bar graphs display quantification as mean ± SEM with individual data points. Statistical significance was determined using an unpaired t test (panels k, l, m, n, o and u). $^{*}P < 0.05$, $^{**}P < 0.01$, $^{***}P < 0.001$.

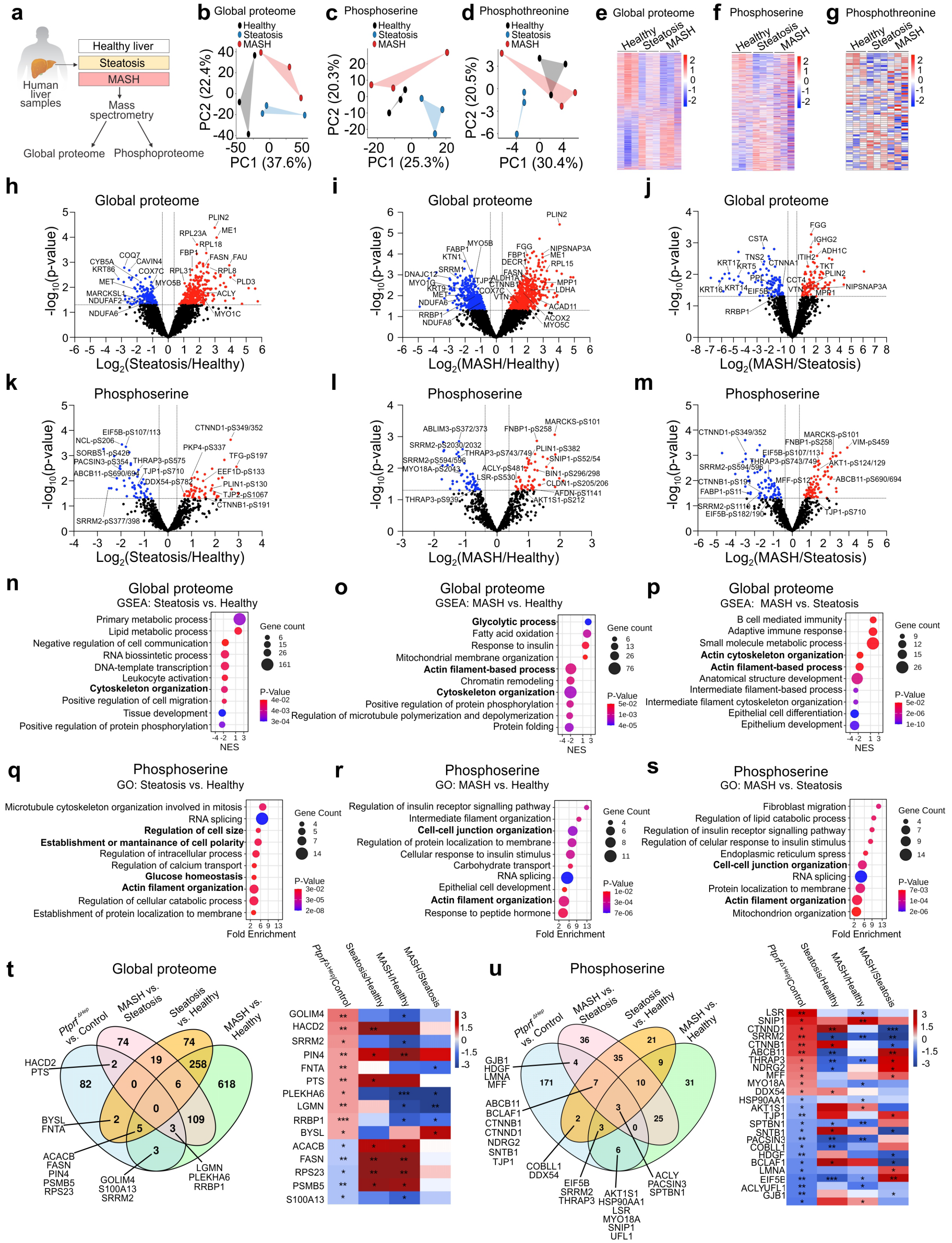


**Supplementary Figure 4. Integrated proteomic and phosphoproteomic analysis of human liver samples across MASLD progression.** (**a**) Schematic overview of the experimental workflow for global proteome and phosphoproteome profiling of human liver samples classified as healthy, steatosis, or MASH. (**b–d**) Principal component analysis of samples based on (b) global proteomics, (c) phosphoserine residues, and (d) phosphothreonine residues. (**e–g**)

Heatmaps showing hierarchical clustering of significantly altered proteins or phosphosites for (e) global proteome, (f) phosphoserine, and (g) phosphothreonine datasets. (**h–j**) Volcano plots depicting differential protein abundance in global proteomics comparisons of (h) steatosis versus healthy, (i) MASH versus healthy, and (j) MASH versus steatosis. (**k–m**) Volcano plots showing differentially phosphorylated serine residues for (k) steatosis versus healthy, (l) MASH versus healthy, and (m) MASH versus steatosis. (**n–p**) Gene set enrichment analysis of global proteomics data for (n) steatosis versus healthy, (o) MASH versus healthy, and (p) MASH versus steatosis. (**q–s**) Gene Ontology enrichment analysis of differentially phosphorylated serine residues for (q) steatosis versus healthy, (r) MASH versus healthy, and (s) MASH versus steatosis. (**t–u**) Integrative analysis between human liver proteomic datasets and murine $Ptprf^{\Delta Hep}$ versus Control liver proteomics. Venn diagrams display overlapping differentially expressed global proteins (t, left) and phosphoserine modified proteins (u, left), while heatmaps illustrate commonly regulated proteins and phosphosites across human and mouse datasets (t–u, right).

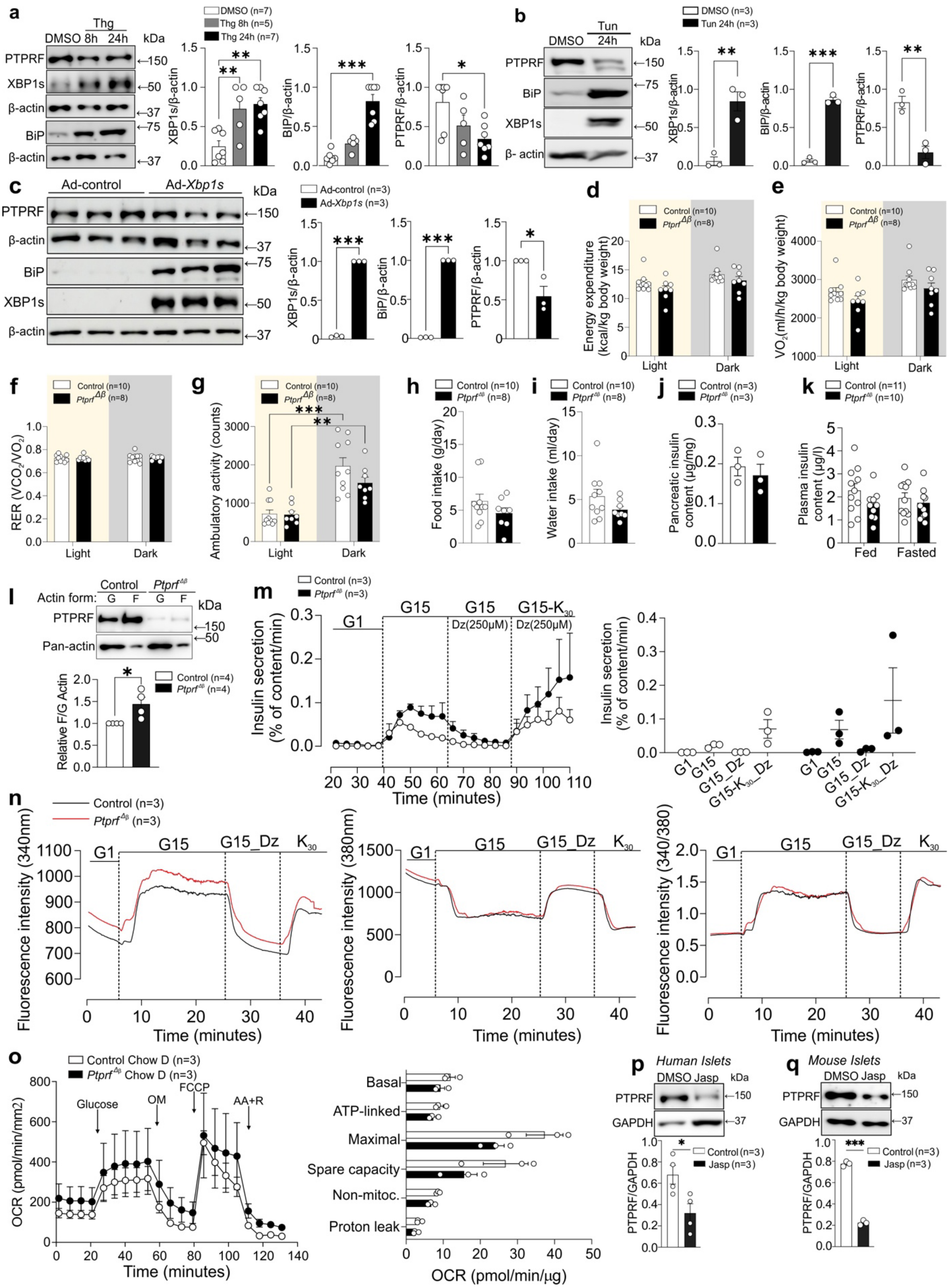


**Supplementary Figure 5. Extended metabolic, functional, and cytoskeletal characterization of human islets, mouse islets, or EndoC-βH1 cells. (a)** Immunoblot analysis of PTPRF protein expression in primary mouse islets following induction of ER stress with thapsigargin, alongside markers of unfolded protein response activation (BiP and spliced XBP1). **(b)** Immunoblot analysis of PTPRF protein expression in EndoC-βH1 following induction of ER stress with tunicamycin, alongside markers of unfolded protein response activation (BiP and spliced XBP1). **(c)** Immunoblot analysis of PTPRF protein expression in EndoC-βH1 following adenoviral-mediated overexpression of spliced XBP1 (XBP-1s), alongside markers of unfolded protein response activation (BiP and XBP1s). **(d-i)** Metabolic cage analyses assessing (d) energy expenditure, (e) oxygen consumption, (f) respiratory exchange ratio, (g) ambulatory activity, (h) food intake and (i) water intake in *Ptprf*$^{\Delta\beta}$ and control mice following HFHFHCD feeding for 12 weeks. **(j)** Quantification of pancreatic insulin content in *Ptprf*$^{\Delta\beta}$ and control mice following 12 weeks HFHFHCD feeding. **(k)** Circulating plasma insulin concentrations measured in fed and fasted states in *Ptprf*$^{\Delta\beta}$ and control mice following 12 weeks HFHFHCD feeding. **(l)** Immunoblot analysis of F and G actin fractions from primary mouse islets of *Ptprf*$^{\Delta\beta}$ and control mice on chow diet **(m)** Dynamic perifusion profiles of insulin secretion (4 min intervals) following sequential stimulation with low glucose (G1), high glucose (G15), diazoxide (Dz), and KCl ($K_{30}$) on isolated *Ptprf*$^{\Delta\beta}$ and control islets under Chow conditions. Scatter plot of insulin secretion across conditions. **(n)** Cytosolic $Ca^{2+}$ dynamics in *Ptprf*$^{\Delta\beta}$ and control islets following HFHFHCD feeding for 12 weeks **(o)** Mitochondrial respiration measurements in *Ptprf*$^{\Delta\beta}$ and control islets under chow conditions. Oxygen consumption rate trace from Seahorse mitochondrial stress test in primary mouse islets from *Ptprf*$^{\Delta\beta}$ and control mice and quantification of mitochondrial parameters for basal respiration, ATP-linked respiration, maximal respiration, and spare respiratory capacity. **(p)** Immunoblot analysis of PTPRF protein levels in primary human islets following jasplakinolide treatment. **(q)** Immunoblot analysis of PTPRF protein levels in primary mouse islets following jasplakinolide treatment In a-k, m, o-q results are shown as means ± SEM. Statistical analyses were done using two-tailed unpaired Student's t test (a-c, h-j, o-q), one-way ANOVA (d-g, k) or two-way repeated measures ANOVA plus Tukey's test (m) and denoted as $*P < 0.05$, $**P < 0.01$, $***P < 0.001$.

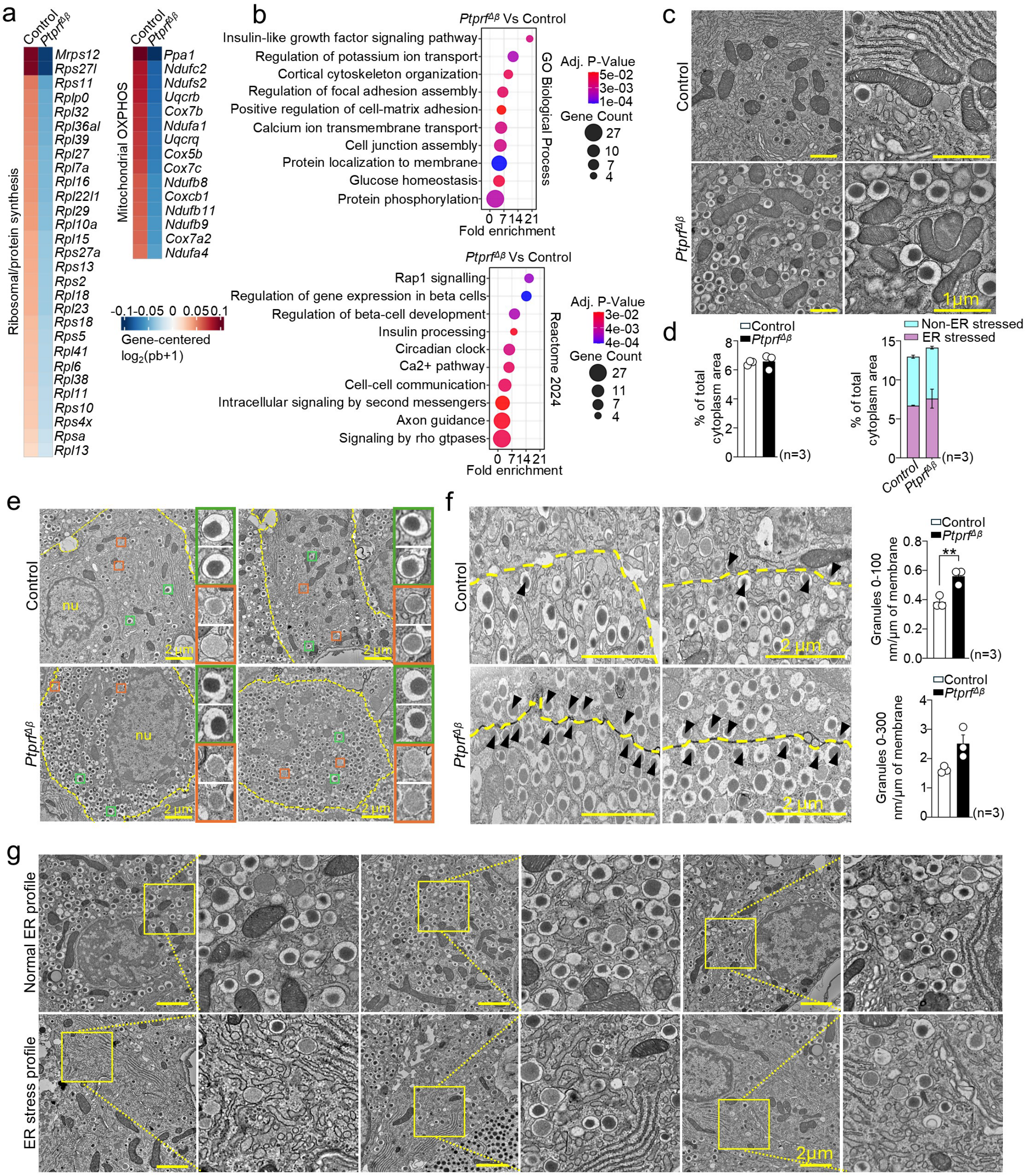


**Supplementary Figure 6. Extended single-cell transcriptomic and ultrastructural characterization of *Ptprf*$^{\Delta\beta}$ and control islets under nutrient stress. (a)** Expanded heatmap showing expression of representative ribosomal overload, and mitochondrial dysfunction genes significantly upregulated with a $\log_2$ fold change > 0.25 and adjusted $P < 0.05$ in control. **(b)** Additional gene ontology and Reactome pathway analyses of the genes significantly upregulated with a $\log_2$ fold change > 0.25 and adjusted $P < 0.05$ in *Ptprf*$^{\Delta\beta}$ showing enrichment of cytoskeletal, vesicle trafficking, and metabolic programs. **(c)** Representative TEM images of β cells from pancreatic islets of HFHFHCD-fed *Ptprf*$^{\Delta\beta}$ and control mice at high magnification, highlighting mitochondrial morphology and distribution relative to insulin granules. Scale bars:

1 µm **(d)** Quantification of the percentage of total cytoplasmic area occupied by mitochondria in $Ptprf^{\Delta\beta}$ and control β cells. **(e)** Representative TEM images of control and $Ptprf^{\Delta\beta}$ β cells at 10,000x magnification. Inset shows enlarged views of representative mature (green) or immature (red) granules. Nu, nucleus. Scale bars: 2 µm. **(f)** Representative TEM images and quantification of insulin granule distribution relative to the plasma membrane in β cells from control and $Ptprf^{\Delta\beta}$ islets. Granule density was quantified within 0–100 nm (cortical) and 100–300 nm (subcortical) regions and normalized to membrane length (granules per µm of membrane). Arrowheads indicate membrane-proximal granules. Scale bars: 2µm. **(g)** Representative TEM images illustrating ER stress, allowing classification of cells as having normal ER morphology or ER under stress. Inset shows enlarged views of representative ultrastructure. In d and f results are shown as means ± SEM. Statistical analyses were done using two-tailed unpaired t-test with Welch's correction (d and g). $^{**}P < 0.01$.

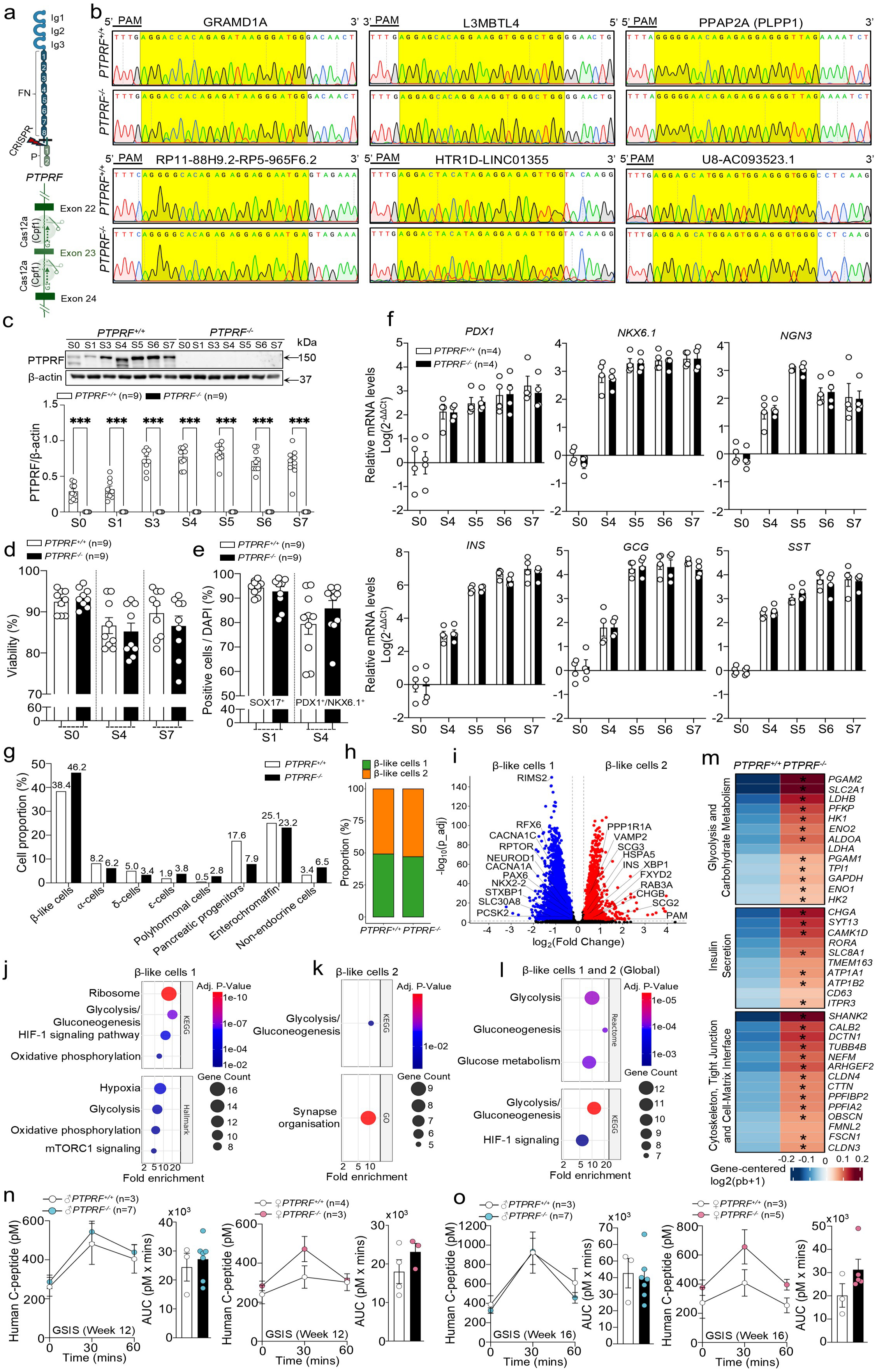


**Supplementary Figure 7. Genome integrity, differentiation efficiency and extended transcriptomic and *in vivo* characterization of *PTPRF*-deficient SC-islets. (a)** Schematic representation of the human *PTPRF* locus indicating extracellular, transmembrane and intracellular phosphatase domains and the Cas12a target site used for gene disruption. **(b)** Sanger sequencing chromatograms demonstrating unaltered DNA sequence at six predicted Cas12a off-target sites in *PTPRF*$^{-/-}$ and *PTPRF*$^{+/+}$ clones. **(c)** Western blot analysis and quantification of PTPRF protein expression during differentiation. **(d)** Viability of *PTPRF*$^{-/-}$ and *PTPRF*$^{+/+}$ cells at the pluripotent, pancreatic endoderm and SC-islet stages. **(e)** Quantification of SOX17$^{+}$ definitive endoderm (stage 1) and PDX1$^{+}$/NKX6.1$^{+}$ pancreatic progenitors (stage 4). **(f)** qPCR analysis of *PDX1*, *NKX6.1*, *NGN3*, *INS*, *GCG*, and *SST* mRNA levels across differentiation stages. **(g, h)** Cell-type composition of endocrine and non-endocrine populations in *PTPRF*$^{-/-}$ and *PTPRF*$^{+/+}$ SC-islets based on single-cell transcriptomics. **(i)** Volcano plot displaying differentially expressed genes between the β-like cell 1 and β-like cell 2 clusters in the integrated PTPRF dataset. **(j–l)** Pathway enrichment analyses showing upregulation of cytoskeletal, glycolytic and insulin secretion pathways in *PTPRF*$^{-/-}$ SC-islets in the β-like cell 1 cluster (j), β-like cell 2 cluster (k) and the merged β-like cell population (l). **(m)** Heatmap of representative genes driving enriched pathways in the merged β-like cell population. Genes marked with an asterisk are significantly upregulated in *PTPRF*$^{-/-}$ SC-islets ($\log_2$ fold change > 0.25 and adjusted $P < 0.05$). **(n, o)** Glucose-stimulated insulin secretion profiles and corresponding area-under-the-curve in male and female mice at 12- (n) and 16- (o) weeks post-implantation of *PTPRF*$^{-/-}$ or *PTPRF*$^{+/+}$ SC-islets. In c-f, n, o results are shown as means ± SEM. Statistical analyses were done using two-tailed unpaired t-test (c-e, n, o) or two-way ANOVA (f) and denoted as $***P < 0.001$.

### Supplementary Material and Methods

#### Human liver samples

Inclusion and exclusion criteria for liver samples obtained from patients with obesity who underwent bariatric surgery at the Virgen de la Arrixaca Clinical University Hospital (Murcia, Spain) have been described elsewhere [1, 2].

#### Antibodies, reagents, recombinant proteins and plasmid constructs

Primary and secondary antibodies used in this study are listed in **Supplementary Table 5** with application and dilution information.

For *in vitro* experiments, sodium pyruvate (#103578 100), glutamine (#103579 100) and glucose (#103577 100) were purchased from Agilent. Human insulin solution (#I9278 5ML), sodium palmitate (#P9767 5G), oleic acid (#O7501 1G), thapsigargin (#T9033 .5MG), cyclopiazonic acid (#C1530 5MG), tunicamycin (#T7765 5MG) and bovine serum albumin (#A8022 100G) were obtained from Sigma Aldrich. Recombinant human IL-6 (#206 IL 010, R and D Systems), recombinant human TGF-β (#100 21C 10 μg, PeproTech), recombinant mouse TNFα (#410 MT 025, R&D Systems) and recombinant murine IFNγ (#315 05 100 μg, PeproTech) were used as indicated.

For *in vivo* experiments, D-glucose (#1083421000, Sigma Aldrich), intraperitoneally injected insulin ProZinc (NDC #0010 4499 01, Boehringer Ingelheim, Rhein, Germany), or intravenously injected Actrapid human insulin, 100 IU per ml (Novo Nordisk) were used.

Full length intracellular domain $His_6$ tagged wild type and C1548S substrate trapping mutant human PTPRF (cysteine 1548 in the D1 domain replaced with serine 1548) were kindly provided by Dr Hayley Sharpe, Babraham Institute, Cambridge, United Kingdom.

For full length PTPRF overexpression, the complete coding sequence of human PTPRF (P10586) was fused to mGreenLantern. Plasmids were synthesized by GenScript and cloned into the pcDNA3.1 vector containing the simian cytomegalovirus major immediate early promoter IE94.

#### *In vitro* culture of primary human islets and EndoC-βH1 cells

Primary human pancreatic islets were provided by the Network for Islet Transplantation (Université Catholique de Louvain). Upon arrival, islets were cultured in Ham's F-10 Nutrient Mixture (Gibco, 41550), supplemented with 1% bovine serum albumin, 2 mM GlutaMAX, 1% penicillin–streptomycin, and 10% fetal bovine serum. IBMX was added at a final concentration of 50 μM. For treatments, islets were cultured in 1% fetal bovine serum. Islets were maintained on ultra-low attachment culture plates at 37°C in a humidified atmosphere containing 5% $CO_2$. Donor characteristics are provided in **Supplementary Table 3**.

The human β-cell line EndoC-βH1 (kindly provided by INNODIA consortium and Prof R. Scharfmann, University of Paris, France) was cultured in Matrigel-fibronectin-coated plates [3].

#### Mice

Animals were housed at 22 °C under a 12-h light and 12-h dark cycle with *ad libitum* access to food and water. C57BL/6N mice were used as non-transgenic controls. A hepatocyte specific (*Ptprf*$^{\Delta Hep}$) and β-cell-specific (*Ptprf*$^{\Delta\beta}$) conditional *Ptprf* knockout lines were generated inhouse on a C57BL/6N background.

*Ptprf* floxed mice (Cyagen Biosciences) were engineered by flanking exons 8 to 10 of *Ptprf*, NCBI Reference Sequence NM_011213.2, with loxP sites via homologous recombination in C57BL/6N embryonic stem cells. These mice were crossed with *Alb-Cre* transgenic mice (The Jackson Laboratory, B6.Cg-*Speer6-ps1*$^{Tg(Alb\text{-}cre)21Mgn}$/J; JAX #003574) or *Ins1-Cre* (The Jackson Laboratory, B6(Cg)-*Ins1*$^{tm1.1(cre)Thor}$/J; JAX #026801) transgenic mice, which expresses Cre recombinase under the albumin or insulin promoter respectively. The Cre expression does not impact on β cells/hepatocyte toxicity or function [4] and data not shown. Both parental lines were homozygous for the *Ptprf*-floxed allele, and only one parent carried the *Alb-Cre* or *Ins1-Cre* transgene in a heterozygous state, to restrict recombination to hepatocytes or β cells. Offspring carrying *Alb-Cre* were classified as *Ptprf*$^{\Delta Hep}$ mice, *Ins1-Cre* classified as *Ptprf*$^{\Delta\beta}$ and littermates *Ptprf*$^{lox/lox}$ served as controls.

Eight-week-old C57BL/6N, *Ptprf*$^{\Delta Hep}$ or *Ptprf*$^{\Delta\beta}$, and littermate controls were randomly assigned to diet intervention studies. Experimental diets from Research Diets, New Brunswick, New Jersey, United States, included: high fat diet, HFD; 60 kcal % fat, D12492; high fat, high fructose, high cholesterol diet, HFHFHCD; 40 kcal % fat, 20 kcal % fructose, 2% cholesterol, D09100310i; methionine and choline deficient high fat diet, HFMCD; 45 kcal % fat, 0.1% methionine, no added choline, A20012301i. Reference diets included a control diet with 10 kcal % fat, D09100304i, or a chow low fat diet with 3 kcal % fat, RM1 (P) 801151, Special Diets Services, United Kingdom. LFD was used for general maintenance and non-obesity experiments. Diet exposure ranged from 2 to 40 weeks.

All animal experiments complied with Belgian regulations for animal care and were approved by the Commission d'Éthique du Bien Être Animal at the Faculty of Medicine, Université libre de Bruxelles, dossiers 732N, 917N and 918N.

**Metabolic phenotyping**

Whole body and liver lean and fat mass were quantified using an EchoMRI body composition analyzer, EchoMedical Systems, Houston, Texas, United States. Body weight was measured every week or every two weeks as indicated. Glucose tolerance tests were performed after 6 h of fasting. Mice received intraperitoneal injections of D-glucose at 2g per kg body weight. Insulin tolerance tests were performed after 4 h of fasting. Mice received intraperitoneal injections of insulin at 0.75 U per kg body weight. Solutions were prepared in phosphate buffered saline immediately before injection. Tail tip blood glucose was measured with an Accu Chek Performa glucometer, Roche, Basel, Switzerland.

*Ptprf*$^{\Delta Hep}$ mice and *Ptprf*$^{\Delta\beta}$ mice that had been fed HFHFHCD for 10 and 12 weeks respectively along with their respective controls were placed in TSE PhenoMaster metabolic cages, TSE, Germany, for 72 h. After a 24 h acclimatization period, indirect calorimetry was performed to measure energy expenditure, oxygen consumption, respiratory exchange ratio, ambulatory activity and food and water intake.

**Primary mouse hepatocyte isolation, cell culture and treatments**

Primary mouse hepatocytes were isolated from *Ptprf*$^{\Delta Hep}$ or control mice following overnight *ad libitum* feeding using a two-step collagenase perfusion through the vena cava. Mice were anaesthetized by intraperitoneal injection of ketamine (100 mg $kg^{-1}$) and xylazine (10 mg $kg^{-1}$), the peritoneum was opened, and the infra-hepatic vena cava was cannulated for perfusion. The portal vein was cut to allow blood outflow from the liver. In the first perfusion step, the liver was perfused for 10 min at 37 °C with calcium-free Hanks' balanced salt solution (HBSS) supplemented with 10 mM HEPES (pH 7.4) and equilibrated with 95% $O_2$ and 5% $CO_2$ (vol/vol). In the second step, collagenase type IV (0.3 mg $ml^{-1}$) in William's E medium was perfused for an additional 10 min to digest the liver tissue. The digested liver was transferred to a sterile plastic dish, gently dissociated in cold William's E medium and filtered through a 75-µm cell strainer to remove clumps. The resulting cell suspension was pelleted by centrifugation at 50 × g for 5 min at 4 °C, resuspended in William's E medium and purified by Percoll® centrifugation to remove dead cells. Hepatocytes were subsequently washed three times in William's E medium (50 × g for 5 min at 4 °C). Viability assessment by Trypan blue exclusion typically yielded 2–4 × $10^7$ cells per preparation with approximately 85% viability. Each experimental data point corresponds to an independent hepatocyte preparation.

For high-fat and low-fat hepatocytes were isolated from HFHFHCD fed mice, as described [5]. Briefly, after isolation the cell suspension was pelleted by centrifugation at 50 × g for 5 min at 4° C once and sequential Percoll® fractionation was performed, allowing the separation of hepatocytes in populations with distinct lipid content, including high-fat and low-fat hepatocytes. These hepatocytes were pelleted and stored at -80° C for protein extraction and proteomic/phosphoproteomic analysis or cultured for 24 h before fixation for lipid staining.

For cell culture, hepatocytes used for immunoblotting were seeded in uncoated 24-well plastic plates at 1 × $10^5$ cells per well in 500 µl of attachment medium consisting of William's E medium with GlutaMAX, 10% FBS, 1% penicillin–streptomycin and 10 mM HEPES. For immunofluorescence and lipid staining, hepatocytes were seeded in polymer-coated Ibidi plates at 5 × $10^4$ cells per well in 250 µl of attachment medium. Cells were allowed to attach for 4 h, after which the medium was replaced with maintenance medium (William's E medium with GlutaMAX, 10% FBS, 1% penicillin–streptomycin, 1% non-essential amino acids, 10 mM HEPES. Unless otherwise indicated, treatments were initiated after overnight culture.

For fatty-acid stimulation, hepatocytes were incubated in serum-free maintenance medium supplemented with 1% BSA containing palmitate (PA, 500 µM), oleate (OA, 500 µM), or PA plus OA (500 µM each). In selected experiments, cells were treated with PA alone (500 µM) under identical conditions. For ER-stress induction, hepatocytes were treated with thapsigargin (1 µM, 24 h), cyclopiazonic acid (75 µM, 6 h), or tunicamycin (5 µg $ml^{-1}$, 6 h). For cytokine stimulation, hepatocytes were treated for 24 h with IFNγ (50 U $ml^{-1}$), IL-6 (100 U $ml^{-1}$), TGF-β (5 ng $ml^{-1}$), or TNFα (100 U $ml^{-1}$). For insulin pulse–chase experiments, hepatocytes were serum-starved for 2 h in medium without FBS and then stimulated with insulin (100 nM) for 10 minutes. Cells were subsequently washed twice with PBS and incubated in insulin-free medium for the chase period. Control samples underwent identical medium changes without insulin. In experiments involving lipid loading and cytoskeletal modulation, hepatocytes were treated with palmitate (400 µM) plus oleate (800 µM) in the presence of 1% BSA, with or without jasplakinolide (1 µM), for 24 h. When lipid accumulation was induced by insulin (100 nM) and glucose (25 mM), cells were collected after 48 h of treatment. For time-in-culture experiments, hepatocytes were harvested 24, 48 or 72 h after plating.

For immunoblot analysis, following treatments, hepatocytes were washed twice with ice-cold PBS and lysed on ice in cell lysis buffer (Cell Signaling, #9803) supplemented with Halt protease and phosphatase inhibitors (Thermo Fisher Scientific, 78446). Lysates were clarified by centrifugation at 15,000 × g for 15 min at 4 °C, and supernatants were collected for downstream immunoblot analysis.

**Primary mouse islet isolation**

Mouse pancreatic islets were isolated by collagenase digestion. Briefly, pancreata were perfused and digested with collagenase type V (Sigma-Aldrich, C9263) prepared in serum-free M199 medium (Gibco, 22350) at a final concentration of 1 mg $ml^{-1}$. Following digestion, islets were purified by repeated hand-picking under a stereomicroscope until a highly enriched islet population was obtained.

**Islet dissociation and fluorescence-activated cell sorting for single-cell RNA sequencing**

For single-cell RNA sequencing experiments, islets were isolated from $Ptprf^{\Delta\beta}$ mice and control littermates after 12 weeks of HFHFHC diet feeding. Isolated islets were washed in cell dissociation buffer (124 mM NaCl, 5.4 mM KCl, 0.8 mM $MgSO_4$, 1 mM $Na_2PO_4$, and 10 mM HEPES). Islets were dispersed into single cells by gentle trituration in trypsin (1 mg $ml^{-1}$; Sigma-Aldrich, T9935) and DNase I (1 mg $ml^{-1}$; Sigma-Aldrich, 10104159001) at 37 °C. Dispersed cells were centrifuged and resuspended in FACS buffer, then passed through 35 µm nylon mesh filter-cap tubes (Fisher Scientific, 10100151) to remove cell aggregates. Live cells were identified by exclusion of propidium iodide (Sigma-Aldrich; 1 µg $ml^{-1}$) and sorted by flow cytometry. Dead cells and doublets were excluded by forward- and side-scatter gating. Sorted cells were filtered through 40 µm cell strainers (Corning) to remove residual clusters, pooled per experimental condition, and counted using a Luna automated cell counter. Only samples with >85% viability were processed for single-cell library preparation. Dead cells and doublets were additionally excluded during computational quality control prior to downstream transcriptomic analysis.

**Viral transduction**

For hepatic PTPRF overexpression *in vivo*, C57BL/6N mice received tail vein injections of $1 \times 10^9$ PFU Ad-*Ptprf* (Vector Biolabs, SKU ADV-220343) in 100 µl PBS. Control mice received $1 \times 10^9$ PFU Ad CMV Null, #1300, Vector Biolabs in 100 µl PBS.

For *in vitro* adenoviral experiments, hepatocytes were exposed 4 h after seeding to medium containing Ad-Control (Vector Biolabs), Ad-*Ptprf* (Vector Biolabs, SKU ADV-220343) or Ad-*Xbp1s* [6]. Unless otherwise indicated, adenoviral transduction was performed at MOI 50. Virus-containing medium was maintained for 24 h, after which it was replaced with fresh maintenance medium. Cells were analyzed 48 h after infection or subjected to subsequent treatments as indicated.

For *in vitro* transduction, EndoC-βH1 were infected with Ad-*Ptprf,* Ad-*Xbp1ss* or Ad-Control (MOI: 50), in DMEM GlutaMAX. Cells were used for experiments 48 h after transduction.

**G-actin/F-actin fractionation assay**

G-actin and F-actin fractions were isolated using the G-actin/F-actin In Vivo Assay Biochem Kit (Cytoskeleton, BK037) according to the manufacturer's instructions. Briefly, $2 \times 10^5$ primary hepatocytes, 2–3 × $10^5$ H1 hESCs, 250 isolated mouse islets, or 100 mg mouse liver tissue were lysed under actin-stabilizing conditions and separated by ultracentrifugation into G-actin-containing supernatant and F-actin-containing pellet fractions at 37 °C. The pellet was resuspended in depolymerization buffer, and both fractions were analyzed by western blotting. The F/G-actin ratio was calculated by densitometric quantification of actin immunoreactivity in the two fractions.

**Western blotting**

Tissue lysates were prepared in RIPA buffer (Cell Signaling, #9806) supplemented with Halt protease and phosphatase inhibitors (Thermo Fisher Scientific, 78446) and centrifuged at 21,130 g for 10 min. For lipid rich liver samples, lysates were centrifuged at 50,000 g for 15 min at 4 °C and the infranatant was collected. Lysates from cells were prepared as described above.

Protein concentration was determined using the Pierce BCA kit (Thermo Scientific, 23225). Lysates were mixed with five times concentrated sample buffer, 1.0 M Tris HCl pH 8.8, 0.5% bromophenol blue, 43.5% glycerol, 10% SDS, 1.3% β-mercaptoethanol, boiled at 98 °C for 10 min and separated by SDS PAGE, 8 to 12% gels. Proteins were transferred to 0.22 μm nitrocellulose membranes, blocked for 1 h in 5% milk in TBS (20 mM Tris-HCl, pH 7.4, 137 mM NaCl) with 0.3% Tween 20, and incubated overnight at 4 °C with primary antibodies in TBS with 5% BSA and 0.2% sodium azide. After three washes, membranes were incubated with HRP-conjugated secondary antibodies for 1 h and developed using SuperSignal West Pico or West Femto (Thermo Fisher Scientific) reagents on an Amersham ImageQuant 800 system. β-actin, GAPDH and α-tubulin were used as loading controls. The list of antibodies is described in in **Supplementary Table 5.**

***In vitro* lipid, F-actin and membrane staining**

Nile Red (5 μg ml$^{-1}$) was used to visualize lipid droplets, CellBrite Fix555 to label cell membranes, SPY555-actin (Spirochrome, SC202) to visualize F-actin, and Hoechst or DAPI for nuclear staining, as indicated, following the manufacturer’s instructions.

**Primary mouse islet insulin and F-actin staining**

Primary mouse islets were isolated from *Ptprf*$^{\Delta\beta}$ mice and control littermates (~12 weeks). Isolated islets were washed in cell dissociation buffer (124 mM NaCl, 5.4 mM KCl, 0.8 mM $MgSO_4$, 1 mM $Na_2PO_4$, and 10 mM HEPES). Islets were dispersed into single cells by gentle trituration in trypsin (1 mg ml$^{-1}$; Sigma-Aldrich, T9935) and DNase I (1 mg ml$^{-1}$; Sigma-Aldrich, 10104159001) at 37 °C. Dispersed cells were centrifuged and 25,000 cells were seeded in μ Slide 8 Well Ibidi plates in RPMI-1640 medium (Gibco, 11875093) with 10% FBS and 50 μM IBMX. Next day, cells were stimulated with low glucose (2 mM) or high glucose (20 mM) after 1 h starvation in Krebs–Ringer buffer containing 2 mM glucose. Cells were fixed in 4% PFA at indicated time points and then stained for Insulin to visualize β cells, SPY555-actin (Spirochrome, SC202) to visualize F-actin, and Hoechst or DAPI for nuclear staining, as indicated, following the manufacturer’s instructions.

**Measurement of fructose 1,6 bisphosphate using HYlight**

Fructose 1,6-bisphosphate was measured with the HYlight biosensor. 50,000 mHep were seeded in μ Slide 8 Well Ibidi plates and transfected 2 h later with 1 μg pCS2 plus HYlight plasmid using Lipofectamine 3000 in William’s medium with GlutaMAX (+ 1% non-essential amino acids, 10 mM HEPES, 5 μM hydrocortisone, 10% FBS, no antibiotics). 12 h after transfection, medium was changed to William's medium with GlutaMAX (+ 1% non-essential amino acids, 10 mM HEPES, 5 µM hydrocortisone, 10% FBS, and 1% penicillin-streptomycin).

On the day of imaging, cells were incubated for 1 h in XF assay medium without glucose. Live imaging was performed at 37 °C on a Nikon AX confocal microscope with a 20X objective and perfect focus system (PFS). HYlight was excited at 488 and 405 nm, and emission was collected through a 525/25 nm emission filter. The $F_{488/F405}$ excitation ratio was quantified over time in manually defined regions of interest using NIS Elements software. Further image processing was carried out using FIJI.

GraphPad Prism software 10.2.0 was used for statistical analysis. Based on the obtained $F_{488/F405}$ values, the area under the curve (AUC) was obtained for each condition.

**Oxygen consumption rate**

Oxygen consumption rate was measured with an XF Mito stress test kits (Agilent, #103010-100, #103016-100) using Seahorse XF HS Mini or Seahorse XFe24 Flex Analyzer. Primary hepatocytes were seeded at 10,000 cells per well in XFp plates. After attachment and adenoviral transduction, cells were incubated for 1 h at 37 °C without $CO_2$ in XF base medium supplemented with 2 mM glutamine, 10 mM glucose and 1 mM pyruvate. The mitochondrial stress test consisted of sequential injections of 6.3 µM oligomycin (OM), 1 µM FCCP, and 2.5 µM each of rotenone & antimycin A (AA+R). At the end of the assay, cells were lysed and stored at −20 °C for protein quantification and immunoblotting.

For palmitate–supported respiration experiments the Seahorse XFe24 Flex Analyzer was used. Primary hepatocytes were seeded at 25,000 cells per well in V7 microplates (Agilent, #100777-004). After viral transfection, cells were incubated overnight in substrate-limited medium consisting of Williams lacking glucose, glutamine, pyruvate, and HEPES, supplemented with 0.5 mM glucose, 1 mM GlutaMAX, 0.5 mM L-carnitine, 1% FBS and 1% penicillin-streptomycin. On the day of the assay, cells were equilibrated in Krebs–Henseleit–based assay medium supplemented with 125 µM palmitate-BSA, 2.5 mM glucose, 0.5 mM L-carnitine, and 5 mM HEPES (pH 7.4). Palmitate–BSA conjugate was prepared in-house by complexing sodium palmitate with fatty acid–free bovine serum albumin at a 6:1 molar ratio and added directly to wells 1 h prior to OCR measurements.

Isolated primary mouse islets were incubated for 1 h at 37 °C without $CO_2$ in XF base medium supplemented with 2 mM glutamine, 1 mM pyruvate and 2 mM glucose. The mitochondrial stress test consisted of sequential injections of 20 mM glucose, 5 µM oligomycin, 3 µM FCCP, 3 µM each of rotenone & antimycin A. For normalization, the cross-sectional area of the islets was calculated from the images taken by EVOS XL Core imaging system (Invitrogen) with a 4X objective lens. The digital images were analyzed using ImageJ software (NIH) and the threshold function was used to quantify the area of the islets, after setting up the threshold of the program to include all the islets in the image.

**Extracellular acidification rate**

Glycolytic activity was measured by extracellular acidification rate using the XF Glycolysis stress test kit (Agilent, #103017-100) using Seahorse XF HS Mini. Primary hepatocytes were seeded at 10,000 cells per well. After attachment and adenoviral transduction, cells were incubated for 1 h at 37 °C without $CO_2$ in XF base medium without glucose, supplemented with 2 mM glutamine.

The glycolytic stress test consisted of sequential injections of glucose, 10 mM, oligomycin (OM), 10 µM, and 2 deoxy D-glucose (2-DG) at a total of 100 mM delivered as two injections of 50 mM. After the assay, cells were lysed and stored at −20 °C.

**Transmission electron microscopy**

For liver, $Ptprf^{\Delta Hep}$ and control mice fed HFHFHCD for 2 weeks were perfused through the inferior vena cava with 2% paraformaldehyde and 2% glutaraldehyde in PBS. A 5 $mm^3$ block from the center of the left lobe was dissected and further fixed in the same solution for 2 h. For pancreas, $Ptprf^{\Delta Hep}$ and control mice fed HFHFHCD for 12 weeks were perfused through the common bile duct inflating the pancreas with 2% paraformaldehyde and 2% glutaraldehyde in PBS. A 5 $mm^3$region from the tail of the perfused pancreas was dissected and further fixed in the same solution for 2 h.

Samples were then further dissected to reach 1 $mm^3$ blocks and additionally fixed in 2.5% glutaraldehyde in 0.1 M sodium cacodylate buffer, washed, and post-fixed in 1% osmium tetroxide plus 1.5% ferrocyanide for 1 h, followed by an additional 1% osmium tetroxide step for 1 h and 1% uranyl acetate step for 2 h. After dehydration in graded ethanol and replacement with propylene oxide, samples were infiltrated and embedded in AGAR100 epoxy resin and cured at 60 °C for 72 h.

Semi-thin (1 µm) sections were performed to identify the regions of interest, then the ultrathin sections, 50 to 70 nm, were cut with a Leica UC6 ultramicrotome, recovered on 100 mesh formvar carbon coated copper grids and stained with 0.026% lead citrate. Images were acquired on a Tecnai 10 microscope at 100 kV with a Veleta camera.

Images were quantified using QPath software and manual delimitation of the structures on 60 to 80 cells per condition.

**Confocal imaging**

Confocal imaging for Nile Red, plasma membrane, full length PTPRF and nuclei was performed on a Zeiss LSM780 with the following channels: Nile Red, excitation 543 nm; CellBrite Fix555, excitation 555 nm; Phalloidin iFluor 633, excitation 633 nm; mGreenLantern, excitation 488 nm; DAPI, excitation 405 nm. Images were analyzed in Zeiss Zen software and mean fluorescence area for Nile Red and DAPI was quantified.

SPY555 F-actin and Hoechst were imaged on a Nikon Eclipse Ti2 inverted microscope with a Nikon AX R confocal system and 20-times objective (CFI Plan Apochromat Lambda NA:0.8).

**Interactome analysis**

*Pervanadate treatment*

$2 \times 10^6$ HepG2 cells in 10 cm dish, and 650,000 in 6 well plate, were cultured for 24 h and stimulated for 30 min at 37 °C and 5% $CO_2$ with complete medium containing 1 mM freshly prepared sodium pervanadate. Cells were then washed with ice cold PBS and lysed in 600 µl lysis buffer containing 50 mM Tris HCl pH 7.5, 150 mM NaCl, 10% glycerol, 1% Triton X 100, 1 mM EDTA, 5 mM iodoacetamide, 1 mM $Na_3VO_4$, 10 mM NaF and protease inhibitors. Lysates were incubated at 4 °C, treated with 10 mM DTT for 15 min, clarified at 14,000 g and snap frozen.

Sodium pervanadate was prepared by mixing 5 µl 3% $H_2O_2$ with 45 µl 20 mM HEPES pH 7.3, then adding 490 µl 100 mM $Na_3VO_4$ and 440 µl water. After 5 min at room temperature, catalase was added to quench $H_2O_2$ and pervanadate was used within 5 min.

*pNPP activity assay*

Phosphatase activity toward p nitrophenyl phosphate was measured in phosphate free buffer with 50 mM HEPES pH 7.4, 150 mM NaCl, 5% glycerol and 5 mM DTT. Recombinant phosphatases were mixed with pNPP, 20 mM, in 96 well plates and absorbance at 405 nm was measured over 70 min on a SpectraMax M5 reader (Molecular Devices).

*Streptavidin gel shift assay*

Biotinylation of AviTag proteins was verified by streptavidin shift. Biotinylated proteins were mixed with SDS sample buffer, heated, cooled and incubated with streptavidin in PBS. Samples were resolved on NuPAGE 4 to 12% gels, stained with InstantBlue and imaged on a ChemiDoc MP system. Band shifts were quantified by densitometry in Fiji.

*Recombinant protein pull downs*

Biotinylated His.TEV.Avi PTPRF ICD domains, 50 µg, were bound to 167 µl streptavidin magnetic beads, 4 mg per ml, in size exclusion buffer, 50 mM HEPES pH 7.5, 150 mM NaCl, 5% glycerol, 5 mM DTT, for 1 to 2 h at 4 °C. Beads were blocked with 5% BSA, and pervanadate treated lysates were precleared. PTPRF coated beads were incubated with 1 ml lysate, 1 mg per ml, for 1.5 h at 4 °C. Beads were washed with 150 and 500 mM NaCl wash buffers and TBS (20 mM Tris-HCl, pH 7.4, 137 mM NaCl).

For immunoblotting, beads were eluted in formamide EDTA and sample buffer containing biotin, heated and analyzed by SDS PAGE. For mass spectrometry, beads were washed and processed by on bead tryptic digest.

*On bead digestion*

Beads were resuspended in 50 mM ammonium bicarbonate pH 8.0, reduced with DTT, alkylated with iodoacetamide, quenched with DTT and digested sequentially with LysC and trypsin at 37 °C. Peptides were acidified, cleared, desalted using Sep Pak C18, eluted in 50% acetonitrile 0.1% TFA, dried and stored at −20 °C before LC-MS.

**LC-MS analysis**

*Phosphotyrosine enrichment*

Approximately $1 \times 10^7$ freshly isolated primary hepatocytes per mouse were lysed in 8 M urea HEPES buffer containing phosphatase inhibitors, reduced with DTT, alkylated with chloroacetamide and digested overnight with trypsin after dilution to 1 M urea. Peptides were acidified, cleared, desalted on HyperSep C18 cartridges and eluted with 50% acetonitrile 0.1% TFA. A fraction was reserved for total proteomics.

The remaining peptides were processed using the PTMScan HS Phospho Tyrosine kit (Cell Signaling #38572). Peptides were incubated with anti phosphotyrosine beads and bound peptides were eluted and cleaned up on SDB stage tips, then eluted with 60% acetonitrile 0.125% ammonia and dried.

*$TiO_2$ phosphopeptide enrichment*

Supernatants from phosphotyrosine enrichment were desalted on C18 columns, eluted, dried and resuspended in Tris buffer, then enriched for phosphopeptides using $TiO_2$ beads in the EasyPhos format [7]. After binding, washes and elution with ammonia containing acetonitrile, samples were further cleaned on RPS stage tips and eluted in 60% acetonitrile 0.125% ammonia, dried and resuspended for LC-MS.

*Total proteomics*

Aliquots of digested peptides were dried, resuspended in TEAB, quantified and labelled with TMT10plex reagents. After quenching with hydroxylamine, labelled samples were pooled, dried and fractionated into six fractions using a high pH reversed phase kit. Fractions were dried, resuspended in 3.5% acetonitrile 0.1% TFA, quantified and injected at 800 ng per run.

*Mass spectrometry*

Peptides were separated on an Acclaim PepMap RSLC column with a multi-step acetonitrile gradient and analyzed on an Orbitrap Exploris 240 in data dependent acquisition mode. For phosphopeptides, MS1 resolution was 120,000 with AGC $4 \times 10^5$ and maximum injection time 50 ms. MS2 was acquired at 60,000 resolution with HCD 30, AGC $1 \times 10^5$ and maximum injection 110 ms, with dynamic exclusion of 30 seconds.

For TMT runs, MS1 resolution was 120,000 and MS2 resolution 50,000 with HCD 38, AGC $1 \times 10^5$ and maximum injection 105 ms, with 60 second dynamic exclusion.

Raw data were processed in Proteome Discoverer 3.1 with Sequest HT against the UniProt mouse database. Trypsin specificity, two missed cleavages, ten ppm precursor and 0.02 Da fragment tolerances, variable modifications including oxidation, phosphorylation, pyroglutamate and acrylamide, and fixed TMT labelling were used. False discovery rate was controlled at 1% at protein, peptide and site levels by Percolator.

**Co-immunoprecipitation**

$1.2 \times 10^7$ HepG2 cells were transduced with AdV-PTPRF at a multiplicity of infection (MOI) of 25. Cells were lysed in cell lysis buffer and subjected to immunoprecipitation using an anti-PTPRF antibody pre-bound to Dynabeads Protein A. Co-immunoprecipitation of LPP and BCAR1 was assessed by immunoblotting using the corresponding antibodies described in **Supplementary Table 5**.

**Dynamic secretion experiments with isolated islets**

Dynamic perifusion assays were performed on batches of 50–150 isolated islets. Islets were perifused at 37 °C at a flow rate of 0.5 ml $min^{-1}$ using a temperature-controlled perifusion system. Prior to experimental stimulation, islets were equilibrated for 20 min in low-glucose buffer (G1; 1 mM glucose). Effluent fractions were then collected at 4-min intervals throughout the experiment. For **Fig. 5k** and **Supplementary Fig. 5m**, isolated primary mouse islets were sequentially stimulated with low glucose (G1; 1 mM), high glucose (G15; 15 mM), diazoxide

(250 µM), and 30 mM KCl, as indicated in the figure schematics. For **Fig. 5n**, isolated primary human islets were sequentially stimulated with low glucose (G1) and high glucose (G15). Following perifusion, total islet insulin content was measured and used to normalize secretion data.

#### Intracellular $Ca^{2+}$ imaging in isolated mouse islets

Following a 20 min equilibration perifusion in Krebs–Ringer buffer containing 1 mM glucose, isolated mouse islets were sequentially stimulated with low glucose (G1; 1 mM), high glucose (G15; 15 mM), diazoxide, and KCl, as indicated in the figures. Cytosolic $Ca^{2+}$ dynamics were measured in isolated mouse islets loaded with the ratiometric dye fura-2 LR. Islets were perifused in Krebs–Ringer buffer at 37°C in a temperature-controlled chamber placed on the stage of a Zeiss Axiovert 100 microscope equipped with a 20x objective. The Fura-2 LR fluorescence ratio (λex 340/380 nm; λem 510 nm) was acquired every 3s with a prime 95B camera (Teledyne, USA). Fluorescence ratios were recorded in small- to medium-sized islets (diameter ~75–150 µm) and quantified using MetaFluor software (Molecular Devices).

#### CRISPR–Cas12a–mediated PTPRF knockout in hESCs

Guide RNAs flanking exon 23 of the human PTPRF locus were designed using Benchling, CRISPOR, CHOPCHOP and Cas-Designer. H1 human embryonic stem cells (hESCs) were electroporated using the Neon Transfection System as previously described [8]. Briefly, Alt-R Cas12a (Cpf1) Ultra nuclease (Integrated DNA Technologies,#10001272) was complexed with Alt-R CRISPR–Cas12a crRNAs (Integrated DNA Technologies; sequences listed in **Supplementary Table 6**) to generate ribonucleoprotein editing complexes prior to electroporation. Genomic modification of PTPRF was screened by PCR using internal and external primer sets (**Supplementary Table 6**). Single-cell–derived clones were isolated, expanded, and validated by Sanger sequencing (Eurofins Genomics) to confirm targeted editing and preservation of the non-edited allele. Potential off-target sites predicted by CRISPOR were amplified by PCR and sequenced to verify the absence of unintended mutations. Sequencing primer sequences are provided in **Supplementary Table 6**.

#### Stem cell culturing

Generated H1-hESC *PTPRF*$^{-/-}$ and *PTPRF*$^{+/+}$ clones [8] were thawed in mTeSR Plus medium (STEMCELL Technologies Cat#100–0276) supplemented with Y-27632 ROCK pathway inhibitor (STEMCELL Technologies, Cat#72304) and seeded into 3.5-cm dishes. Once cells reached 70–80% confluence, they were detached using 0.5 mM EDTA (Invitrogen, Cat#15575020) and passaged at a 1:6–1:10 ratio into new Matrigel-coated vessels (Corning, Cat#356231).

#### Stem cell differentiation into SC-islets

CRISPR–Cas12a–edited H1 hESC *PTPRF*$^{-/-}$ and *PTPRF*$^{+/+}$ clones were differentiated into stem cell–derived islets (SC-islets) using a previously established seven-stage protocol [8, 9]. Briefly, hESCs were dissociated into single cells using Accutase and seeded at $2 \times 10^6$ cells per well in 6-well plates or $2.5 \times 10^5$ cells per well in 8-well chamber slides. Differentiation through definitive endoderm and pancreatic progenitor stages (stages 1–4) was performed in adherent monolayer culture. At stage 4, cells were dissociated into single-cell suspensions and reaggregated at $1 \times 10^6$ cells per well in 24-well AggreWell plates (STEMCELL Technologies) for three-dimensional differentiation through endocrine induction and maturation (stages 5–7).

For single-cell RNA sequencing, stage-7 SC-islets were dissociated into single cells using trypsin and DNase I, filtered through 40 µm strainers to remove aggregates, and resuspended in PBS containing 0.04% BSA. Cell concentration and viability were assessed using a Luna automated cell counter, and only samples with >85% viability were processed for single-cell library preparation.

**SC-islet implantation in NOD–SCID mice and *in vivo* functional assessment**

Stem cell–derived islets (SC-islets) were implanted under the kidney capsule of immunodeficient NOD–SCID mice to assess *in vivo* maturation and function. Briefly, 3,000 SC-islets (~$2.25 \times 10^6$ cells) derived from *PTPRF*$^{-/-}$ or *PTPRF*$^{+/+}$ hESCs were implanted under the kidney capsule of 8–12-week-old male or female mice and allowed to mature *in vivo* for up to 16 weeks. To assess graft dependence of glucose homeostasis, endogenous murine β cells were ablated after graft maturation by two intraperitoneal injections of streptozotocin (STZ; 150 µg $g^{-1}$ body weight) administered 1 week apart. One week after the second STZ injection, the graft-bearing kidney was surgically removed to induce insulin deficiency. Non-fasted plasma human C-peptide concentrations were measured at week 8 post-implantation. Fasted plasma human C-peptide concentrations were measured at week 12 and 16 post-implantation. Random blood glucose levels were measured at weeks 8, 12, and 16, and then daily following STZ treatment until diabetes onset. All random glycaemia measurements were performed at a standardized time of day (09:00–10:00) to control for diurnal variation.

Intraperitoneal glucose tolerance tests and *in vivo* glucose-stimulated insulin secretion assays were performed at weeks 12 and 16 post-implantation. Mice were fasted for 6 h before intraperitoneal administration of D-glucose (2 g $kg^{-1}$ body weight; Millipore, Cat#108342) in 200 µl DPBS. Blood glucose was measured from tail vein blood using a glucometer (Accu-Chek Performa, Roche). Additional blood samples were collected into EDTA-coated capillary tubes (Sarstedt) for quantification of plasma human C-peptide using an ultrasensitive ELISA (Mercodia, Cat#10-1141-01).

**Immunofluorescence**

H1-hESCs, HLCs and SC-islets were seeded or differentiated in 8-well culture slides (Corning) and fixed with 4% PFA. Cells were permeabilized with 0.5% Triton X-100 in DPBS, followed by incubation with UltraVision Protein Block (Fisher Scientific Cat#15169666) to reduce nonspecific background staining. Primary antibodies (**Supplementary Table 5**) were diluted in 0.1% Tween in DPBS and incubated overnight at 4 °C. After washing with DPBS, secondary antibodies (**Supplementary Table 5**) were diluted in 0.1% Tween in DPBS and incubated for 1 h at room temperature in the dark, followed by washing with DPBS. Slides were mounted with VECTASHIELD Antifade Mounting Medium with DAPI (Vector Laboratories, Cat#VEC.H-1200) and imaged with Zeiss Observer coupled with Colibri 5 Multicolor LED Light Source (Carl Zeiss Microscopy, Gmbh). For image acquisition, ZEN 3.2 software (Carl Zeiss Microscopy, Gmbh) was used.

**RNA extraction and qRT PCR**

Total RNA from tissues and hepatocytes was extracted using the RNeasy Mini Kit, QIAGEN. cDNA was synthesized using Eurogentec reverse transcriptase. qRT PCR was performed on a Bio Rad CFX system with SYBR Green chemistry. Reactions were run in duplicate, and values were normalized to either the highest value or the mean of the control group as specified. Primer sequences are listed in **Supplementary Table 6**.

Poly(A) + mRNA was extracted from H1-hESCs and cells from different stages during HLCs and SC-islets differentiation using Dynabeads mRNA DIRECT kit (Invitrogen, Cat#61012). The differentiation timeline and gene expression analysis were conducted based on an established protocol [8]. The extracted mRNA was reverse transcribed with a reverse transcriptase kit (Eurogentec, Cat#RT-RTCK-03). qPCR was performed with SYBR Green reagent (Bio-Rad Laboratories, Cat#1725274) using a Bio-Rad CFX machine (Bio-Rad Laboratories). β-actin (for H1-hESCs), Cyclophilin G (for SC-islets) were used as the endogenous housekeeping gene to calculate ΔCt values, which were normalized to undifferentiated H1-hESCs to derive ΔΔCt values and relative gene expression. Oligonucleotide sequences for qPCR primers are listed in **Supplementary Table 6**.

**Histology and immunohistochemistry**

Human and mouse livers were rinsed in PBS and fixed in 4% buffered formaldehyde. Tissues were paraffin embedded, sectioned at 7 μm and stained with hematoxylin and eosin.

For PTPRF immunohistochemistry in human liver samples, sections were mounted on charged slides, subjected to citrate-based antigen retrieval, permeabilized with 0.1% Triton X-100, blocked with 2% milk and 10% normal goat serum, and incubated overnight at 4°C with anti-PTPRF antibody (SAB4200321, 1:100). The antibody has been validated by decreased intensity in paraffin-embedded human HepG2 cells with siRNA-mediated PTPRF knockdown compared to controls. Sections were then incubated with HRP-conjugated goat anti-rabbit secondary antibody (P0448, 1:200) for 1 h. Negative controls omitting the primary or secondary antibody were included. Slides were scanned using a NanoZoomer Digital Pathology system and analyzed with CellProfiler automated counting software (Broad Institute, Cambridge, MA, USA). For quantification, five random fields of view were selected from each liver section across the whole tissue area.

**Pancreatic and graft immunofluorescence**

Pancreata were harvested from sacrificed mice, rinsed in saline, and fixed in 4% buffered formaldehyde. Tissues were embedded in paraffin and sectioned at 7 μm thickness using a rotary microtome. Sections were mounted on positively charged slides and processed for immunofluorescence staining. Paraffin sections were deparaffinized and rehydrated, followed by antigen retrieval in heated 10 mM citrate buffer. Sections were permeabilized with 0.1% Triton X-100 and sequentially blocked with 2% milk (15 min) and 10% normal goat serum (30 min, room temperature) to reduce non-specific binding. Sections were incubated overnight at 4 °C with primary antibodies against insulin (Dako Agilent, #A0564) and glucagon (Sigma-Aldrich, #G2654). After washing, sections were incubated for 1 h at room temperature with fluorophore-conjugated secondary antibodies: goat anti–guinea pig IgG for insulin (Thermo Fisher, #A21435 or #A11073) and donkey anti–mouse IgG for glucagon (Sigma-Aldrich, #A21202 or #A32794). Sections incubated with secondary antibodies alone served as negative controls. Nuclei were counterstained with DAPI (Vector Laboratories, #H-1200) before mounting. Images were acquired using an Axio Observer D1 fluorescence microscope (Carl Zeiss). Quantification of insulin- and glucagon-positive cells was performed using CellProfiler, calculating the percentage of hormone-positive cells per islet.

For graft analysis, retrieved SC-islet–containing kidneys were fixed in 4% buffered formaldehyde for 24 h, paraffin-embedded, and sectioned at 7 μm. Immunofluorescence staining was performed as described above using antibodies listed in **Supplementary Table 5**. Graft sections were imaged using a Zeiss Observer microscope equipped with a Colibri 5 multicolor LED light source. Image acquisition was performed using ZEN 3.2 software (Carl Zeiss), and quantification of insulin- and glucagon-positive cells within graft islet-like structures was performed using CellProfiler.

**Pancreatic Insulin Content**

Pancreas pieces close to the intestine and liver were collected followed by homogenization with ethanol/water/acid (75/23.5/1.5, % v/v). Homogenized samples were incubated on a tube rotator for 24 h at 4 °C and centrifuged at 1500 g for 30 min. Supernatant was collected and used for insulin measurement using a commercial Insulin ELISA kit (Mercodia, #10-1247-10). Insulin content was expressed as µg insulin/g pancreas weight (µg/g).

**Bioinformatic analysis**

Complete computational workflows, including R and Python scripts, parameter settings and software versions, are deposited alongside processed data to enable independent replication. Raw mass spectrometry files, single-cell sequencing data, metadata and custom analysis scripts will be deposited in public repositories upon acceptance and are available from the corresponding author upon reasonable request during peer review.

*Preprocessing and quantification*

Global proteome and phosphoproteome datasets were generated for mouse samples (total protein, phosphoserine, phosphothreonine and phosphotyrosine) and human samples (total protein and phosphoserine). Raw intensity tables from MaxQuant were imported into R version 4.1.2. Protein-level matrices were filtered to remove entries with incomplete quantification or ambiguous annotations. Phosphosite-level data were summarized per site without collapsing across sites within proteins, and the technical replicate with the highest intensity was retained for each phosphosite. Entries with ambiguous mapping or poor site localization were excluded. Imputation was applied only to rows with at least 75% observed values using a group-wise minimum approach.

*Normalization and differential analysis*

Data were $\log_2$-transformed and normalized using the voom method in limma version 3.50.1. Principal component analysis was used to assess sample structure and batch effects; where required, unwanted variation was removed with RUVg from RUVSeq, and relevant principal components were included as covariates in downstream models. Differential abundance was assessed using empirical Bayes-moderated $t$-statistics in limma. For proteomics and phosphoproteomics, significance was defined as $P < 0.05$ and absolute $\log_2$ fold change > 0.3785.

*Pathway enrichment and integration*

Over-representation analysis was performed using KEGG and Gene Ontology 2023 gene sets with clusterProfiler and enrichR. Gene set enrichment analysis was applied to ranked human gene lists to test for coordinated pathway-level shifts. Cross-species integration of shared and unique proteins, phosphosites and pathways was visualized using Venn diagrams and heatmaps generated with pheatmap.

*Interactome analysis*

Interactome profiling was performed on affinity purification–mass spectrometry datasets from mouse hepatocytes and HepG2 cells under three conditions: wild-type PTPRF intracellular domain, substrate-trapping CS mutant and beads-only control. Raw files were processed in Proteome Discoverer 2.5 and further analyzed in Perseus. High-confidence interactors were defined as >4-fold enriched relative to beads-only, >2-fold differentially enriched between bait conditions and statistically significant at $P < 0.05$. Gene Ontology enrichment was performed on the resulting interactor lists, and semantically related GO terms were clustered using REVIGO to generate tree maps and reduce redundancy.

*XBP1 ChIP–seq analysis and motif identification*

Publicly available XBP1 ChIP-seq datasets from mouse liver (GSE150889) and human HepG2 cells (ENCODE ENCSR988EVQ) were analyzed to examine regulatory interactions between XBP1 and the *Ptprf/PTPRF* locus. ChIP-seq enrichment peaks across the mouse *Ptprf* locus were identified using an enrichment threshold of >1.5-fold over input. Peaks were scanned for canonical XBP1 binding motifs corresponding to ER stress response elements (ERSE; CCACG/CGTGG). Conservation of regulatory elements was assessed by mapping human ENCODE XBP1 ChIP-seq peaks to the orthologous mouse genomic region (chr4:117,839,323) by syntenic alignment. Non-canonical peaks lacking ERSE motifs were further analyzed for enrichment of transcription factor binding motifs associated with unfolded protein response regulatory complexes, including NF-Y, ATF6, ATF4, C/EBP, CREB and AP-1.

**Single-cell RNA sequencing analysis**

*Human pancreatic single cell RNA-sequencing analysis*

Single-cell RNA sequencing data from human pancreatic islets of nine female donors were obtained from the Human Pancreas Analysis Program (HPAP, https://hpap.pmacs.upenn.edu) and stratified into two metabolic groups: Obese non- and T2D. All samples were uniformly processed with Cell Ranger v10.0.0 (10x Genomics) using the GRCh38 reference genome with intronic read inclusion (--include-introns=true) to maximize sensitivity for lowly expressed genes such as receptor-type protein tyrosine phosphatases. Quality-controlled cells (200–6,000 genes, <20% mitochondrial reads) were integrated using the SCTransform and CCA anchor-based workflow in Seurat v5, with 3,000 shared integration features filtered to genes present across all SCT models. Clustering was performed at resolution 0.1 on 50 principal components, and cell types were annotated based on canonical marker expression (*INS* for β cells, *GCG* for α, *SST* for δ, *KRT19/SOX9* for ductal, *PRSS1/CPA1* for acinar, *COL1A1/ACTA2* for stellate, *PECAM1/VWF* for endothelial, and *PTPRC/CD68* for immune cells). PTPRF expression in β beta cells was quantified using SCTransform-corrected counts, with pairwise Wilcoxon rank-sum tests for group comparisons and pseudobulk donor-level means to account for the nested structure of cells within donors. Donor characteristics are provided in **Supplementary Table 4**.

*Mouse pancreatic islets*

Single-cell gene expression profiles were generated from pancreatic islets of $Ptprf^{\Delta\beta}$ and control mice following HFHFHC diet feeding using the 10x Genomics Chromium Single-Cell 3′ platform. Raw sequencing data were processed using Cell Ranger. Ambient RNA contamination was corrected with SoupX, and doublets were identified and removed using scDblFinder. Quality control retained cells with <10% mitochondrial reads. Data were normalized using SCTransform, and the first 50 principal components were used for UMAP visualization and shared nearest-neighbor clustering at resolution 0.1. Cell types were manually annotated based on canonical markers: β cells (*Ins1*, *Ins2*, *Iapp*, *Mafa*), α cells (*Gcg*) and δ cells (*Sst*, *Hhex*). A small non-endocrine population was identified by the absence of canonical endocrine markers. To characterize β-cell heterogeneity, β cells were subset and reclustered using SCTransform with the first 30 principal components and clustering resolution 0.4, revealing transcriptionally distinct β-cell subpopulations.

*Differential expression, visualization, and enrichment analysis*

Differential expression analysis was performed using FindMarkers [10] (Wilcoxon rank-sum test, min.pct = 0.1). Genes with $\log_2$ fold change > 0.25 and Bonferroni-adjusted $P < 0.05$ were considered significant. Volcano plots were generated with significance thresholds of adjusted $P < 0.05$ and $|\log_2 \text{fold change}| > 1$, displaying and labeling biologically meaningful DEGs using MagmaFlow. Pathway enrichment was performed using GO [11], KEGG [12], and Reactome [13] databases; mouse data additionally used MSigDB [14] and Tabula Muris [15].

For heatmaps, pseudobulk expression was calculated per genotype, $\log_2$-transformed, and gene-centered. Genes were grouped into functional categories (secretion machinery, $Ca^{2+}$/ion handling, cytoskeleton/adhesion, β-cell identity, ribosomal/protein synthesis, mitochondrial OXPHOS, ER stress/proteostasis, secretory stress, immune/oxidative stress) and visualized using ComplexHeatmap [16].

*Human stem cell-derived islets*

Single-cell gene expression profiles were generated from *PTPRF*$^{-/-}$ and *PTPRF*$^{+/+}$ stem cell-derived islets (SC-islets) at stage 7 of differentiation using the 10x Genomics Chromium Single-Cell 3′ platform. Raw sequencing data were processed using Cell Ranger. SoupX [17] correction was applied to reduce ambient RNA contamination, and doublets were identified and removed using scDblFinder [18]. Quality control filtering retained cells expressing 1,000–10,000 genes, <20% mitochondrial reads, and 4–25% ribosomal content. Data from both genotypes were merged and normalized using SCTransform [19] with regression of sequencing depth and mitochondrial percentage. The first 30 principal components were used for UMAP visualization and SNN clustering (resolution = 0.2).

Cell types were assigned using module scores calculated with AddModuleScore for positive and negative markers: β-like cells (positive: *INS, IAPP, MAFA, UCN3, PDX1, NKX6-1, PCSK1, CHGA*; negative: *GCG, SST, PPY, FEV*), α-like cells (*GCG, ARX*), δ-like cells (*SST, HHEX*), ε-like cells (*GHRL*), pancreatic progenitors (*PDX1, NKX6-1, SOX9, HNF1B*), enterochromaffin cells (*TPH1, FEV*), ductal-like cells (*KRT19, KRT7, SOX9, CFTR*), and mesenchymal-like cells (PRSS1). Polyhormonal cells were identified by co-expression of *INS, GCG*, and *SST*.

To characterize β-cell heterogeneity, β-like cells were subset and reclustered using SCTransform [20]. The first 20 principal components were used for UMAP visualization and clustering (resolution = 0.2), revealing transcriptionally distinct β-like cell populations (β-like cells cluster 1 and β-like cells cluster 2).

**Statistical analysis**

Sample size *n* refers to individual mice, independent hepatocyte and islet preparations, individual human liver samples, independent experiments or individual donors as indicated in figure legends. Data are presented as mean ± standard error (SE) of the mean unless stated otherwise. For physiological, metabolic and biochemical endpoints, comparisons between two groups were performed using two-sided Student's *t*-tests. Comparisons among three or more groups used one-way, two-way or repeated-measures analysis of variance followed by appropriate post-hoc tests. Analyses were performed in GraphPad Prism. Sample sizes were chosen based on variability from previous experiments and pilot data. For global proteomics and phosphoproteomics, significance was defined as $P < 0.05$ with empirical Bayes moderation. For single-cell and single-nucleus RNA sequencing, differential expression $P$ values were adjusted using Bonferroni correction, and pathway enrichment analyses were corrected by the Benjamini–Hochberg false discovery rate method. Exact $P$ values are reported in figure panels and source data. Statistical tests, multiple-comparison procedures and software versions are specified in figure legends.

### Supplementary Tables:

| ID | Gender | Age (y) | Weight (kg) | BMI (kg/m2) | Glucose (mg/dl) | Insulin (U/L) | HOMA -IR | TG (mg/dl) | Diagnosis | Steatosis score |
|---|---|---|---|---|---|---|---|---|---|---|
| EG28 | M | 52 | 132.5 | 41.8 | 83 | 23 | 4.71 | 98 | Healthy | 0 |
| EG48 | F | 20 | 135.2 | 50.9 | 88 | 44.5 | 9.63 | 101 | Healthy | 0 |
| OM-91 | F | 30 | 120.2 | 44.7 | 104 | 8.2 | 2.12 | 121 | Healthy | 0 |
| OM-88 | F | 31 | 119.8 | 41.5 | 86 | 11.7 | 2.5 | 181 | Healthy | 0 |
| EG29 | F | 32 | 122.4 | 43.4 | 96 | 25.9 | 6.16 | 206 | Healthy | 0 |
| EG01 | F | 35 | 116.4 | 48.4 | 63 | 9.6 | 1.5 | 87 | Healthy | 0 |
| EG12 | F | 68 | 106.8 | 36.9 | 106 | 6 | 1.57 | 154 | Healthy | 0 |
| EG31 | F | 31 | 119.3 | 48.9 | 102 | 43.1 | 10.85 | 183 | Healthy | 0 |
| OM-66 | F | 40 | 117.4 | 39.7 | 94 | 1.4 | 2.12 | 68 | Healthy | 0 |
| OM-67 | F | 40 | 113.2 | 44.2 | 90 | 11.4 | 2.53 | 104 | Healthy | 0 |
| EG46 | F | 41 | 116.7 | 45 | 81 | 12.6 | 2.53 | 170 | Healthy | 0 |
| EG33 | F | 42 | 104.7 | 41.4 | 94 | 4.3 | 0.98 | 143 | Healthy | 0 |
| EG20 | F | 44 | 114.1 | 46.3 | 89 | 10 | 2.19 | 146 | Healthy | 0 |
| OM-76 | F | 45 | 128.8 | 45.6 | 104 | 9.4 | 2.4 | 124 | Healthy | 0 |
| EG45 | F | 52 | 101.1 | 40.5 | 85 | 4.3 | 0.91 | 112 | Healthy | 0 |
| OM-71 | F | 54 | 108.7 | 37.6 | 102 | 4.9 | 1.25 | 106 | Healthy | 0 |
| EG30 | F | 55 | 121.5 | 42 | 97 | 6.2 | 1.4 | / | Healthy | 0 |
| OM-86 | F | 55 | 118.1 | 40.9 | 93 | 6.8 | 1.56 | 166 | Healthy | 0 |
| OM-78 | F | 61 | 98.6 | 43.8 | 85 | 42.1 | 8.84 | 213 | Healthy | 0 |
| OM-73 | M | 30 | 173.1 | 49.5 | 87 | 21.3 | 4.58 | 229 | Steatosis | 1 |
| EG04 | M | 43 | 121.1 | 39.1 | 101 | 11.9 | 2.98 | 170 | Steatosis | 1 |
| EG40 | M | 44 | 145 | 52 | 128 | 31.3 | 9.86 | 153 | Steatosis | 1 |
| EG13 | M | 52 | 149.1 | 43.6 | 107 | 11.9 | 3.14 | 126 | Steatosis | 1 |
| OM-80 | M | 54 | 117.8 | 33.7 | 87 | 14.8 | 3.16 | 279 | Steatosis | 1 |
| EG16 | M | 56 | 142.4 | 52.7 | 84 | 34 | 7.08 | 101 | Steatosis | 1 |
| EG26 | M | 58 | 117.4 | 42.6 | 85 | 6 | 1.26 | 101 | Steatosis | 1 |
| EG19 | M | 64 | 97.3 | 36.6 | 101 | 31.2 | 7.77 | 246 | Steatosis | 1 |
| EG49 | F | 43 | 117.3 | 52.1 | 99 | 12.5 | 3.05 | 130 | Steatosis | 1 |
| EG32 | F | 24 | 116.2 | 41.2 | 82 | 11.5 | 2.33 | 519 | Steatosis | 1 |
| OM-92 | F | 38 | 89.8 | 43.3 | 84 | 16.7 | 3.47 | 172 | Steatosis | 1 |
| EG44 | F | 46 | 101.1 | 39.8 | 58 | 4.1 | 0.59 | 101 | Steatosis | 1 |
| OM-89 | F | 49 | 105.1 | 41.1 | 100 | 17.6 | 4.32 | 210 | Steatosis | 1 |
| EG35 | F | 51 | 134.1 | 46.4 | 117 | 9 | 2.6 | 163 | Steatosis | 1 |
| EG43 | F | 51 | 109.4 | 44.4 | 84 | 9 | 1.86 | 155 | Steatosis | 1 |
| OM-72 | F | 51 | 139 | 51.1 | 97 | 18.5 | 4.43 | 153 | Steatosis | 1 |
| OM-74 | F | 51 | 79.1 | 32.9 | 92 | 11.3 | 2.57 | 255 | Steatosis | 1 |
| EG02 | F | 55 | 84.2 | 40.6 | 90 | 16.1 | 3.58 | 565 | Steatosis | 1 |
| EG36 | F | 58 | 94 | 39.1 | 148 | 13.3 | 4.88 | 162 | Steatosis | 1 |
| OM-85 | F | 47 | 99.5 | 35.7 | 152 | 15.3 | 5.76 | 204 | Steatosis | 1 |
| EG07 | F | 45 | 135.5 | 52.9 | 94 | 9.2 | 2.13 | 89 | MASH | 1 |
| EG05 | F | 51 | 75.5 | 35.4 | 105 | 7.8 | 2.03 | 131 | MASH | 1 |
| EG09 | F | 54 | 91.8 | 37.7 | 166 | 31.8 | 13.03 | 174 | MASH | 1 |
| EG15 | F | 54 | 112 | 44.3 | 93 | 58.6 | 13.46 | 129 | MASH | 1 |
| OM-75 | F | 42 | 152 | 53.2 | 327 | 0.4 | 0.32 | 106 | MASH | 1 |
| OM-83 | M | 27 | 127 | 45 | 88 | 25.4 | 5.52 | 183 | Steatosis | 2 |
| OM-77 | M | 61 | 112.4 | 34.7 | 174 | 10.7 | 4.6 | 358 | Steatosis | 2 |
| EG37 | F | 34 | 113.9 | 38 | 90 | 17 | 3.79 | 204 | Steatosis | 2 |
| EG34 | F | 62 | 108 | 37.8 | 109 | 35.5 | 9.55 | 230 | Steatosis | 2 |
| OM-69 | M | 33 | 141.5 | 48.4 | 227 | 35.1 | 19.68 | 421 | MASH | 2 |
| OM-65 | M | 41 | 163.1 | 49.2 | 83 | 26.5 | 5.4 | 169 | MASH | 2 |
| EG06 | F | 27 | 115.4 | 45.1 | 103 | 16 | 4.07 | 67 | MASH | 2 |
| EG17 | F | 40 | 122.1 | 47.7 | 79 | 11 | 2.14 | 309 | MASH | 2 |
| EG41 | M | 55 | 137.6 | 51.8 | 248 | 22 | 13.47 | 164 | Steatosis | 3 |
| EG39 | F | 21 | 119.5 | 43.9 | 81 | 17.3 | 3.44 | 311 | Steatosis | 3 |
| EG03 | F | 50 | 113.3 | 40.6 | 106 | 10.6 | 3.14 | 114 | Steatosis | 3 |
| EG42 | F | 53 | 99.5 | 41.4 | 103 | 19.3 | 4.89 | 171 | Steatosis | 3 |
| OM-70 | F | 53 | 117.5 | 50.9 | 82 | 19.2 | 3.89 | 224 | Steatosis | 3 |
| EG38 | F | 55 | 120 | 44.6 | 96 | 14.3 | 3.37 | 290 | Steatosis | 3 |
| OM-79 | F | 70 | 114.8 | 43.2 | 145 | 6 | 2.17 | 212 | Steatosis | 3 |
| EG21 | M | 55 | 133 | 41 | 94 | 16.4 | 3.8 | 264 | MASH | 3 |
| EG18 | M | 61 | 124.4 | 39.2 | 151 | 20.6 | 7.68 | 256 | MASH | 3 |
| EG27 | F | 44 | 112.4 | 43.9 | 123 | 18.2 | 5.55 | 230 | MASH | 3 |
| EG25 | F | 47 | 147.1 | 53.4 | 95 | 12.7 | 2.98 | 243 | MASH | 3 |
| EG22 | F | 52 | 103.4 | 40.4 | 206 | 6.2 | 3.13 | 161 | MASH | 3 |
| EG10 | F | 57 | 94.9 | 38.5 | 113 | 9.4 | 2.64 | 194 | MASH | 3 |
| EG23 | F | 64 | 93.9 | 37.6 | 116 | 14.7 | 4.2 | 167 | MASH | 3 |

**Supplementary Table 1:** Human liver donor characteristics for immunohistochemistry.

| ID | Gender | Age (y) | Weight (kg) | BMI (kg/m$^2$) | Glucose (mg/dl) | Insulin (U/L) | HOMA-IR | TG (mg/dl) | Diagnosis |
|---|---|---|---|---|---|---|---|---|---|
| OM-062 | F | 49 | 120 | 43 | 77 | 5.7 | 1.08 | 116 | Healthy |
| OM-049 | F | 52 | 101.1 | 40.5 | 85 | 4.3 | 0.91 | 112 | Healthy |
| OM-071 | F | 54 | 108.7 | 37.6 | 102 | 4.9 | 1.25 | 106 | Healthy |
| OM-053 | F | 43 | 117.3 | 52.1 | 99 | 12.5 | 3.05 | 130 | Steatosis |
| OM-042 | F | 55 | 120 | 44.6 | 96 | 14.3 | 3.37 | 290 | Steatosis |
| OM-046 | F | 53 | 99.5 | 41.4 | 103 | 19.3 | 4.89 | 171 | Steatosis |
| OM-011 | F | 57 | 94.9 | 38.5 | 113 | / | / | 194 | MASH |
| OM-027 | F | 47 | 147.1 | 53.4 | 95 | 12.7 | 2.98 | 243 | MASH |
| OM-025 | F | 64 | 93.9 | 37.6 | 116 | 14.7 | 4.2 | 167 | MASH |

**Supplementary Table 2:** Human liver donor characteristics for mass spectrometry analysis.

| Parameter | HIRP-18 | HIRP-19 | HIRP-23 | HIRP-24 | HIRP-27 | HIRP-31 |
|---|---|---|---|---|---|---|
| Age (years) | 76 | 46 | 70 | 51 | 64 | 64 |
| Sex | F | F | M | F | F | M |
| BMI (kg/m$^2$) | 27 | 31 | 25 | 26 | 20 | 18 |
| HbA1c (%) | 5.7 | 5.4 | 5.2 | 4.4 | 6.3 | 5.7 |
| Glycemia (mg/dL) | 103 | 102 | 118 | 126 | 144 | 146 |
| Diabetes | No | No | No | No | No | No |
| IEQ (day 1) | 20,800 | 148,680 | 191,400 | 118,664 | 266,980 | 69,600 |
| IEQ/g pancreas | 353 | 1,549 | 2,696 | 2,373 | 5,037 | 1,481 |
| Purity (%) | 55 | 75 | 87 | 65 | 76 | 69 |

**Supplementary Table 3:** Characteristics of human islet preparations used for functional studies.

| Group | Donor ID | Age (years) | BMI (kg/m²) | HbA1c (%) | β-cells (n) | PTPRF+ cells (%) | Mean expression |
|---|---|---|---|---|---|---|---|
| **Obese** | HPAP-054 | 40 | 30.9 | 4.8 | 365 | 16.2% | 0.1203 |
| | HPAP-063 | 45 | 38.4 | 6.3 | 304 | 22.4% | 0.1802 |
| | HPAP-101 | 55 | 38.0 | 5.0 | 3,261 | 15.5% | 0.1141 |
| | HPAP-103 | 48 | 36.4 | 6.0 | 1,325 | 8.2% | 0.0583 |
| | HPAP-118 | 51 | 33.0 | 5.7 | 2,283 | 13.7% | 0.1059 |
| | **Group mean** | **48** | **35.3** | **5.6** | **7,538** | **14.0%** | **0.1060** |
| **T2D** | HPAP-058 | 34 | 29.3 | 9.4 | 134 | 31.3% | 0.2427 |
| | HPAP-085 | 48 | 39.8 | 7.3 | 424 | 22.2% | 0.1837 |
| | HPAP-096 | 56 | 38.7 | 7.7 | 441 | 25.9% | 0.2125 |
| | HPAP-109 | 59 | 29.5 | 7.5 | 863 | 24.3% | 0.2031 |
| | **Group mean** | **49** | **34.3** | **8.0** | **1,862** | **24.7%** | **0.2072** |

**Supplementary Table 4.** HPAP Dataset: Female donor characteristics and PTPRF expression in β cells (obese and T2D groups).

| No. | Antibody | Supplier | Dilution |
|---|---|---|---|
| 1 | PTPRF/LAR (SAB4200321) | Sigma-Aldrich | 1:100 (IHC) |
| 2 | PTPRF/LAR (17164) | Cell Signaling Technology | 1:500 (WB)<br>1:200 (IP) |
| 3 | pIR (Tyr1150/1151) (3024) | Cell Signaling Technology | 1:500 (WB) |
| 4 | IR (3025) | Cell Signaling Technology | 1:1000 (WB) |
| 5 | pAKT (Ser473) (4060) | Cell Signaling Technology | 1:1000 (WB) |
| 6 | AKT (2920) | Cell Signaling Technology | 1:1000 (WB) |
| 7 | β-actin (A1978) | Sigma-Aldrich | 1:5000 (WB) |
| 8 | Tubulin (T5168) | Sigma-Aldrich | 1:5000 (WB) |
| 9 | GAPDH | Trevigen | 1:5000 (WB) |
| 10 | PPARγ (2443) | Cell Signaling Technology | 1:500 (WB) |
| 11 | pSTAT3 (Tyr705) (9131) | Cell Signaling Technology | 1:500 (WB) |
| 12 | STAT3 (9139) | Cell Signaling Technology | 1:1000 (WB) |
| 13 | pSTAT1 (Tyr701) (560310) | BD Biosciences | 1:1000 (WB) |
| 14 | STAT1 (14994) | Cell Signaling Technology | 1:1000 (WB) |
| 15 | XBP1s (40435) | Cell Signaling Technology | 1:500 (WB) |
| 16 | CHOP (2895) | Cell Signaling Technology | 1:500 (WB) |
| 17 | pTyr 1000 (8954) | Cell Signaling Technology | 1:1000 (WB) |
| 18 | LPP (mouse monoclonal, 8B3A11) | Cell Signaling Technology | 1:1000 (WB) |
| 19 | BCAR1 (p130 Cas, rabbit monoclonal, E1L9H) | Cell Signaling Technology | 1:1000 (WB) |
| 20 | Goat anti-rabbit IgG (P0448) | Dako Agilent Technologies | 1:6000 (WB) |
| 21 | Goat anti-mouse IgG (P0447) | Dako Agilent Technologies | 1:6000 (WB) |
| 22 | Rabbit PTPRF/LAR (E6W4X) monoclonal | Cell Signaling Technology | 1:500 (WB/IF) |
| 23 | Goat anti-human SOX17 polyclonal | R&D Systems | 1:500 (IF) |
| 24 | Mouse anti-NKX6.1 (R11-560) monoclonal | BD Biosciences | 1:400 (IF) |
| 25 | Rabbit PDX1 (D59H3) XP monoclonal | Cell Signaling Technology | 1:400 (IF) |
| 26 | Guinea pig insulin polyclonal (FLEX RTU) | Agilent Technologies | Ready-to-use (IF) |
| 27 | Rabbit glucagon polyclonal | Cell Signaling Technology | 1:500 (IF) |
| 28 | Rat somatostatin (human/mouse) monoclonal | Bio-Techne | 1:500 (IF) |
| 29 | Donkey anti-rabbit IgG (H+L), Alexa Fluor 488 | Thermo Fisher Scientific | 1:500 (IF) |
| 30 | Donkey anti-rabbit IgG (H+L), Alexa Fluor Plus 555 | Thermo Fisher Scientific | 1:500 (IF) |
| 31 | Donkey anti-mouse IgG (H+L), Alexa Fluor Plus 555 | Thermo Fisher Scientific | 1:500 (IF) |
| 32 | Donkey anti-mouse IgG (H+L), Alexa Fluor 488 | Thermo Fisher Scientific | 1:500 (IF) |
| 33 | Donkey anti-goat IgG (H+L), Alexa Fluor Plus 555 | Thermo Fisher Scientific | 1:500 (IF) |
| 34 | Goat anti-guinea pig IgG (H+L), Alexa Fluor 488 | Thermo Fisher Scientific | 1:500 (IF) |
| 35 | Donkey anti-rat IgG (H+L), Alexa Fluor 647 | Thermo Fisher Scientific | 1:500 (IF) |
| 36 | β-actin (rabbit polyclonal) (4967) | Cell Signaling Technology | 1:5000 (WB) |
| 37 | Pan-actin (mouse polyclonal, AAN02 ) | Cytoskeleton Inc | 1:1000 (WB) |

**Supplementary Table 5:** Primary and secondary antibodies used in this study. IHC: immunohistochemistry, WB: Western blotting, IF: immunofluorescence, IP: immunoprecipitation.

| REAGENT or RESOURCE | SOURCE | IDENTIFIER |
|---|---|---|
| **Single guide RNAs** | | |
| PTPRF guide RNA G1 intron 22-23: Alt-R A.s. Cas12a crRNA | Integrated DNA Technologies | TCTGCCACACTGCTCAAGCCTCA |
| PTPRF guide RNA G2 intron 23-24: Alt-R A.s. Cas12a crRNA | Integrated DNA Technologies | AGGAGCACAGAGAGGAGGGTTGG |
| **Oligonucleotides** | | |
| hPTPRF Forward (Fw) primer P1 | Eurogentec | GTCTCCCATGGTGACTGCCA |
| PTPRF Reverse (Rv) primer P2 | Eurogentec | TGAGCACGTGTCTGAGTGTCC |
| PTPRF Fw primer P3 | Eurogentec | GCCTGTGTATGCCCTACCTC |
| PTPRF Rv primer P4 | Eurogentec | CCTCCTCTCTGTGCTCCTGAA |
| PTPRF sequencing (Seq) Fw primer | Eurogentec | GAGGAAGGAAGCCTCTCTTCCATC |
| PTPRF Seq Rv primer | Eurogentec | GGTGTCACTCACAGCCTTTTG |
| PTPRF standard qPCR Fw primer | Eurogentec | GGAAAAGGACCCACTCTCCG |
| PTPRF standard qPCR Rv primer | Eurogentec | GTTCCCACACCATCCTCCAG |
| PTPRF qPCR Fw primer | Eurogentec | CGATGGCCTCAAGTTCTCCC |
| PTPRF qPCR Rv primer | Eurogentec | TCTTGGGCTTGTTCACCTCC |
| GRAMD1A Fw primer | Eurogentec | CTCAGGTGAACAAGTGGGCT |
| GRAMD1A Rv primer | Eurogentec | CTCACAAGCACTACGCCTGA |
| GRAMD1A Seq Fw primer | Eurogentec | AAATCAGCTAGGCATGGTGG |
| U8-AC093523.1 Fw primer | Eurogentec | CACAAAACTCCCATCTTGCCAT |
| U8-AC093523.1 Rv primer | Eurogentec | ACTCACCTGCATACATAACTAGCTC |
| U8-AC093523.1 Seq Fw primer | Eurogentec | TCATTCCCTCATATGCTTCTTCTG |
| RP11-88H9.2-RP5-965F6.2 Fw primer | Eurogentec | GCCTTCTTAACCTGGGGGAG |
| RP11-88H9.2-RP5-965F6.2 Rv primer | Eurogentec | TGCCTCCCTGCTTATCACAC |
| RP11-88H9.2-RP5-965F6.2 Seq Rv primer | Eurogentec | GAATTTCAGTCAGGTTCCAGG |
| HTR1D-LINC01355 Fw primer | Eurogentec | TTCATGGGCTCCATCAACCC |
| HTR1D-LINC01355 Rv primer | Eurogentec | GTGCACAAAATATGAGGCAATTTTT |
| HTR1D-LINC01355 Seq Rv primer | Eurogentec | TTTAAAGATGAGGCAGGTCAGGC |
| L3MBTL4 Fw primer | Eurogentec | GCCAGTGCTTCTGCTCATTC |
| L3MBTL4 Rv primer | Eurogentec | AGCAGTTCCCTCCAAAACCC |
| L3MBTL4 / Seq Fw primer | Eurogentec | GGGCAGTCTCCTGTGTGTTTGAAG |
| PPAP2A (PLPP1) Fw primer | Eurogentec | AGACCAGGCTTCAGGACACA |
| PPAP2A (PLPP1) Rv primer | Eurogentec | ACACAGACAGGTGAGCGAAA |
| PPAP2A (PLPP1) Seq Rv primer | Eurogentec | ACAAACCACAATAGCTAGTGAGAC |
| ACTB qPCR Fw primer | Eurogentec | CTGTACGCCAACACAGTGCT |
| ACTB qPCR Rv primer | Eurogentec | GCTCAGGAGGAGCAATGATC |
| NGN3 qPCR Fw primer | Eurogentec | GACGACGCGAAGCTCACCAA |

| | | |
|---|---|---|
| NGN3 qPCR Rv primer | Eurogentec | TACAAGCTGTGGTCCGCTAT |
| NKX6.1 qPCR Fw primer | Eurogentec | GGCCTGTACCCCTCATCAA |
| NKX6.1 qPCR Rv primer | Eurogentec | CTCTCTGTCATCCCCAACGA |
| PDX1 qPCR Fw primer | Eurogentec | GCTGCCTTTCCCATGGATGA |
| PDX1 qPCR Rv primer | Eurogentec | CAACATGACAGCCAGCTCCA |
| INS qPCR Fw primer | Eurogentec | TCTACCTAGTGTGCGGGGAA |
| INS qPCR Rv primer | Eurogentec | CTCCACCTGCCCCACCT |
| GCG qPCR Fw primer | Eurogentec | AACATTGCCAAACGTCACGA |
| GCG qPCR Rv primer | Eurogentec | GGGAAATCTCGCCTTCCTCG |
| Ptprf (ex8-10) standard qPCR Fw primer | Eurogentec | GGCTCCTCGTTTCTCC ATCC |
| Ptprf (ex8-10) standard qPCR Rv primer | Eurogentec | GTGGTAAGGAGTCC TGCGTC |
| mPtprf (ex8-10) qPCR Fw primer | Eurogentec | CATGATAGAAGCCA CGGCCC |
| Ptprf (ex8-10) qPCR Rv primer | Eurogentec | AGGCCACCAATGCTG TATCG |
| Ptprf (ex23) standard qPCR Fw primer | Eurogentec | GGCTCAAGTTCTCCC AGGA |
| Ptprf (ex23) standard qPCR Rv primer | Eurogentec | ACTCCACAGTGTCC ACCAG |
| Ptprf (ex23) qPCR Fw primer | Eurogentec | ACAGCAGTTCACAT GGGAGA |
| Ptprf (ex23) qPCR Rv primer | Eurogentec | ATCCTCCAGAAATCGC CCAT |
| Pparα standard qPCR Fw primer | Eurogentec | GCATGTGAAGGCTGT AAGGGC |
| Pparα standard qPCR Rv primer | Eurogentec | GACAAAAGGCGGGT TGTTGCT |
| Pparα qPCR Fw primer | Eurogentec | GCTGTAAGGGCTTC TTTCGG |
| Pparα qPCR Rv primer | Eurogentec | GCGAATTGCATTGTGTC GACAT |
| Pparγ1 standard qPCR Fw primer | Eurogentec | GCTCCAAGAATACCAA AGTGCG |
| Pparγ1 standard qPCR Rv primer | Eurogentec | AACCTGATGGCATT GTGAGACA |
| Pparγ1 qPCR Fw primer | Eurogentec | CCAAGAATACCAAA GTGCGATCA |
| Pparγ1 qPCR Rv primer | Eurogentec | AAAACCCTTGCATCCT TCACAA |
| Pparγ2 standard qPCR Fw primer | Eurogentec | AGCATGGTGCCTTCG CTGAT |
| Pparγ2 standard qPCR Rv primer | Eurogentec | GCCCAAACCTGATG GCATTGTG |
| Pparγ2 qPCR Fw primer | Eurogentec | TGCCTATGAGCACTT CACAAG |
| Pparγ2 qPCR Rv primer | Eurogentec | TCTACTTTGATCGCAC TTTGGTA |
| Acaca (Acc) standard qPCR Fw primer | Eurogentec | GCGCTTACATTGTGGA TGGC |
| Acaca (Acc) standard qPCR Rv primer | Eurogentec | AAGCCTTCACTGTG CCTTCA |
| Acaca (Acc) qPCR Fw primer | Eurogentec | AGCCAGAAGGGACA GTAGAA |
| Acaca (Acc) qPCR Rv primer | Eurogentec | CTCAGCCAAGCGGAT GTAAA |
| Fasn standard qPCR Fw primer | Eurogentec | ACTTCCTCTGGGATGT GCCT |
| Fasn standard qPCR Rv primer | Eurogentec | GTCAGCACTGCTCT CGTTGA |

| | | |
|---|---|---|
| Fasn qPCR Fw primer | Eurogentec | CACAGTGCTCAAAG GACATGCC |
| Fasn qPCR Rv primer | Eurogentec | CACCAGGTGTAGTGC CTTCTC |
| Acly standard qPCR Fw primer | Eurogentec | ACACCATCATCTGTGC TCGG |
| Acly standard qPCR Rv primer | Eurogentec | ATCCCAGGGGTGAC GATACA |
| Acly qPCR Fw primer | Eurogentec | TTCGTCAAACAGCA CTTCC |
| Acly qPCR Rv primer | Eurogentec | ATTTGGCTTCTTGGAG GTG |
| Cpt1α standard qPCR Fw primer | Eurogentec | CCACACAAACGGCAG AGCA |
| Cpt1α standard qPCR Rv primer | Eurogentec | TCAGGAGCAACACC TATTCATTTG |
| Cpt1α qPCR Fw primer | Eurogentec | GCTGATGACGGCTA TGGTGT |
| Cpt1α qPCR Rv primer | Eurogentec | AAAGCGGTGTGAGTC TGTCT |
| Nrf2 standard qPCR Fw primer | Eurogentec | ACTACAGTCCCAGCA GAGTGAT |
| Nrf2 standard qPCR Rv primer | Eurogentec | AGACACTGCACTGC AACAAG |
| Nrf2 qPCR Fw primer | Eurogentec | ACTACAGTCCCAGC AGAGTGAT |
| Nrf2 qPCR Rv primer | Eurogentec | TCACACACTTTCTGCG TGCT |
| Actb standard qPCR Fw primer | Eurogentec | AGAGGAAATCGTGCG TGAC |
| Actb standard qPCR Rv primer | Eurogentec | TCTCCTTCTGCATCC TGTCA |
| Actb qPCR Fw primer | Eurogentec | ACGGCCAAGGTCAT CACTATT |
| Actb qPCR Rv primer | Eurogentec | GTTGGCATAGAGGTCT TTACG |
| Gapdh standard qPCR Fw primer | Eurogentec | ATGACTCTACCCACGG CAAG |
| Gapdh standard qPCR Rv primer | Eurogentec | TGTGAGGGAGATGC TCAGTG |
| Gapdh qPCR Fw primer | Eurogentec | AGTTCAACGGCACA GTCAAG |
| Gapdh qPCR Rv primer | Eurogentec | TACTCAGCACCAGCAT CACC |

**Supplementary Table 6.** Single guide RNAs and primers.
